# Comprehensive Structure and Functional Adaptations of the Yeast Nuclear Pore Complex

**DOI:** 10.1101/2021.10.29.466335

**Authors:** Christopher W. Akey, Digvijay Singh, Christna Ouch, Ignacia Echeverria, Ilona Nudelman, Joseph M Varberg, Zulin Yu, Fei Fang, Yi Shi, Junjie Wang, Daniel Salzberg, Kangkang Song, Chen Xu, James C. Gumbart, Sergey Suslov, Jay Unruh, Sue L Jaspersen, Brian T. Chait, Andrej Sali, Javier Fernandez-Martinez, Steven J. Ludtke, Elizabeth Villa, Michael P. Rout

**Affiliations:** Department of Physiology and Biophysics, Boston University School of Medicine, 700 Albany Street, Boston, Massachusetts 02118, USA; Section of Molecular Biology, Division of Biological Sciences, University of California San Diego, La Jolla, CA, 92093, USA; Department of Biochemistry and Molecular Pharmacology, University of Massachusetts Medical School, 364 Plantation St, Worcester, MA 01605; Department of Bioengineering and Therapeutic Sciences, University of California, San Francisco, San Francisco, CA, USA; Laboratory of Cellular and Structural Biology, The Rockefeller University, New York, NY, 10065, USA; Department of Cell Biology, University of Pittsburgh, Pittsburgh, PA, USA; Laboratory of Mass Spectrometry and Gaseous Ion Chemistry, The Rockefeller University, New York, NY, USA; School of Physics, Georgia Institute of Technology, Atlanta, GA 30332, USA; Stowers Institute for Medical Research, Kansas City, MO, USA; Department of Molecular and Integrative Physiology, University of Kansas Medical Center, Kansas City, KS, USA; Verna and Marrs McLean Department of Biochemistry and Molecular Biology, Baylor College of Medicine, 1 Baylor Plaza, Houston, Texas 77030, USA; Department of Cellular and Molecular Pharmacology, San Francisco, San Francisco, CA 94158, USA; Quantitative Biosciences Institute, University of California San Francisco, San Francisco, CA 94158, USA; Department of Pharmaceutical Chemistry, University of California San Francisco, San Francisco, CA 94158, USA; Howard Hughes Medical Institute, University of California San Diego

**Author notes:** These authors contributed equally (Christopher W. Akey, Digvijay Singh, Christna Ouch, Ignacia Echeverria).

**Keywords:** Nuclear pore complex, nucleocytoplasmic transport, cryo-electron microscopy, cryo-electron tomography

## Abstract

Nuclear Pore Complexes (NPCs) mediate the nucleocytoplasmic transport of macromolecules. Here we provide a structure of the yeast NPC in which the inner ring is resolved by cryo-EM at - helical resolution to show how flexible connectors tie together different structural and functional layers in the spoke. These connectors are targets for phosphorylation and regulated disassembly in cells with an open mitosis. Moreover, some nucleoporin pairs and karyopherins have similar interaction motifs, which suggests an evolutionary and mechanistic link between assembly and transport. We also provide evidence for three major NPC variants that foreshadow functional specializations at the nuclear periphery. Cryo-electron tomography extended these studies to provide a comprehensive model of the in situ NPC with a radially-expanded inner ring. Our model reveals novel features of the central transporter and nuclear basket, suggests a role for the lumenal ring in restricting dilation and highlights the structural plasticity required for transport by the NPC.

## Introduction

Nuclear pore complexes (NPCs) are cylindrical assemblies with 8-fold symmetry that form a gateway in the nuclear envelope (NE) for the highly selective exchange of macromolecules between the cytoplasm and nucleus. These multifaceted transport machines stabilize a pore in the NE, termed the pore membrane, and are key players in RNA processing and chromatin organization (1–4). Given this central role, it is not surprising that defects in NPC components are linked to many diseases and that nuclear transport is a valid target for therapeutics (5, 6). NPCs from the yeast Saccharomyces are smaller (52 MDa) (7) than their vertebrate cousins (∼ 109 MDa) (8); however, both scaffolds are composed of about 30 conserved proteins termed Nucleoporins (Nups). Depending upon the Nup and species, these proteins may be present in 8, 16, 32 or 48 copies per NPC (7, 8)

In the yeast NPC, ∼ 550 Nups are arranged in several coaxial rings: an inner ring (IR) with 8 massive subunits (termed spokes) is divided into two nearly perfect half-spokes, which are flanked by cytoplasmic and nuclear outer rings to form the core scaffold. A lumenal ring is anchored at the midline of the membrane pore and is formed by Pom152 (7). Finally, a cylindrical basket extends from the nuclear side and in concert with a cytoplasmic platform comprised of the Nup82 complex, forms a pipeline for RNA processing and export (9, 10). The cylindrical scaffold provides anchor points for an array of Nups with intrinsically disordered Phe-Gly (FG) repeat domains that project inwards to form the central transport path (7, 11); these FG domains provide binding sites for the bi-directional, facilitated diffusion of nuclear transport factors with their macromolecular cargos (12–14)

To better understand NPC function, positional information for yeast Nups was determined by integrative modeling with biochemical and physical restraints to create a “nearestneighbor” map (15). In subsequent work, human (16–18) and algal (19). NPCs were studied with cryo-electron tomography (cryo-ET) and sub-tomogram averaging to provide 3D maps at 23-30 Å resolution. Molecular models for human and algal NPCs were then determined by computational docking of ∼ 25 Nup domain crystal structures and homology models (17, 19, 20). In addition, cryo-electron microscopy (cryo-EM) of Xenopus oocyte NPCs and single particle analysis have revealed domains of the cytoplasmic, double outer ring at α-helical resolution and provided details of the lumenal ring (21, 22). For the yeast NPC, cryo-ET and integrative structure determination were used to generate a model of the purified assembly (7) that in turn, was used to help model the in situ complex (23).

When taken together, the data have revealed a remarkable structural redundancy and plasticity that underlies construction of the NPC (7). For example, a lateral offset is present between Nup pairs or paralogs in each half-spoke, that places similar Nups in non-equivalent environments (7, 15, 17, 20, 24). Structural redundancies are also present elsewhere in the NPC (9, 16, 19), as exemplified by multi-tasking Nups that reside in different sub-complexes (16, 18, 25–27). In addition, the number and composition of outer rings can vary between species (7, 16, 19). In human NPCs, large variations in diameter occur in situ (28) and cell cycle dependent changes in the functional size of the transport channel have been observed (29). More recently, 3D studies of NPCs with cryo-ET of focused ion beam (FIB) milled cryo-sections from C. reinhardtii and S. cerevisiae have revealed large-scale changes in the scaffold that involve a radial expansion of the inner ring (19, 23).

Even with these steps forward, our understanding of NPC structure and its role in transport remain far from complete. To address this issue, we applied a multi-pronged strategy to build a comprehensive model of the NPC. We determined a 3D structure of isolated yeast NPCs with cryo-EM and single particle analysis to resolve the inner ring in a radially contracted conformation at α-helical resolution. Structure-based modeling with this map was augmented by a novel, integrative threading method to reveal Nic96 connectors in unprecedented detail within the spoke and the resulting model was validated with crosslinking data. In addition, a complete model was built for a novel, double outer ring at 11 Å resolution. To extend this work, we used cryo-ET and sub-tomogram analysis to determine a 3D structure for the in situ NPC in a radially dilated conformation. Collectively, our data provide new insights into the structure, assembly, transport mechanism and evolution of the NPC.

## Results

### A composite structure of the isolated NPC

The large size, flexibility and dynamic properties of the NPC in the cell present a formidable challenge to structural studies. Thus, a divide and conquer approach with different methods is needed to determine a comprehensive structure of this massive transport machine. We started with isolated NPCs to allow the collection of single particle data with cryo-EM. S. cerevisiae cells present a tractable system to study isolated NPCs as they lack a nuclear lamina; hence, NPCs can be released gently from the nuclear envelope (NE) with detergent-extraction of a cell cryo-lysate and purified with a single affinity step (7), a process that has been significantly improved upon in this work (Methods). For this project, we reasoned that multi-body processing with density subtraction, focused classification and 3D refinements might achieve α-helical resolution, as a step forward. To minimize domain movements, we used mild DSS crosslinking after a buffer optimized, one-step pullout of NPCs tagged on Mlp1, a major protein in the nuclear basket (Fig. S1A) (7). Cryo-grids were prepared with NPCs exhibiting a wide range of views while supported on a thin carbon film (Fig. S1B, left and right panels), as verified by 2D classification (Fig. S1C). Initial 3D maps were refined with C8 symmetry (Fig. S2A, S2B), followed by focused refinement with appropriate masks to sharpen ring density (Fig. S2C, S2D). After multi-body processing of extracted C8 rotational subunits (C8 protomers) and step-wise refinements of the spoke (Fig. S3, left), the reassembled inner ring had a global resolution of ∼7.6 Å resolution, with 8-fold symmetry along z and two nearly perfect, local 2-fold symmetry axes located 22.5 degrees apart in the xy-plane (Fig. 1A,C,D and Fig. S4A-D).

**Fig. 1.**
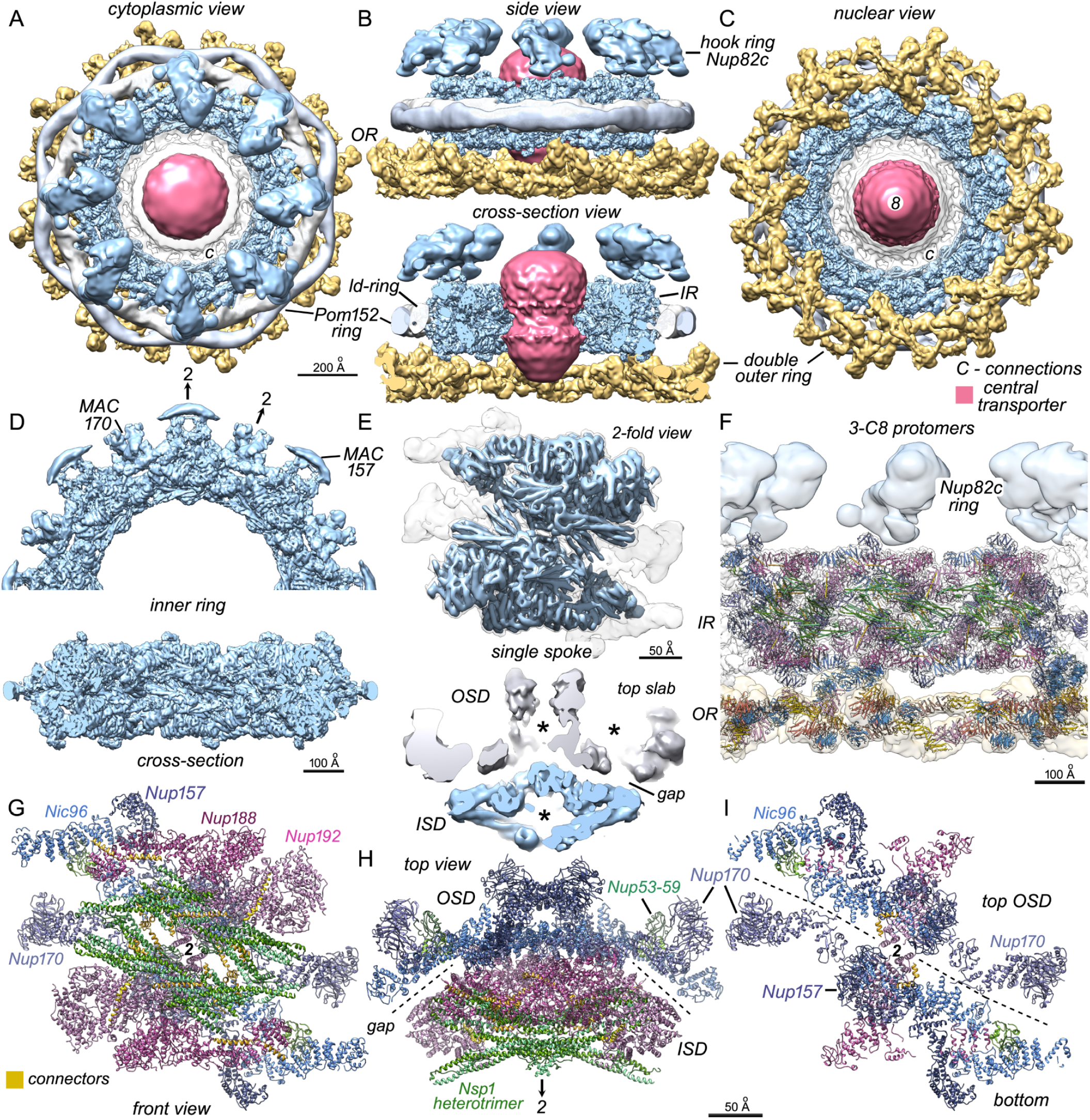
Architecture and structure of the isolated yeast NPC. **A**. A composite 3D map of the isolated NPC is viewed along the 8-fold axis from the cytoplasmic side. Major components are labeled and color coded: double nuclear outer ring (OR, gold), inner ring of spokes (IR, light blue), hook ring/Nup82 complex (blue; note that the full cytoplasmic outer ring has not been resolved), Pom152 ring density (grey), pore membrane remnant (white), central transporter (pink) and spoke-transporter connections at a lower threshold (outlined transparent density, marked “C”). **B**. Side views are shown of the isolated NPC. (top) Full composite 3D map with hook ring/Nup82 complex (Nup82c) indicated. (bottom) A central cross-section is shown with the inner ring (IR, OD=860 Å, ID=450 Å), double outer ring (OD=1160 Å), lipid-detergent ring (ld-ring; a remnant of the pore membrane; ID ∼860 Å), Pom152 ring (average diameter=950 Å) and the uncut central transporter. **C**. A view from the nuclear side shows the double outer ring formed by pairs of Y-shaped Nup84 complexes. **D**. (top) One half of the inner ring (oriented as in panel A) is shown with 2-fold axes (labeled 2) that align with membrane anchor complexes (MAC) for Nup170 and Nup157. (bottom) A cross-section of the inner ring shows α-helical resolution throughout (also see panel E). **E**. (top) A single spoke colored at two thresholds (blue and light gray) reveals α-helices in this feature. (bottom) A central, horizontal slice through the spoke, as viewed from the cytoplasmic side shows the inner and outer spoke domains (ISD, OSD). A gap between them is indicated and large hollows are marked (*). **F**. A cross-section of the NPC is shown with models for three docked spokes and 3 pairs of Y-shaped molecules in the outer ring (oriented as in panel B, bottom). A guide for Nup coloring is presented in panel G and Fig. 2, β-propellers in the Y-complexes are in light blue. **G**. A close-up of a modeled spoke is viewed from the front along the central 2-fold axis. **H**. When viewed from the top (a cytoplasmic view), the spoke is radially separated into two spoke domains (dashed lines, gap; labeled ISD, OSD) that reflect their distinctive functions. **I**. A thin slab viewed along the central 2-fold axis of the spoke reveals a vertical separation of the OSD (dashed lines) into top and bottom halves.

A similar multi-body approach was used on the ring of “hood-shaped” densities (Fig. S2A) to resolve overlapping, Y-shaped Nup84 complexes (30, 31)(Fig. S3, right). This analysis revealed a flat, double outer ring on the nuclear side with a resolution of ∼ 11.3 Å (Fig. 1A-C and S4E-4H). Other regions of the isolated NPC were visualized at lower resolution including the export platform, comprised of flexible Nup82 complexes on the cytoplasmic face (23), the lumenal Pom152 ring and the central transporter with connections to the inner ring (Methods, Fig. S4I-4L). These density maps were combined to create a composite 3D map (Fig. 1A-C); integrative structures of NPC components (7, 15) and crystal structures of the Nup84 complex (32) were then docked into the maps to create models for major sub-assemblies of the isolated yeast NPC (Fig. 1F-I). This has resulted in a model of the yeast NPC based on the highest resolution structures to date (7-11 Å; versus 23-28 Å for previous tomographic models (7, 23). An overview of Nups in the C8 protomer and scaffold is shown in Fig. S5.

### Functional domains of the inner ring at α-helical resolution

A majority of the ∼ 715 α-helices in a spoke were resolved in the single particle structure (Fig. 1D-E). Thus, 28 Nup domains were modeled within the spoke (Fig. 1F-I, S6) with an average root mean square deviation (rmsd) of ∼ 8Å for Cα main chains, when compared to an earlier integrative model (7); this model is independently supported by the mapping of 206 relevant NPC crosslinks to the inner-ring with a 95 % satisfaction (Fig. S8B, 8C). In this model, the protomer of the inner ring is resolved into an inner spoke domain (ISD), comprising two layers, and an outer spoke domain (OSD) with a single layer (see dashed line in Fig. 1H). The ISD is composed of symmetry-related pairs of laterally-offset Nsp1 heterotrimer complexes (Nsp1, Nup49 and Nup57) in the inner layer and two pairs of Nup188 and Nup192 in the adaptin-like layer, with an array of “orphan” densities that correspond to connectors (Fig. 2, see next section). The OSD contains four C-terminal domains (CTDs) of Nic96, which form keystone supports of the inner ring (7); they provide an interaction scaffold for Nup157 and Nup170 (2 copies each) in the membrane interaction layer, while Nup53 and Nup59 heterodimers form bridges between Nic96 CTDs (Fig. 2A and S6C, top left). The OSD can be further delineated into top and bottom halves relative to the central spoke 2-fold axis (dashed line in Fig. 1I, Fig. 2A, middle), with contacts formed by loops extending from α-helical solenoids belonging to Nic96 and Nup170.

**Fig. 2.**
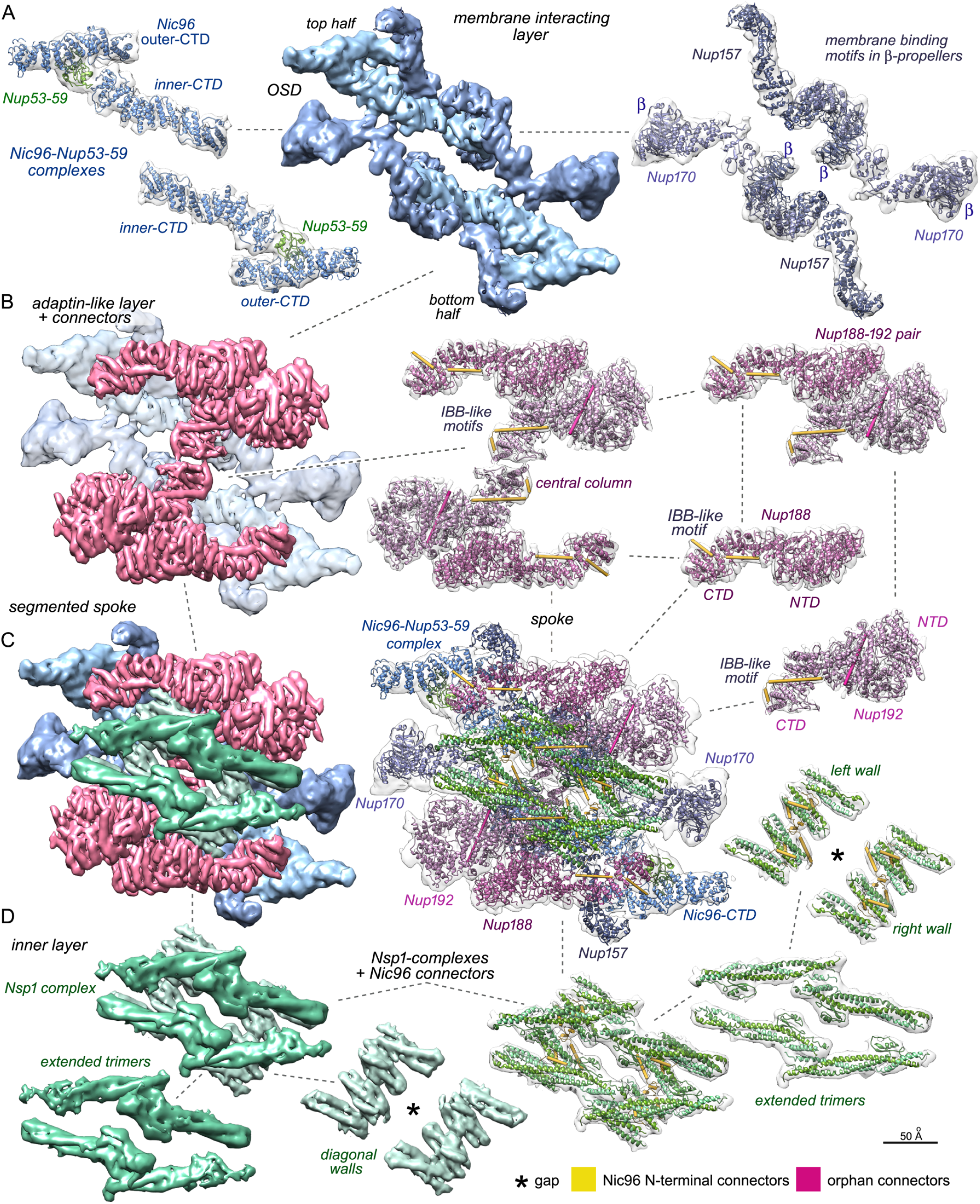
A diagram of the yeast spoke at α-helical resolution. The yeast spoke is comprised of 3 layers that are visualized with segmented 3D maps and docked Nups as ribbons. Inter-connecting dashed lines show the relationship between various Nups in the spoke. **A**.The outer spoke domain is comprised of the membrane interacting layer (center) and contains 2-fold relatedNic96 CTD pairs linked by a Nup53-Nup59 heterodimers (left) and two Nup157-Nup170 pairs (right). **B**. In the inner spoke domain, a central adaptin-like layer contains two copies of laterally-offset Nup188-Nup192 pairs with bound connector helices. Nup192 CTDs form a central column (middle). **C**. The full spoke is shown with segmented 3D maps (left) and ribbons (center). **D**.(left) The inner layer contains pairs of laterally offset Nsp1 complexes related by two-fold symmetry to form the inner face of the spoke. This layer can be further sub-divided into extended trimers that face into the central channel and two diagonal walls, which are formed by shorter, 3-helix bundles. (right) Ribbons for Nsp1 complexes and connector rods are docked into transparent 3D maps. Note the large internal gap that is present between the diagonal walls (asterisk).

The separation of the spoke into two domains with 3 layers is correlated with function (Fig. 1H, Fig. 2C). The ISD forms a scaffold for the transport path; lateral interactions between adjacent Nups stabilize the inner ring and position FG domains from Nsp1, Nup57 and Nup49 within the central transporter. In contrast, the OSD is largely devoted to shaping and stabilizing the pore membrane (7, 17, 20). Thus, Nups 157 and 170 in the membrane interacting layer have amphipathic membrane binding motifs (MBMs) in their β-propellers, while Nup53 contains an additional C-terminal membrane interaction motif (33) (Fig. 2A). In the middle layer, Nups 192 and 188 contain α-helical, adaptin-like domains (Fig. 2B, Fig. S6B) (34, 35) that stabilize the spoke by connecting the inner layer of Nsp1 complexes to the membrane interacting layer (Fig. 2B-D). By comparison with crystal structures and homology models, the CTDs of Nup192 and Nup188 in our α-helical resolution map flex and rearrange when they interact with partner Nups and connectors; for example Nup192 CTDs are bent and stack tail-to-tail to form a central column (Fig. 2B, center panel). The quality of the 3D map is documented further in Fig. S6. On the inner face of the spoke elongated, α-helical coiled coil trimers from four Nsp1 complexes align with their neighbors to create a ring with an inner diameter of ~425 Å that defines the central channel (7, 17, 20) (Fig. 1F, Fig. 2D). Each of the four Nsp1 complexes also contributes a pair of 3-helix bundles that extend inwards in pairs to form two diagonal walls (Fig. 2D and S6A). Thus, each wall is composed of four 3-helix bundles (3HBs) that originate from two Nsp1 heterotrimers, with one pair of 3HBs coming from the top half of the spoke and one pair of 3HBs from the bottom half. This unusual alignment creates a local 2-fold axis for structural elements in each wall.

A detailed examination of the inner ring supports the idea that the NPC evolved for flexibility (7, 36–38), which allows movements both within and between the spokes (this work, (7, 19, 23)) that are facilitated by gaps and voids between Nups (see asterisks in Fig. 1E, bottom panel). Regions in close proximity to the pore membrane exhibit lower resolution (Fig. S4D), due to flexibility that accrues from the architectural division of the spoke and the loss of an intact pore membrane. Multiple contacts between adjacent spokes are formed by limited edge-wise interactions, and provide flexible hinge-like contacts (7). A comparison of our structure with NPCs that have a dilated inner ring in growing S. cerevisiae cells (this work, (23), suggests that the radially-contracted inner ring may represent a “relaxed” conformation. Moreover, our model for the yeast spoke can be docked accurately into cryo-ET maps of human, algal and yeast NPCs with only local changes (17, 19, 23).

### The role of Nic96 connectors in NPC structure and assembly

Biochemical and structural data have begun to fill in the puzzle of how flexible connectors belonging to Nic96, Nup53/59 and their partners may stabilize the spoke (7, 19, 20, 39, 40). Major structural modules of the NPC are held together by flexible connectors that provide strength and resilience in the manner of a suspension bridge (7). In particular, two conserved sequence motifs, known as SLiMs (short linear motifs), have been recognized in the N-termini of yeast Nic96 and its human homolog, Nup93. The first motif is thought to interact with the Nsp1 and human Nup62 complexes, respectively, which are located in the inner layer (20, 33, 41), while the second motif is ambidextrous and interacts with Nup188/Nup192 or human Nup188/Nup205 in the middle, adaptin-like layer (Fig. S7A; (20, 39, 42)).

At the current resolution, orphan densities were revealed that could not be accounted for by docking crystal structures and homology models into the cryo-EM map. In brief, three orphan densities interact with α-helical bundles from each of the four Nsp1 complexes in the diagonal walls and a gap at the center of each wall is filled by 4 orphan densities (Fig. 2D, Fig. 3A-B and S6A), which accounts for 20 orphan elements. We also identified 3 orphan helices that bind to Nup192 and two orphan helices that bind to Nup188, and these Nups are present in two copies each within the spoke. Thus, 30 orphan densities have been mapped within the spoke that arise from flexible connectors.

**Fig. 3.**
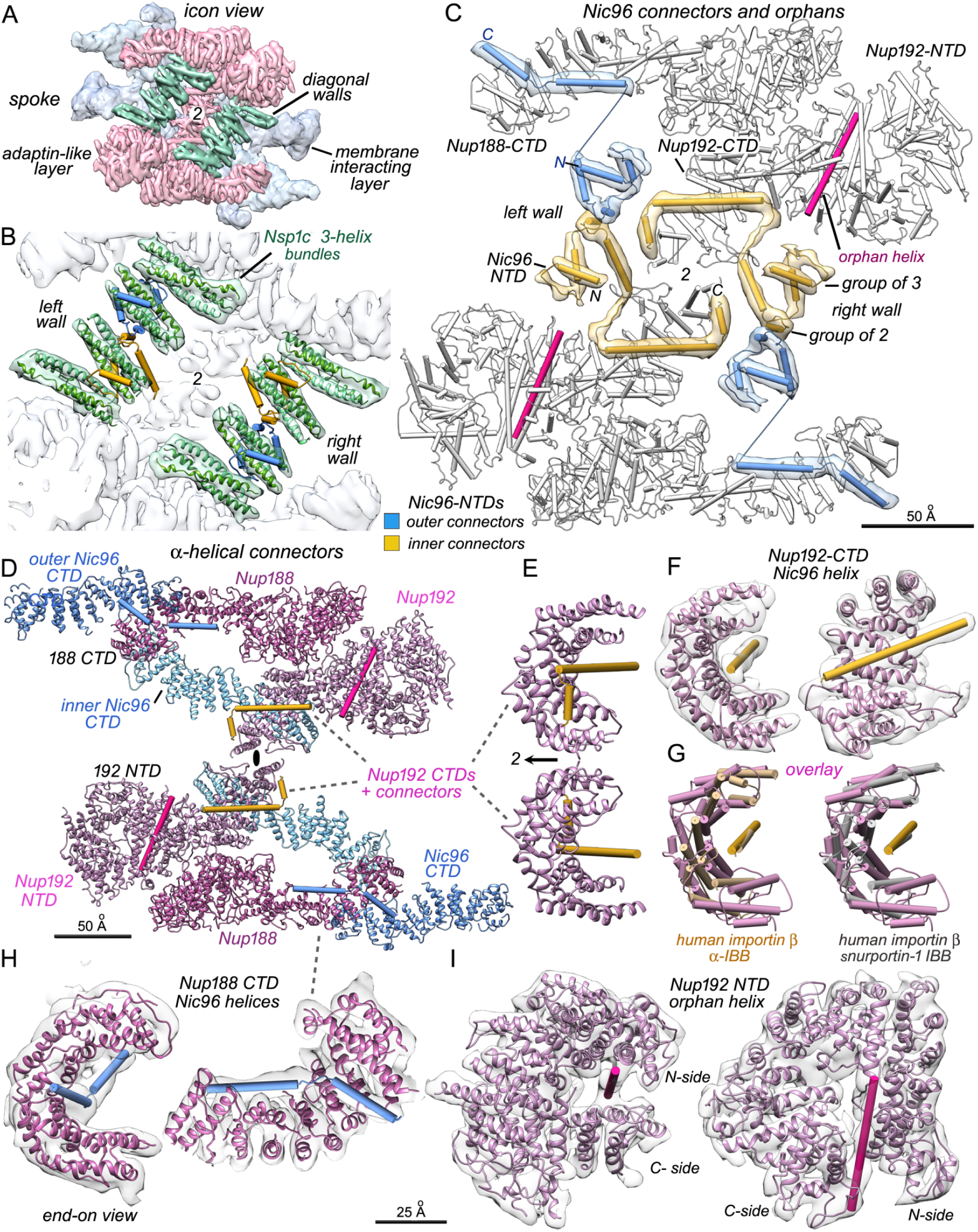
Assignment of orphan structural elements in the spoke and interaction motifs for Nup192 and Nup188 with Nic96 NTDs. **A**. Three-helix bundles from Nsp1 complexes and orphans from Nic96 NTDs form two diagonal walls, which are shown as a green iso-surface with the inwards facing, extended trimers of the Nsp1 complex removed. Color coding and orientation as in Fig. 2. **B**. Four connector α-helices from Nic96 (gold and blue cylinders) form a bridge between opposing half walls, while other connectors bind to 3-helix bundles (green ribbons) in the walls. **C**. Orphan helices and strands are superimposed on Nups 188 and 192 (in white) near the spoke center. Connectors from the inner and outer copies of the Nic96 CTD are colored in gold and blue with overlayed density for these features. Each Nic96 NTD forms a group of 3 structure elements, followed by a group of 2 helices, which are linked to two helices that interact with an NTD of Nup192 or Nup188. A remaining orphan helix in each half spoke is shown in pink. **D**. An overview of connector interactions with Nup α-helical domains is shown. Outer and inner Nic96 CTDs (blue, light blue) form keystone supports behind Nups 192 and 188. CTD tails of Nup192 form IBB-like motifs with Nic96 connector helices and interact with each other at the central 2-fold axis (black ellipse). Nic96 NTD α-helices are shown as cylinders (gold and blue as in panel C). **E**.A close-up is shown of symmetry-related Nup192 IBB-like motifs at the spoke center with the 2-fold axis in the plane. **F**.Two views are shown of a Nup192-IBB motif with the 3D density map super-imposed. A small C-terminal connector helix has been omitted for clarity. **G**. (left) An overlay is shown of a Nup192-IBB motif with a human importin β- IBB crystal structure (in tan, PDB 1QGR). Helices in the arch-like walls have a similar topology. (right) An overlay is shown for a Nup192 motif with the human importin β-snurportin 1 IBB motif (in grey, PDB 2Q5D). **H**.Two views are shown of the Nup188 CTD-Nic96 motif with overlayed 3D density map. The extended arch-like tail of Nup188 binds two consecutive Nic96 N-terminal α-helices in a groove; the binding site for the shorter helix is reminiscent of a transport factor IBB motif. **I**.Two views are shown of a novel interaction motif formed by the Nup192 NTD with a single long helix that may originate from an N-terminal tail of Nup59 (but see Main text). Panels E-I are shown at the same scale.

An accurate identification and topology map of these connectors is needed to understand how the OSD and the ISD are tied together. To solve this problem, we applied an integrative threading approach (43) to compute Nup identity, copy identifier, and sequence segments for each of the structure elements (SEs) that occupy orphan densities (Fig. S7, Methods, Table S2). This resulted in 28 of the 30 orphan SEs being confidently assigned to contiguous sequences in four copies of the Nic96 NTD (Fig. S7E). In our model, the topology of the first group of 3 orphan densities (Fig. 3C, group of 3) is similar to that of a helix-strand-helix motif in the crystal structure of a Nic96 SLiM bound to the CtNsp1 complex (PDB 5CWS; Fig. S8F) (33). Differences between our model and the X-ray structure include a repositioning of the C-terminal 3-helix bundles of each Nsp1 complex within the diagonal wall, and this alters the packing of the Nic96 helix-strand-helix motif. Two closely paired α-helices at the center of the wall (group of 2, Fig. 3C) immediately follow the helix-strand-helix motif in the Nic96 N-terminus (Fig. 3C, S7 and S8A). The final two orphan helices in each Nic96 NTD bind to CTDs of Nup188 or Nup192: these -helix pairs originate from the same sequence region in the outer and inner copies of Nic96, respectively, and contain helices of different lengths (Fig. 3C-E, Fig. 3H, S7 and S8A). Thus, our analysis highlights a remarkable plasticity that allows the same sequence of Nic96 to form two distinctive, -helix pairs that function as connectors and bind to either Nup188 or Nup192 in the spoke. In total, the N-termini of four Nic96 molecules contribute 28 connectors to each spoke with 24 α-helices and 4 strands, while two additional orphan helices are bound to the NTDs of Nup192 (Fig. 3C, S7B, Fig. 7F). These long connectors may originate from Nup59 instead of Nup53, as Nup59 and Nup192 are found in the same sub-complex with affinity capture data (15). However, alternative assignments include connectors from Nup100, Nup116, Nup145N, and possibly from Ndc1 and Pom34, instead of Nup59 (Fig. S7G).

The 30 connectors that have been resolved in our study play important roles in spoke architecture. First, we showed that Nic96 interacts with every other Nup in the inner ring, to form an essential keystone that holds much of this structure together (7); here, we find that N-termini from Nic96 are critical for the recruitment of Nsp1 complexes and tie together the three layers of the spoke (Fig. 2, Fig. 3A-C). This network of connectors could provide the inner ring with enough flexibility to sustain expansion-contraction motions that may occur during transport (see below). Second, our structure also provides key insights into the mechanism by which NPCs assemble and disassemble. Rapid assembly of the interphase NPC may be dependent in part on the ability of flexible connectors to form extensive interactions between Nups in different layers, tying these components together quickly and efficiently into the mature inner ring. Moreover, homologous regions in human Nups are hyper-phosphorylated at mitosis, including the NTD connectors of Nup93 (see predicted sites, Fig. S8A), in a process that disrupts interactions within the spoke and promotes NPC dis-assembly (44–46) (see Discussion).

### Nucleoporins and transport factors share common interaction mechanisms

The extended α-helical solenoids of Nup188 and Nup192 are structurally similar to the karyopherin family of nuclear transport factors, which indicates that Nups and Karyopherins share a common evolutionary origin (15, 24, 34, 35, 47, 48). Here, we show that pair wise interactions are formed by CTDs of Nup192 and Nup188 with α-helices from Nic96 NTDs. These data provide further insight into how the NLS recognition system may have arisen in nucleocytoplasmic transport through divergent evolution and point to a link between spoke assembly and transport. In particular, a long -helix of Nic96 is cupped by a spiral of α-helices in the Nup192 CTD; this structural motif resembles the interaction between karyopherin β and -adaptors for cargo molecules (49, 50). In addition, an adjacent, short helix may stabilize this interaction (Fig. 3D, Fig. 3E), which occurs twice at the spoke center (Fig. 3D-F). This novel motif is overlaid with a crystal structure of karyopherin / importin β1 and the α-helical NLS-like IBB (importin beta binding) domain of karyopherin / importin (Fig. 3G, left panel, PDB 1QGK in tan) (50). A second comparison is shown for the karyopherin / importin β1 complex with the β-snurportin 1 IBB (Fig. 3G, right panel; PDB 3LWW in grey) (49). Importantly, the topology of α-helices in the arch-like walls of the Nup192 CTD and β-importin are similar although some divergence has occurred at the C-terminus of Nup192, because this region interacts with another copy of itself at the spoke center.

The curved C-terminal tail of Nup188 also interacts with two consecutive α-helices from the flexible NTD of Nic96 (see blue rods in Fig. 3D, Fig. 3H, S7). Once again, the curved topology of the Nup188 CTD and its juxtaposition with the shorter Nic96 -helix is reminiscent of a transport factor-IBB interaction (49, 50). The α-helical domains in C-termini of Nup192 and Nup188 were adjusted from previous models to fit the electron density map and to accommodate partner helices from Nic96. As noted, a second novel interaction is formed between the NTD of Nup192 and a long orphan helix (Fig. 3C, Fig. 3D, Fig. 3I).

### A double outer ring in a subset of yeast NPCs

The outer rings in yeast NPCs are formed by the head-to-tail arrangement of 8 Y-shaped Nup84 complexes to form single cytoplasmic and nuclear rings (15, 23, 30, 32, 51). In detergent-solubilized yeast NPCs the outer rings are rather disordered (7), due to inherent flexibility and the loss of stabilizing interactions with the pore membrane. However, we identified a subset of isolated NPCs (∼ 20%) with a double outer ring of Y-complexes and solved this structure at ∼ 11.3 Å resolution (Fig. S3, Fig. S4E-H). The double outer ring rests in a flat orientation relative to the inner ring (Fig. 1A-C) and sits on the side opposite to density assigned to the Nup82 complex. Hence, the double ring is present on the nuclear surface - as confirmed by in situ fluorescence quantitation (see next section). Although a double outer ring of Nup84 Y-complexes has been observed on the nuclear face of the NPC in other species (16, 19, 21), this is the first time two such NPC variants have been identified in the same organism: one with two single outer rings and a second with a single outer ring and a double outer ring.

We modeled 14 Nups (two Y-shaped Nup84 complexes) in the C8 protomer of this novel double ring, starting with Nup domain crystal structures (32). The complete model provides a snapshot of Y-complexes in two different ring geometries (Fig. 4A-D); the ring proximal to the membrane is continuous while the distal ring contains a gap between adjacent Y-complexes (Fig. 4C, top). The two Y-shaped molecules in the C8 protomer form typical staggered contacts (Fig. 4C) that are visible in the offset between V-shaped regions consisting of Nup120 and Nup85/Seh1 (Fig. 4D, right side). Distinct conformations of the Nup84 complex are present in the two rings: the proximal Y-complexes are extended and curved at the ends (when viewed from the side), while the distal Y-complexes are kinked with a different tail conformation (Fig. 4D, left side).

**Fig. 4.**
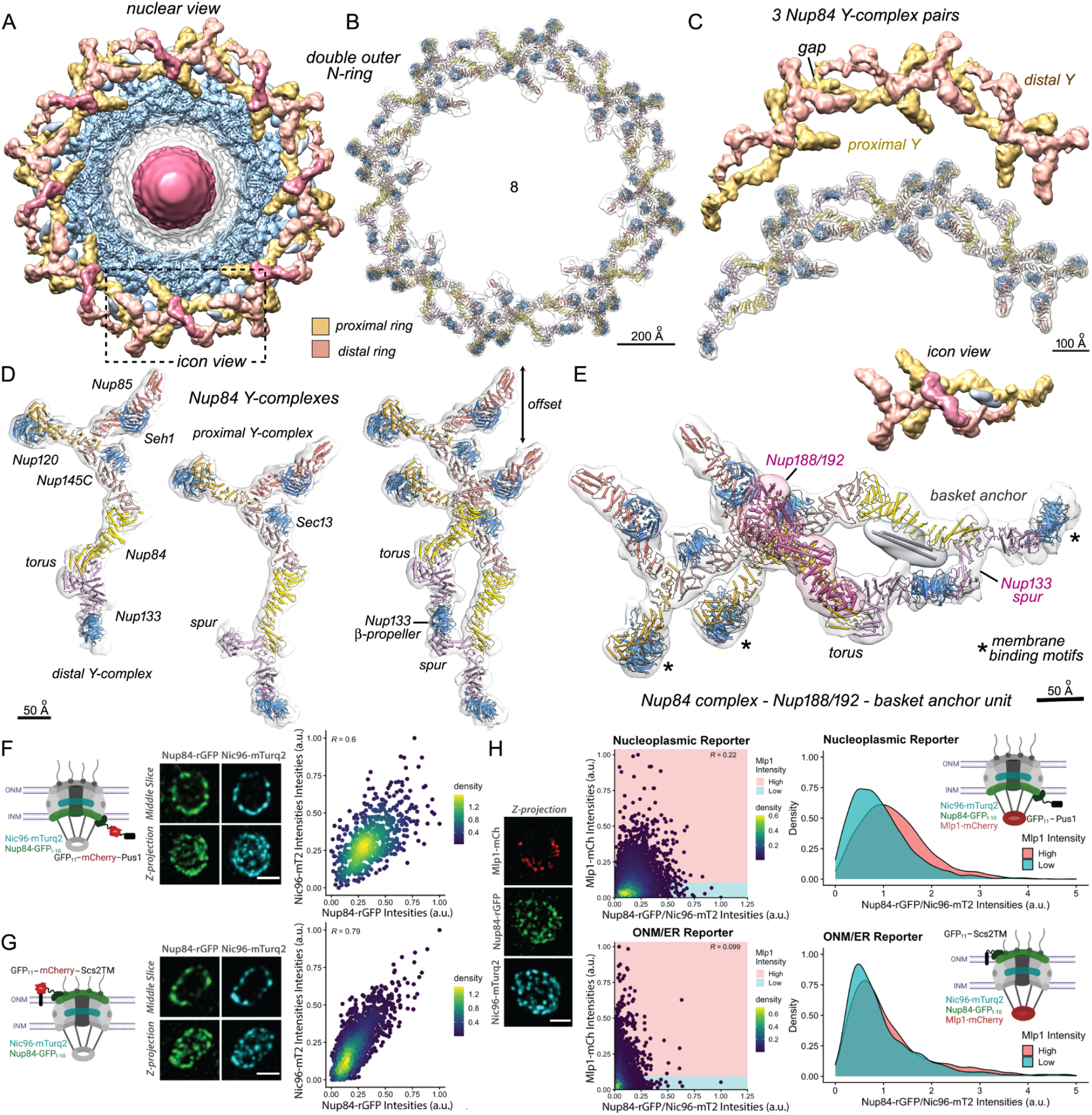
A double outer ring in a subset of yeast NPCs. **A**. A nuclear view is shown of the isolated yeast NPC with a double outer ring segmented into proximal (gold) and distal (tan) rings. Density features for putative Nup188/192 molecules (red) and basket anchors (silver) are also shown. An individual C8 protomer for the double ring is outlined (dashed box, “icon view”). **B**. A model of the double outer ring is shown with an overlayed, transparent density map. **C**. (Upper panel) Three paired Nup84 complexes are shown after segmentation from the 3D density map. A gap is present between neighboring Y-complexes in the distal ring (in tan). (Lower panel) Three pairs of Nup84 complex models in the density map. **D**. Single Nup84 Y-complexes are shown from distal (left) and proximal (right) outer rings along with a staggered pair which forms the C8 protomer for the double ring. A torus-like bulge and an extended Nup133 spur are present in respective tails of the distal and proximal Y-complexes. **E**. A Y-complex pair is shown with a putative Nup188/192 molecule and basket anchor along with an icon view to orient the point of view, relative to panel A. **F-G**. (left) Schematic of the split-GFP system used to visualize outer ring complexes at the nucleoplasmic face of the NPC using a nuclear-localized GFP11-mCherry-Pus1 reporter or at the cytoplasmic face using a GFP11-mCherry-Scs2TM reporter localized to the outer nuclear membrane. Interaction of the GFP11 reporter with Nup84-GFP1-10 results in reconstitution of GFP (rGFP) and fluorescence; (middle and right), representative images and 2D density plots of Nup84-rGFP and Nic96-mTurq2 fluorescence intensities from strains expressing the nucleoplasmic (panel F) or the ONM (panel G) GFP11 reporter. Equivalent reconstitution of rGFP signal for both reporters shows that all NPCs have outer ring complexes on both their nuclear and cytoplasmic sides. A total of 662 (panel F) and 2720 NPCs (panel G) were analyzed. R, Pearson’s correlation coefficient. **H**. (left) A representative image of cells expressing Nup84-GFP1-10, Nic96-mTurq2 and Mlp1-mCherry with a nucleoplasmic GFP11-Pus1 reporter. (middle and right) Density plots showing the distribution of Nup84-rGFP intensities (normalized to Nic96-mTurq2 intensities) for NPCs with Mlp1 intensities greater than (“High”) or below (“Low”) the mean Mlp1 intensity value. Additional rGFP signal in the “High” Mlp1 NPCs for only the nuclear reporter shows that those NPCs have additional nuclear outer rings. A total of 4277 (nucleoplasmic reporter) and 4202 (ONM reporter) NPCs were analyzed. Schematics for panels F-H created with BioRender.

Structural plasticity is exhibited by Y-complexes in NPCs from different species (7, 18, 19, 21, 23); however, by comparison with vertebrates and algae, the tail of the distal Y-complex in the yeast double outer ring shows a more extreme adaptation to changes in local geometry. The β-propellers of Nup120 and Nup133 contain amphipathic MBMs; however, only three of the four β-propellers in each protomer of the double ring may be involved in direct membrane contacts. Flex points located in Nup84 and Nup133 allow sub-domains to form a torus in the tail of the distal Y-complex (Fig. 4D, left) that pulls the N-terminal β-propeller of the distal Nup133 upwards from the putative membrane surface. Instead, the Nup133 β-propeller makes a direct contact with an α-helical domain of Nup133 in an adjacent Y-complex to stabilize the double ring (Fig. 4D, right).

We then identified two additional components within orphan densities that associate with Y-complexes in the double outer ring. First, an orphan density runs along the top of the distal Y-complex (Fig. 4A, Fig. 4E, in red) that strongly resembles the paralogous Nup188 and Nup192. We were able to dock a copy of Nup188 into this density (Fig. 4E), while Nup192 did not fit as well. This unexpected moonlighting by Nup188/192 parallels earlier observations that Nup205 and/or Nup188 are present in vertebrate double outer rings (16, 17, 20, 21). However, yeast Nup188/192 sits on the upper surface of the flat double ring, whereas two binding sites for vertebrate homologs in their highly tilted double rings occupy inward facing regions on the underside of the Y-complexes (21). While the function of this moonlighting subunit in the yeast double ring is not known, we hypothesize that Nup188/192 may stabilize the unusual tail conformation of the distal Y-complex and confer additional rigidity to the double ring. Second, we found a density rod (∼ 100 Å long) adjacent to Nup84 and Nup133 in the membrane proximal Y-complex that could be modeled with a two-helix coiled coil (Fig. 4E). We tentatively identify this orphan density as a basket anchor because a similar feature is observed in the in situ NPC (see below), and this feature is consistent with the fact that these NPCs were purified by pulling on genomically-tagged-Mlp1, a major basket protein. Together, these observations highlight the remarkable structural versatility of the outer ring.

### Each nucleus contains NPC variants with single or double outer rings on their nucleoplasmic faces

Previous measurements of Nup stoichiometry for isolated and in situ NPCs are consistent with two single outer rings per NPC with one ring on each face (7, 15, 52). This raises the question of how NPCs with additional outer rings may be compatible with these measurements, an issue we have addressed with quantitative fluorescence measurements in yeast cells (53). First, we asked whether all NPCs have outer rings on both their nuclear and cytoplasmic faces. We tagged Nup84 (an outer ring component) with one module of a split GFP to monitor the outer rings: the remaining module on a reporter was specifically localized to either the nucleoplasmic or cytoplasmic side of the NE. In this way, the presence of an outer ring on either the nuclear or cytoplasmic face of the NPC could be determined by measuring reconstitution of the full fluorescent GFP signal with appropriately localized reporters (Methods, Fig. 4F-G) (53). The distribution of GFP signal indicates that indeed, all NPCs have outer rings on their nuclear and cytoplasmic sides, in agreement with previous structural and stoichiometry data (Fig. 4F-G) (7, 23, 52, 54).

Yeast nuclei are divided into two morphologically and functionally distinct regions: the nucleolus, which fills roughly one-third of the nucleus and forms a dense crescent appressed to the inner nuclear membrane, while the remaining region is filled with non-nucleolar chromatin. The NEs over these two regions can be distinguished, as NPCs localized over the dense crescent are depleted in the basket protein Mlp1 (55–57). We confirmed that there is a population of NPCs located over the dense crescent with low levels of Mlp1 and a population that is excluded from the dense crescent with higher levels of Mlp1. Moreover, isolated NPCs with a sub-population containing a double outer ring were purified by using Mlp1 as an affinity handle. We therefore hypothesized that Mlp1 abundance may be correlated with NPCs that carry a double outer ring. To test this, we measured the abundance of Nup84 as a marker for Y-complexes in NPCs with “high” Mlp1 signal versus those with “low” Mlp1 signal. Moreover, by using a reporter that specifically localized to either the nucleoplasmic or cytoplasmic side, we could ascertain the side with additional Nup84. Signal distribution curves show that a subset (∼ 20%) of the “high” Mlp1 NPCs have, on average, more outer ring signal than “low” Mlp1 NPCs, and that this extra outer ring signal is on the nuclear side. Thus, the data are consistent with double outer rings being located on the same side as the nuclear basket. It is important to note that most NPCs (>70%) do not display the extra Nup84 signal attributed to Y-complexes in the outer ring. Our estimate of 20% for isolated particles with a double nuclear outer ring, based on single particle 3D classification and in cell fluorescence, likely represents an upper bound and is consistent with the precision of previous stoichiometry measurements (7, 52), such that a 1:1 arrangement of nuclear and cytoplasmic outer rings reflects the predominant form of the NPC.

When taken together, the data are consistent with the presence of at least three NPC variants in yeast cells. One form has two single outer rings that frame the inner ring (Form I), while a second form has a single outer ring on the cytoplasmic surface paired with a double outer ring on the nuclear surface (Form II), and both forms have nuclear baskets. A third NPC variant (Form III) has two single outer rings, lacks the nuclear basket and is located over the dense crescent. Notably, we find that different populations of NPCs co-exist within individual cells, rather than in different cell types.

### A comprehensive structure of the yeast NPC in situ

Our analysis of the isolated NPC has provided detailed information about the fine structure, arrangement and connectivity of Nups. However, the NPC is a dynamic assembly that has long been understood to undergo conformational changes with a range of effective diameters (28, 29, 58), and recent studies have shown that the inner ring can undergo a significant dilation in cells (19, 23). To provide a comprehensive picture of the NPC, we used cryo-ET of cryo-FIB milled yeast cells harvested in mid-logarithmic phase (Fig. S9A-H and subtomogram analysis to generate a 3D density map of the in situ NPC. First, we aligned full NPCs and then performed symmetry expansion of individual C8 protomers, which were refined to a local resolution of ∼ 30-40 Å for the core scaffold (Methods; Table S3, Fig. S10A-B). Features in our map are comparable to those in a recently published structure (23), (Fig. S10C-D, Methods) and importantly, our analysis reveals additional features including nuclear basket anchors, connections to FG domains in the central channel, and pore membrane contacts.

We leveraged models for the spoke and Nup84 complex from the isolated NPC, along with an integrative model of the Nup82 complex (9) and data from a recent study (23) to build a nearly complete model of the in situ NPC. In this synergistic analysis, the inner and outer rings were modeled as rigid bodies, starting with conformations observed in isolated NPCs, followed by manual rebuilding and MDFF refinement (59). We then segmented relevant rings from the 3D density map to create a composite map of the in situ NPC. An overview of the in situ NPC with the pore membrane is shown in Fig. 5A (panels 1-3) along with an exploded view of the core scaffold and its constituent rings (Fig. 5A, panels 4-6).

**Fig. 5.**
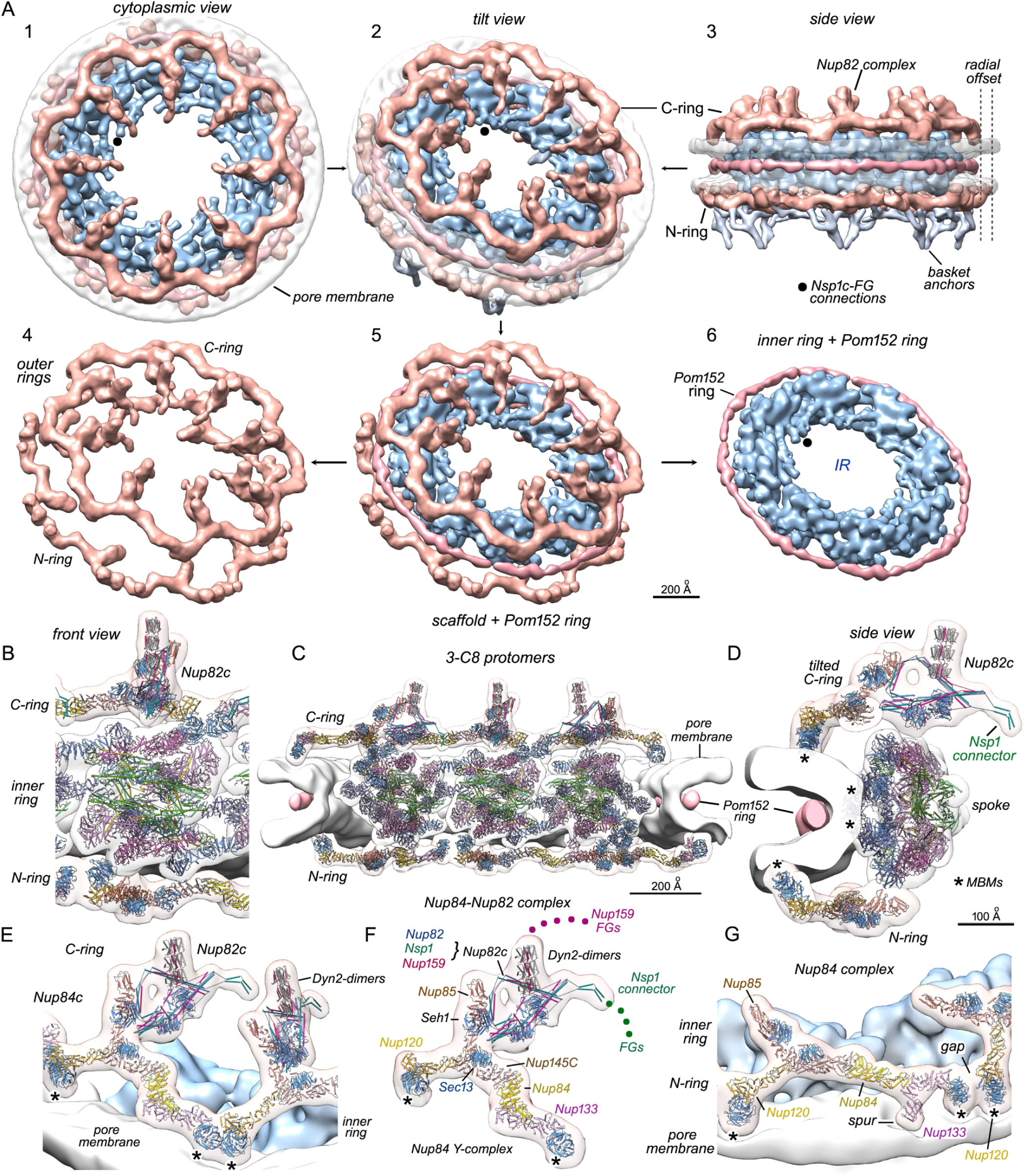
An overview of the core scaffold in the cryo-ET map of the in situ NPC. **A**. (panels 1-3) Cytoplasmic, tilted and side views are shown for the in situ yeast NPC reconstituted from the 1.5X protomer density map. Segmented features correspond to the inner ring (blue), the outer rings (tan) including the cytoplasmic outer ring (C-ring) and nucleoplasmic outer ring (N-ring), the Pom152 ring (red-brown), pore membrane (transparent white) and basket anchors (silver). Putative connections from Nsp1 complexes to FG domains in the central channel are indicated with black dots. The radial offset in diameter of the outer rings (∼50 Å per side) is indicated with grey dashed lines. (panels 4-6) The core scaffold is shown with the lumenal Pom152 ring and is separated into component rings. **B**. A front view of a nearly complete protomer is shown with docked models for the spoke, outer rings and Nup82 complex (Nup82c). Panels B, D, and E-G are at the same scale. **C**. A cross-section of the in situ NPC is shown with molecular models for 3 protomers, the pore membrane and Pom152 ring. **D**. A side view is shown of a nearly complete protomer with molecular models. Approximate positions of membrane binding motifs (MBMs) in the β-propellers are indicated (*). **E**./F. A molecular model for the tilted, cytoplasmic outer ring (C-ring) is shown along with the Nup82 complex (Nup82c). Approximate positions of MBMs are indicated (*). In panel F, Nups are labeled along with colored dots for N-terminal FG domains of Nup159 and Nsp1. **G**. A tilted side view is shown of the nucleoplasmic outer ring with docked models. A gap is present between adjacent Y-complexes and approximate positions of MBMs are indicated (*).

The canonical yeast NPC has single outer rings on both faces of the inner ring and our 3D map of the in situ complex captures this major form. The inner ring has a radially-expanded configuration (Fig. 5A, panel 1) (19, 23)), with a concomitant expansion of the pore membrane and Pom152 ring (Fig. 5A, panel 6). A direct comparison of isolated and in situ NPC structures provides insights into membrane contacts of the scaffold (Supplementary Results and Discussion; Fig. S10A-F) and suggests that the Pom152 ring, which is linked to the spokes at 8 points (Fig. S11G, Fig. S11H), may act as a belt to restrict the extent of dilation by the inner ring and pore membrane (shown schematically in Fig. S8E). Moreover, there is a striking difference in the geometry and diameter (∼ 100 Å) of the opposed outer rings in the in situ NPC (see radial offset, Fig. 5A, panel 3). The cytoplasmic outer ring is tilted relative to the plane of the NE, due to interactions with the Nup82 complex, while the outer ring on the nuclear face is much flatter as it spans the interface between the membrane surface and inner ring. Three C8 protomers in our model show the relationship between the inner and outer rings (Fig. 5C), which is revealed in more detail in two views of a single protomer (Fig. 5B, Fig. 5D). In the spoke, 3-helix bundles from opposing Nsp1 complexes extend into the interior to form two diagonal walls which tie the top and bottom halves together (Fig. 6A, panels 1, 2). A cavity is located between the diagonal walls and lies in front of a central column formed by Nup192 CTDs (Fig. 6A, panels 2-4), while the membrane interacting layer is resolved at higher radius (Fig. 6A, panel 5). A local asymmetry in the spoke may arise in part from differential contacts to the outer rings (Fig. S6B, right panel). Thus, half spokes are in slightly different environments with regards to the outer rings and pore membrane. For example, Nup188 in the cytoplasmic half of the spoke has undergone a larger upwards movement toward the cytoplasmic outer ring, relative to its symmetry mate.

**Fig. 6.**
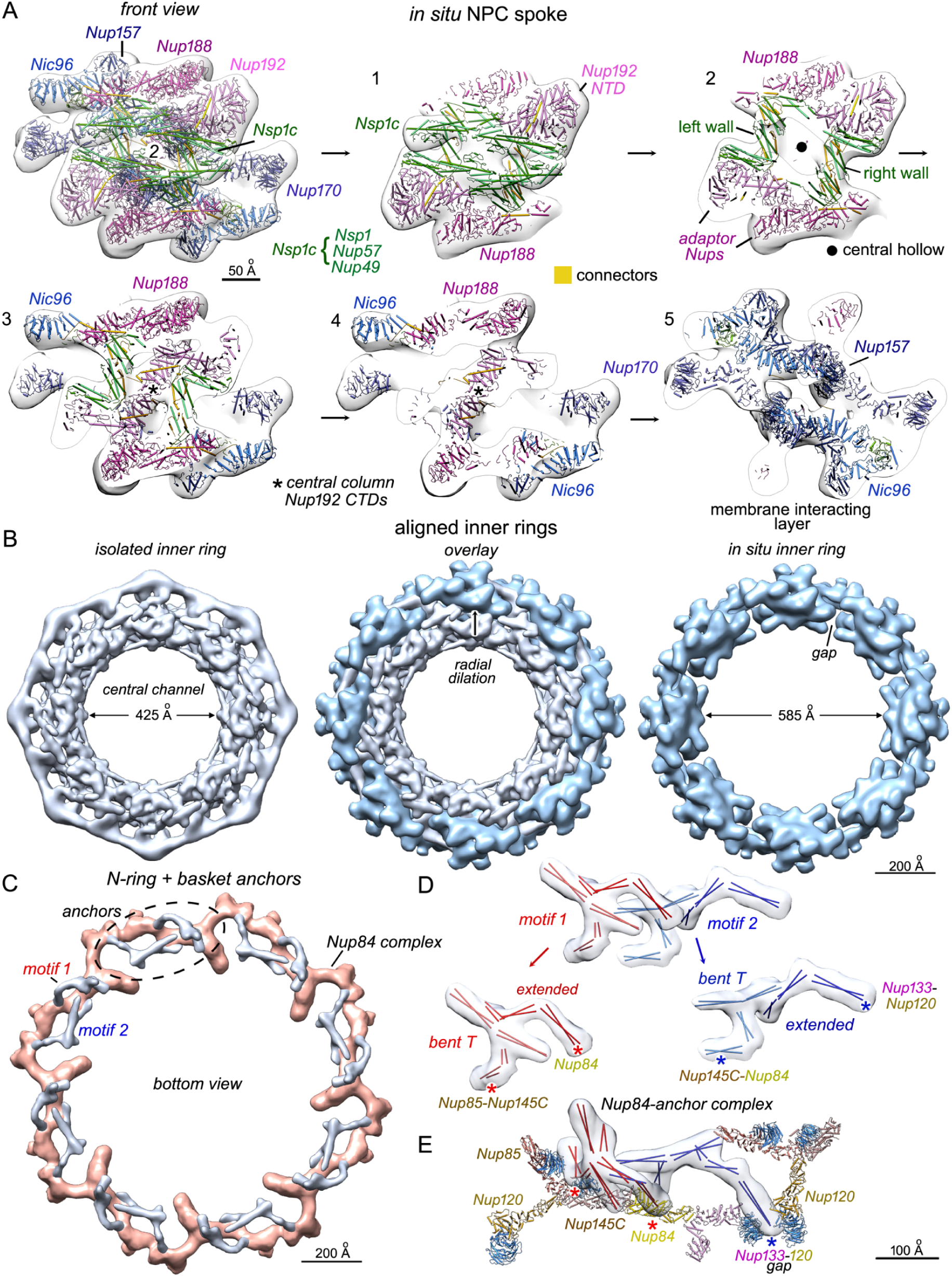
Architecture of the in situ spoke, dilation of the inner ring and identification of putative basket anchors. **A**. Cross-sections are shown through the in situ spoke in 6 panels, as viewed along the central 2-fold axis (2) with docked molecular models for the Nups. The quality of the electron density is sufficient to identify two diagonal walls with a central hollow, a central column formed by Nup192 CTDs and to delineate most of the Nups. **B**. Aligned inner rings from the isolated (left) and in situ NPC (right) are overlaid in the center panel. The radial dilation that occurs between the two conformations is indicated by a vertical arrow. Note that the isolated inner ring has been low pass filtered to match the resolution of the in situ ring and retains part of the ld-ring at higher radius. The approximate diameter of the central channel is indicated for both configurations, as measured from a central protrusion of the Nsp1 complexes. In addition, radial expansion of the inner ring creates a small rotational offset between the local 2-fold axes of opposing spokes (right panel). **C**. The nuclear outer ring with basket anchor complexes (in grey) is viewed from the nucleoplasm. Two anchor motifs (in silver) are present for each Y-complex (dashed oval). **D**. Basket anchor complexes are divided into two similar structural motifs (in red and blue) that were mapped with 2-helix coiled coils (as rods). Each motif contains a 4-piece feature shaped like a “bent T” and an extended 3-piece structure element. Sites used for docking these anchor motifs onto the Nup84 complex are indicated by colored asterisks and their contact points are labeled. **E**. A tilted view of a Nup84 Y-complex in the N-ring is shown with associated basket anchor motifs (in red and blue)

The Nup82 complex interacts with Nup85 in the Y-complex of the cytoplasmic outer ring and overhangs the central channel (Fig. 5E, Fig. 5F). We also modeled three Dyn2 dimers with bound Nup159 peptides containing DID repeats (10, 60) in a prominent, vertically-extended feature of the Nup82 complex; however, the precise rotational alignment of the Dyn2 dimers has not been determined. This feature may direct the unstructured, N-terminal FG domains of Nup159 into a region above the central channel along with Nup42 and Dbp5, an RNA helicase. In addition, connector density extends from a pair of Nsp1 molecules into the transport path (Fig. 5F, see FGs; (23)). Conversely, the single outer ring on the nuclear side in our model has an unusual conformation: the ring of Nup84 complexes is incomplete (Fig. 5G) with a significant gap between adjacent Y-complexes that is bounded by the β-propellers of Nups 133 and 120. These gaps increase the circumference and diameter of the outer ring on the nuclear side, relative to the outer ring on the cytoplasmic side (Fig. 5A, panel 3). A possible explanation for the role of these gaps is provided by our analysis of putative basket anchors (see next section). Unexpectedly, both the N-terminal β-propeller and spur domain of Nup133 appear to interact with the surface of the pore membrane (Fig. 5G). Finally, we note that a 3D map of the in situ NPC with a double outer ring has not been determined, due to the low frequency of this feature in our data set.

A comparison between aligned inner rings of the isolated and in situ NPC underscores this assembly’s inherent plasticity. The spoke, and by inference the pore membrane (7, 17), undergo a significant (∼ 80 Å) radial movement during dilation of the inner ring (Fig. 6B). Hence, the diameter of the central transport conduit is increased by ∼ 160 Å, which nearly doubles the volume that is accessible to FG domains which project from the scaffold into the central channel. Gaps are formed between adjacent spokes in the inner ring in situ (Fig. 6B, right panel) and lateral contacts observed in the isolated NPC are lost, while four Nsp1 complexes in each spoke remain roughly aligned and face into the central channel. Nup170 β-propellers in adjacent spokes reorient through changes in their α-helical domains to maintain contact with the displaced pore membrane and with each other, while the NTD of Nup192 and a CTD of Nic96 reorient to make an inter-spoke contact.

The structural plasticity exhibited by the yeast inner ring may be facilitated by a lack of extensive linkers between the spokes and outer rings, by limited interactions between adjacent spokes and by internal gaps within each spoke. The aligned models of isolated and in situ spokes have an average rmsd of ∼ 9.5Å for Nup C backbones (after excluding Nup170). Hence, radial displacement of the spoke during dilation is accompanied by small but significant local changes in Nup positions, coupled with larger rearrangements at the periphery. In retrospect, the conformation of the inner ring is not determined solely by the presence or absence of the pore membrane, as NPCs in isolated NEs have a contracted conformation (17, 20), rather than the expanded configuration observed in situ (19, 23). We suggest that a radially-dilated configuration of the inner ring with an expanded membrane pore may represent a “tense state”, which can relax back to a more compact conformation as typified by isolated NPCs (7, 17, 20).

### Basket anchor pairs on the nuclear outer ring

The presence of a nuclear basket has been established in multiple organisms, including yeast, although with little structural detail (61, 62). The paralogs Mlp1 and Mlp2 are the major basket components in yeast and are predicted to be coiled coil proteins (57, 63). The improved resolution of our 3D map revealed a pair of bifurcated densities that associate with each Nup84 Y-complex in the outer ring and point down towards the nuclear interior (Fig. 5A, panel 3, in silver; Fig. 6C, dashed ellipse). Their extended shape and position on the nuclear outer ring suggests that these features may correspond to anchor regions of the flexible nuclear basket. We traced the paths of these putative basket anchors and docked 14 segments containing two-helix coiled coils into the density (Fig. 6D), which may correspond to anchoring regions in Mlp1 and Mlp2 (63). The resulting model contains two similar motifs (Fig. 6D, in red and blue), each with a “bent T” feature that anchors this density to the Y-complex and an elongated, structural element with 3 segments that extends from a binding site on the outer ring (Fig. 6D; contact sites are marked with red and blue asterisks). A close-up view shows how the red anchor motif is linked to a Y-complex: the base of the bent T interacts with the Nup85-Nup145C interface and a head-on interaction occurs between the elongated structural element and Nup84 (Fig. 6D, Fig. 6E, red asterisks). The blue anchor motif also has two anchor points on the Y-complex but only one is visible in this view (Fig. 6E, blue asterisk). In this case, we placed a short coiled-coil from the base of the “bent T” into an orphan density associated with the Nup145C-Nup84 interface that is located over the V-shaped region, while the elongated element originates from a gap between adjacent Y-complexes, which is bounded by β-propellers of Nups 133 and 120. Thus, our model for the basket anchors provides a rationale for the presence of gaps between adjacent Nup84 Y-complexes that increase the diameter of the nuclear outer ring.

The topology of the basket anchors suggests that coiled coils and segments from multiple proteins, including Mlp1/2, Nup145N and Nup60 (7), could contribute to these features. At this stage, the docked rods serve only to highlight density features in our model. However, the striking observation of two topologically similar motifs that associate with each Y-complex serves to validate these features and also suggests that the recurring theme of Nup duplicates or paralog pairs within the NPC may pertain to the basket anchors (e.g. Mlp1 and Mlp2) (7, 15, 24, 47).

### Nsp1 complex connections to FG domains in the central channel

The central transporter and connections from the inner ring have been visualized previously in isolated vertebrate and yeast NPCs (7, 64–66). However, the central transporter has been described as a “ghost in the machine” due to its variable appearance (Akey, 2010). Importantly, the NPC scaffold contains anchor points for flexible FG repeat domains that extend radially into the central channel to form the transporter (7, 11). In this study, the central transporter was observed in individual NPCs and 2D class averages (Fig. S1B-C). In addition, visible connections span a ∼ 50-80 Å gap between the spokes and central transporter in 2D class averages (Fig. S1C).

We focused initially on connections in the isolated NPC that link Nsp1 complexes to their flexible N-terminal FG domains, which account for ∼ 38% of FG mass in the central channel (Table S4; (7)). Focused classification provided an improved 3D density map in which the connections were resolved at ∼ 20Å resolution (Fig. S4K, Fig. S4L). Four connections are present in two groups per spoke that span the gap at the waist of the central transporter, with one group located at the central 2-fold axis of the spoke and a second group at the boundary between spokes (Fig. 1A, Fig. 1C, Fig. 7A, Fig. 7C). Visualizing individual links will require higher resolution data; however, three rods per Nsp1 complex were readily docked into the density map – as linear markers – to track these features (Fig. 7A, Fig. 7C, red and blue rods). This connection pattern can be explained by trimeric coiled coils from Nsp1 complexes that form a cylindrical array in the inner ring. In brief, their N-termini are positioned accurately in our 3D map at 7.6 Å resolution and the pattern of connections arises from the lateral off-set between each Nsp1 complex in the half spoke, convolved with the local 2-fold symmetry (7). This places N-termini of the Nsp1 complexes at the connection densities where their flexible FG repeats may extend into the transport path.

**Fig. 7.**
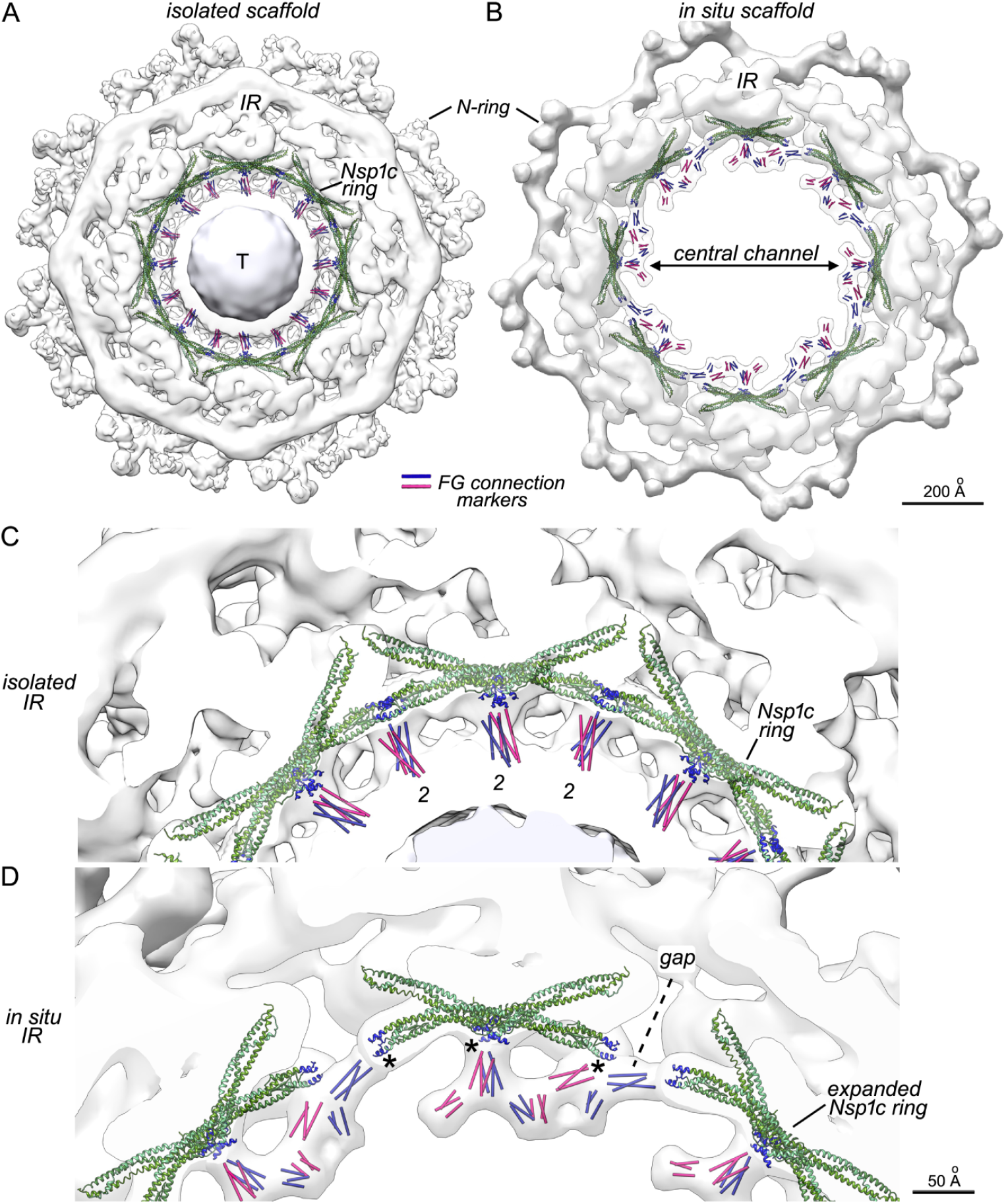
Nsp1 complex connections to FG repeats in the central channel. **A**. A top view is shown for a consensus 3D density map of the isolated NPC scaffold (in white) with the outer C-ring density removed. The central transporter (in silver) is present with a threshold appropriate to the intermediate resolution of the map, while the connections are displayed at a lower threshold. Extended trimers of the Nsp1 complex form a ring (in green) with their N-termini colored in dark blue; connecting bridges are modeled with 3 red and 3 blue rods that were used to map linkers from Nsp1-complexes located above and below the central plane. Local resolution maps for the transporter and connections are shown in Fig. **??**4 (page 2). **B**. A cytoplasmic view of the in situ NPC scaffold without the outer C-ring reveals connecting bridges from Nsp1 complexes that extend into the central channel. **C**. Well resolved connections between the inner ring of the isolated core scaffold and waist of the central transporter are aligned with N-termini (dark blue) of Nsp1 heterotrimers (in green) at local 2-fold axes (2). **D**. Connections in the inner ring of the in situ NPC originate from three points on each spoke (*) that align with the N-termini of Nsp1 complexes (dark blue). Radial expansion creates a gap between adjacent spokes and separates the extended, coiled coil trimers of adjacent Nsp1 complexes; this leads to a local rearrangement of the connections and a new lateral contact between adjacent spokes at the gap.

Our 3D map of an in situ NPC also displays prominent connections from each spoke that extend into the central channel but with a different configuration (Fig. 7B, Fig. 7D, see asterisks). This pattern of connections (again modeled with linear markers) can be explained as follows: in the transition from a contracted to a dilated configuration, each spoke moves radially outwards and a gap opens between adjacent spokes and Nsp1 complexes within the cylindrical array (compare Fig. 7C, Fig. 7D). This causes paired connections at the interface between adjacent spokes in the radially contracted inner ring to split laterally and creates a visible bridge for each Nsp1 complex at the lateral edges of each spoke, with their N-termini facing the gap. At the same time, connections in the center group remain intact as they leave the spoke and then split as they enter the central channel. Thus, dilation of the inner ring creates a pattern of visible connections for each spoke that once again can be accounted for by the distribution of Nsp1 complexes. Moreover, the reorganization of FG connections during radial dilation provides a new lateral contact between adjacent spokes that may compensate for interactions that are lost within the gap (dashed line, Fig. 7D).

For the isolated NPC, we were able to calculate a global average of the central transporter after focused 3D classification to eliminate noisy particles. This region is viewed at a higher threshold to provide a realistic view, due to its low resolution, flexibility and the entrapment of cargo complexes (7) (Fig. 1A-C). The central transporter has two cylindrical, high density regions with a less dense, central waist (∼ 310 Å diameter by ∼ 460-500 Å height) and connecting densities from the spokes encircle the waist (Fig. 7A, Fig. 7C). Based on local resolution (Fig. S4I, Fig. S4J), transporter density is less ordered near the central 8-fold axis and at regions facing into the cytoplasm and nucleoplasm. This is consistent with the expected behavior of flexible FG domains as they extend from their connection points into the upper and lower halves of the central transporter (7, 67). In addition, domains from other FG Nups probably contribute to cylindrical regions of the transporter (Table S4).

Collectively, our data suggests that the hour-glass shaped transporter is created by densely-packed FG domains within the central transport conduit, along with transport factors and cargo complexes that may be trapped on- or near-axis within the FG domains (7). We note that heterogeneous density is present for the central transporter in our 3D map of the in situ NPC after per particle refinement; however, a much larger data set will be required to resolve this feature. Even so, the transition from a radially contracted inner ring to a dilated configuration increases the volume of the central channel by ∼ 2-fold. We suggest that this dramatic expansion of the inner ring may alter the distribution of FG domains within the in situ NPC to facilitate transport (Fig. 8D).

**Fig. 8.**
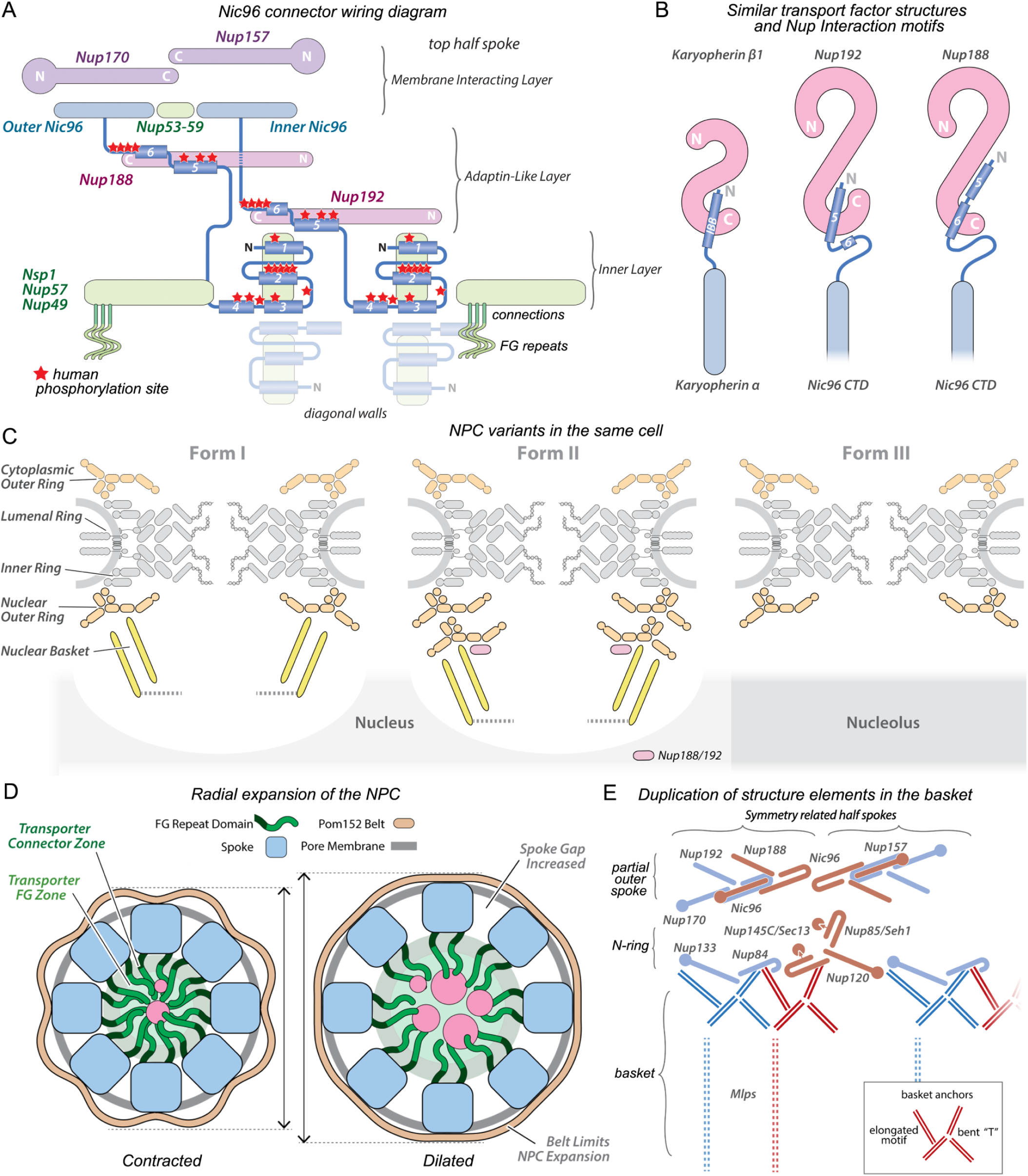
Summary of major functional results on the yeast NPC A. **A** wiring diagram of extended, flexible connectors from Nic96 that tie together the three major structural and functional layers of the yeast spoke. These connectors extend from Nic96 CTDs in the membrane interacting layer to the diagonal walls of the inner layer. and may play important roles in assembly. Red stars indicate approximate positions of phosphorylation sites in the human Nup93, a Nic96 homolog, that are involved in the regulation of mitotic disassembly and reassembly. Color coding: blue-purple for membrane interacting layer with Nic96 CTD, red - adaptin-like layer and green for the inner layer. The extended trimers of the Nsp1 complex have been truncated in laterally to allow visualization of the two diagonal walls that are deeper within the spoke.. **B**. Cartoon comparing similar interaction motifs for Importin-β/karyopherin β1 and the IBB helix from importin-/karyopherin, with Nic96 N-terminal helices bound to Nup192 and Nup188. The similarities suggest a common evolutionary lineage for Nups in the NPC scaffold and transport factors. **C**. Structural and functional variants of NPCs are present in the same yeast cell including: Form I with singe outer rings and a basket, Form II with a single cytoplasmic outer ring and a double nuclear outer ring plus basket, and Form III with two single outer rings and no basket (the latter is localized over the nucleolus). **D**. A diagram of two conformations of the yeast NPC highlights major changes that occur upon dilation of the inner ring, including radial expansion of the spokes with formation of gaps between adjacent spokes; this is accompanied by a displacement of the pore membrane and distention of the Pom152 ring. Radial expansion of the in situ NPC nearly doubles the accessible volume in the central channel for FG domains that extend from the spokes via a zone of connections. The Pom152 ring may act like a belt to help restrict the extent of radial expansion.

### Membrane interactions of the inner ring

Interactions with the NE are critical for assembly and for maintaining the correct positioning of the NPC within the pore membrane aperture. In the isolated NPC, a large remnant of detergent-resistant lipids and detergent form a ring that encircles the inner ring at a radius of ∼ 430 Å (ld-ring, Fig. 1A-B). This ld-ring is centered at the midline and interactions with the transmembrane nucleoporins Pom152, Pom34 and Ndc1 can be identified, as density is partly resolved for their transmembrane domains (TMDs) (Fig. 1D, dashed ovals in S11A, top view, S11B). Fig. 2g

The data show a clustering of amphipathic MBMs in β-propellers from Nup157 and Nup170 (68–71) located on the outer surface of the inner ring where they contact the pore membrane. These sites alternate around the periphery at 16 equispaced positions that correspond to local 2-fold axes (Fig. 1D and S11), and their interactions are consistent with the protocoatomer hypothesis (7, 24). Our 3D density map for the isolated NPC, along with peptide crosslink data from mass spectrometry (7), indicates that β-propellers in Nup157 interact with Pom152 near the central 2-fold axis of the spoke (black asterisks, Fig. S11). Based on the size of the TMDs in the 3D map, Nup170 β-propellers and Nup53-59 heterodimers from adjacent spokes appear to interact with Ndc1 dimers in the isolated NPC (blue asterisk, Fig. S11) as suggested by previous studies (72, 73). The membrane interacting layer of the spoke is formed by two pairs of Nups 157 and 170, four Nic96 CTDs and two Nup53-59 heterodimers. In the isolated NPC, these layers are arranged in a nearly continuous band on the membrane-facing surface of the inner ring with MBMs from Nups 157 and 170 near the midline (Fig. S11, red dots). Similar membrane contacts are present in the in situ NPC, although with a noticeably larger spacing between adjacent membrane interacting layers (Fig. S11, Fig. S11E, red dots in Fig. S11F). We surmise that MBMs in Nup157 and 170 may interact with the membrane, they may bind to membrane protein anchors or they could have both functions.

Paralog pairs of Nup53-59 are located between adjacent Nic96 CTDs in each half spoke and have amphipathic helices at their C-termini that help tether the inner ring to the pore membrane (cf. Fig. S22 in (20); (7, 74)). Thus, Nup53 in each Nup53-59 heterodimer is positioned in close proximity to the ld-ring in isolated NPCs (Fig. S11C, green dots) and direct membrane contacts are visualized in the in situ pore complex (Fig. S11E, S11F, green asterisks and dots). However, a local asymmetry is present in the spoke at higher radius that may alter membrane interactions, as contacts for Nup170, Nups 53-59 and Nup157 are more extensive on the left side of the spoke, when viewed from the cytoplasm (see asterisks, Fig. S11E). Thus, our data suggest that spoke contacts with Pom152 and Ndc1 may follow the local 2-fold symmetry of the radially-contracted inner ring, while these interactions may be altered during radial expansion.

In the isolated NPC, two Pom152 molecules are anchored to the outer rim of each spoke and their CTDs form a lumenal ring at the midline of the scaffold (7) (Fig. 1A). Each Pom152 has 10 immunoglobulin-like repeats (75, 76) in a CTD that overlaps with a neighbor in an anti-parallel manner to form a lumenal arch. In total, 8 arches create a sinusoidal ring (Fig. S11G, bottom half, in silver). The Pom152 CTD is connected to a transmembrane domain that crosses the pore membrane and an N-terminal cytoplasmic domain, which interacts with the spoke (7); Fig. 8D; thus, the Pom152 ring and inner ring are linked together. Crosslinking data suggests that Pom34 may associate with the transmembrane domain of Pom152 at the central 2-fold axis of the spoke (7).

The Pom152 ring in the isolated NPC has an outer contour length of ∼ 400 Å for each repeat, while dilation of the inner ring expands the ring into an octagonal profile with a repeat distance of ∼ 440 Å (Fig. S11G, bottom half). In addition, the expanded Pom152 ring is in close contract with the lumenal surface of the pore membrane (Fig. 5C-D). Integrative methods were used to create models for the lumenal rings based on crosslinking data, models of the Ig-like domains in Pom152 (76) and low resolution 3D density maps (Fig. S11G, top half). In each model, a ribbon is formed by two anti-parallel strands of Ig-like domains that are aligned roughly in a side-by-side manner (Fig. S11H). However, the plane of the two-stranded ribbon in the isolated NPC is tilted when viewed from above, whereas the ribbon is viewed nearly edge-on in the expanded conformation. In the in situ Pom152 ring, the two strands appear to slide relative to each other to compensate for the ∼ 40 Å increase in the contour repeat. The position, robust construction, and anchoring of the Pom152 ring to the spokes suggests that this belt-like feature may help limit the extent of radial expansion by the inner ring and pore membrane in the NPC.

## Discussion

In this work, a multi-pronged approach has been used to provide the most comprehensive picture of an NPC in any organism to date. This in turn, has produced novel insights into the architectural principles, evolutionary origins and structure-function relationships of the NPC.

### A layered organization and flexible connectors suggest a mechanism for reversible NPC assembly

We resolved the inner ring of the NPC at α-helical resolution and have identified 28 novel connectors from Nic96 NTDs that tie together the inner and outer domains of the yeast spoke. Our improved models support the idea that the NPC contains a layered organization of Nup folds that reflect its evolution. The outermost layer contains Nups that interact with the pore membrane, including Y-shaped Nup84 complexes in the outer rings and Nups in the membrane interacting layer of the inner ring. Many of these Nups share similar folds with the COPI and COPII families of clathrin-like vesicle coating proteins that probably merged to form a proto-NPC at an early point in its evolution (Field and Rout, 2019). The β-propellers in these Nups contain MBMs that help anchor the NPC to the pore membrane. Moving radially inwards, the next layer is comprised of two adaptin-like proteins, underscoring the common evolutionary origin of vesicle coating proteins and the NPC. These proteins share structural homology with nuclear transport factors. The innermost layer contains coiled-coil, heterotrimeric Nsp1 complexes that form two diagonal walls and provide anchor points to position N-terminal FG domains within the central transport path (Fig. 8A).

A key question in cell biology is “how can huge NPCs assemble and disassemble so quickly”?. We find that flexible connectors from Nic96 “tie together” multiple layers in the inner ring of the NPC (see wiring diagram, Fig. 8A), which suggests that they may play a crucial role in eukaryotes during assembly of new NPCs and during mitosis. Connectors in the NTD of Nic96 (and its homolog Nup93) are known targets for reversible phosphorylation in eukaryotes that have a rapid mitotic dis-assembly and re-assembly of the NPC, along with SLiMs of Nup53/Nup59, Nup100, Nup116, and Nup145N. (45, 46, 77). In brief, the disassembly process may be correlated with a phosphorylation-dependent “untying” of these connectors due to their extensive biochemical modification (see red stars for human sites mapped onto Nic96 NTDs, Fig. 8A). In addition, the extent of phosphorylation may determine the degree of dis-association of interacting Nups within the spoke. For example, human Nup93 can be isolated in two sub-complexes with known partners from the inner ring in mitotically-arrested cells. Thus, Nup93 interacts with the p62 complex along with Nup188/Nup205 and forms a second sub-complex with Nup188/Nup155 (78). In yeast, this would be equivalent to Nic96 interacting with the Nsp1 complex and Nup188/Nup192, while forming a second complex with Nup188 and Nup157/Nup170, which are homologs of Nup155. In actuality, the pattern of free Nups and thier complexes could vary between different cell types, tissues or organisms during mitotic disasembly.

NPC reassembly would involve a “re-tying” together of Nups within the spoke at the end of telophase, which may occur upon dephosphorylation (see (79)). In the simplest model, Nic96 CTDs may associate with Nup157 and Nup170 to create the membrane interacting layer at the pore membrane (79). This would provide anchor points for flexible Nic96 NTDs that connect the membrane interacting layer with the central adaptin-like layer and the inner layer, thereby guiding assembly and helping to stabilize the spoke. This process could use individual Nups, sub-complexes or a mixture of the components and a similar, stepwise assembly may occur for new NPCs. The presence of these flexible connectors may also allow the NPC to accommodate large changes in diameter and adapt to local stresses during transport and other processes.

### A link between nucleoporin assembly and NLS mediated transport

Our analysis of the inner ring at α-helical resolution suggests that spoke assembly may be dependent on interaction motifs that are similar to those found between - and β-transport factors (shown schematically in Fig. 8B). Nucleoporins and nuclear transport factors share common structural features and we now show that they share interaction mechanisms. Thus, Nup structural motifs mimic interactions of -adaptors with β-transport factors at several levels. First the α-helical solenoid of the Nic96 CTD is structurally similar to karyopherin (34). Second, Nup192 and Nup188 may share a common ancestor with β-karyopherins based on their structural homology (34, 80, 81). Third, interactions of the N-terminal domain of Nic96 with Nup192 and to a lesser degree with Nup188, strongly resemble the binding of the IBB domain of karyopherin with karyopherin β1 (Fig. 8B; (50), and are similar to other NLS – karyopherin β interactions (49). These observations have significant implications for the origin of nuclear transport; not only do transport factors and nucleoporins share structural similarities, but our higher resolution model now reveals that NPC assembly and transport factor interactions with cargo use similar mechanisms. This provides structural evidence for a common evolutionary origin of the NPC and its soluble transport machinery.

### NPCs are modular structures with multiple forms in yeast nuclei

The high degree of structural conservation in the spoke is in contrast to an expanding list of NPCs from different species with variable configurations for the outer rings. These differences are achieved by altering the copy number of Y-complexes, by the addition of side specific Nups and by repurposing Nups for alternate structural roles as linkers or structural elements in the outer rings (18). When taken together, these factors account for most of the differences in the size and mass of human and yeast NPCs (7, 17). A single outer ring in the yeast NPC consists of eight copies of the Y-shaped, seven-protein Nup84 complex arranged in a circular “head-to-tail” arrangement. We showed previously that two such rings are generally found in the yeast NPC, with one on the nuclear side and one on the cytoplasmic side (15, 23, 30, 32, 51).

In this study, we addressed a crucial question that has been asked in cell biology for decades, namely: are all NPCs in the same cell and nucleus fundamentally the same? Remarkably, the answer we find here is a clear: “No”. Thus, we show that nuclear double outer rings are present in a subset of yeast NPCs, while the majority of particles have single rings. This large scale pleomorphism contributes to three structural variants that coexist in the same nucleus and localize to distinct regions of the NE (summarized in Fig. 8C); These variants may foreshadow functional adaptations of the NPC in local regions of the nuclear periphery. This finding requires the NPC to be viewed as a modular assembly with a “Lego”-like ability to have major scaffold elements added, reorganized and/or removed. Differences in the number of outer rings could affect the overall rigidity of the NPC scaffold; this property may be adapted to the requirements of a particular cell type and organism or even a distinct region at the nuclear periphery. It also seems reasonable to suggest that these structural variants may represent functionally distinct NPCs that might be specialized in transporting specific cargos or in performing regulatory roles, such as transcriptional control or directing chromatin organization (2, 82). The additional nuclear outer ring may, for example, provide extra anchor sites for regulatory factors. We speculate that the outer rings have evolved for adaptability both within and between species (37). This variation may be germane to the large number of nuclear transport-related diseases and developmental disorders (5, 83, 84).

### Dynamic alterations of the NPC play a central role in transport

The NPC is a dynamic assembly with a range of effective diameters (28, 29, 58) and recent studies have revealed that the inner ring can dilate in cells to create a larger central channel (19, 23). Our analysis provides additional structural evidence for changes that occur in the inner ring, as NPCs transition from a constricted to a radially dilated conformation of the spokes. This radial movement of the spokes is coupled with a similar displacement of the pore membrane and alterations in the lumenal ring formed by Pom152 (see Fig. 8D); these changes may reflect a transition from a relaxed to a tense configuration of the inner ring. We also provide new insights into the roles of membrane protein interactions that anchor the inner ring within the membrane pore to stably position this transport machine. Spoke interactions with the pore membrane are mediated by Pom152, Pom34 and Ndc1; these interactions are 2-fold symmetric in the isolated NPC but may be altered locally during radial dilation of the inner ring. The lumenal Pom152 ring is also connected to the spokes and thus, may function like a belt to limit the extent of radial dilation of the inner ring, which in turn may modulate transport activity (Discussion).

Our comprehensive model of the in situ NPC reinforces the idea that local duplication and structural plasticity play critical roles in the assembly and function of this transport machine. The local 2-fold symmetry of the inner ring is broken by the outer rings and their associated components including the cytoplasmic Nup82 complex and the nuclear basket. The latter extends into the nucleoplasm and plays an important role in targeting and pre-processing of nuclear export complexes, including mRNPs, and may serve other unknown functions. For the first time, we mapped pairs of basket anchors which extend from the nuclear outer ring and show that these motifs appear to follow the structural theme of local duplication established for the NPC scaffold. Hence, the duplication of Nups or the presence of paralog pairs within a rotational unit of the NPC may extend to basket anchor complexes and thus, into the basket itself (Fig. 8E).

Finally, the two configurations of the inner ring display distinctive patterns for connections from Nsp1 complexes to FG domains in the central channel. This transition is associated with an approximate 2-fold increase in the volume of the central channel (see Fig. 8D), which may help explain changes in nuclear transport induced by mechanical forces (85–87). In both configurations, the central channel may be partitioned into distinct regions including a zone of connections and a zone of FG repeats. We suggest that the packing density of FG domains in the central channel must be altered during inner ring dilation, which will have important consequences for transport. Indeed, fundamental principles that govern dynamic FG domain organization remain to be uncovered in this “undiscovered” country.

## Acknowledgments

We thank members of the Chait, Villa and Rout laboratories for technical and intellectual support. D.S. is supported by a Damon Runyon Postdoctoral Fellowship (DRG-2364-19). V.J.M. was funded by an Ruth L. Kirschstein NRSA Postdoctoral Fellowship (F32 GM133096). This work was supported by NIH P41 GM109824 (B.T.C., M.P.R., A.S.), NIH R01 GM112108 (M.P.R.), NIH R01 GM45377 (C.W.A.), NIH P01 GM121203 and NIH R01 GM121443 (S.J. L.), NIH DP2 GM123494 (E.V.), NSF-1818129 (J.F-M.), the Pew Scholars program (E.V), NSF MRI DBI 1920374 (E.V.) and the Stowers Institute for Medical Research (S.L.J.). We acknowledge the use of the UC San Diego cryo-EM facility, which was built and equipped with funds from the University of California San Diego and an initial gift from Agouron Institute. We thank Prof. Ricardo Henriques’s Lab for LATEX template used for typesetting this document.

## Author Contributions

- C.W.A., J.F-M., I.N., D.S., S.S., K.S., C.Y. performed investigations.
- C.W.A., C.O., D.S., S.J.L., I.E., J.C.G., and J.U. performed analysis.
- C.W.A., J.F-M., D.S., J.M.V., S.L.J., S.J.L., A.S., B.T.C., I.E., E.V. and M.P.R. wrote the manuscript.
- C.W.A., E.V., S.J.L., A.S., S.L.J. B.T.C., and M.P.R. provided funding support.
- C.W.A, E.V., S.J.L., S.L.J., A.S., B.T.C. and M.P.R. supervised the project

## Methods

### Yeast strains and materials

Unless otherwise stated, all S. cerevisiae strains used in this study were grown at 30°C in YPD media (1% yeast extract, 2% bactopeptone, and 2% glucose).

### Immuno-purification of the endogenous S. cerevisiae NPC

Intact S. cerevisiae NPCs were isolated following the method described (7). To minimize local domain movements, purified NPCs were mildly cross-linked by the addition of DSS (DiSuccinimidylSuberate, Creative Molecules) to yield a final concentration of 0.5 mM and incubated for 30 min at 25°C with gentle agitation in a shaker (900 rpm). The reaction was then quenched by addition of ammonium bicarbonate to a final concentration of 50 mM. After Cysteine reduction, cross-linked samples were separated by NuPage SDS-PAGE (4-12%, Invitrogen) for quality control and concentration estimation. Gels were briefly stained by GelCode Blue Stain Reagent (Thermo Fisher Scientific) to enable the visualization of the protein complex. Aliquots of crosslinked NPCs (∼ 0.3 mg/ml) were flash frozen and stored at -80º C in small aliquots in buffer: 20 mM HEPES (pH7.5), 50 mM Potassium acetate, 20 mM NaCl, 2 mM MgCl2, 1 mM DTT, 0.1% Tween 20) supplemented with 10% glycerol. Samples were thawed only once prior to making grids for data collection.

### Chemical cross-linking and MS (CX-MS) analysis of affinity-purified yeast NPCs

NPCs were immuno-purified from Mlp1 tagged S. cerevisiae strains (7). After native elution, 1.0 mM disuccinimidyl suberate (DSS) was added and the sample was incubated at 25ºC for 40 min with shaking (1,200 rpm). The reaction was quenched by adding a final concentration of 50 mM freshly prepared ammonium bicarbonate and incubating for 20 min with shaking (1,200 rpm) at 25ºC. The sample (50 μg) was then concentrated and denatured at 98ºC for 5 min in a solubilization buffer (10% solution of 1-dodecyl-3-methylimidazolium chloride (C12-mim-Cl) in 50 mM ammonium bicarbonate, pH 8.0, 100 mM DTT). After denaturation, the sample was centrifuged at 21,130 g for 10 min and the supernatant was transferred to a 100 kDa MWCO ultrafiltration unit (MRCF0R100, Microcon). The sample was quickly spun at 1,000 g for 2 min and washed twice with 50 mM ammonium bicarbonate. After alkylation (50 mM iodoacetamide), the cross-linked NPC in-filter was digested by trypsin and lysC O/N at 37ºC. After proteolysis, the sample was recovered by centrifugation and peptides were fractionated into 10-12 fractions by using a stage tip self-packed with basic C18 resins (Dr. Masch GmbH). Fractionated samples were pooled prior to LC/MS analysis.

Desalted cross-link peptides were dissolved in the sample loading buffer (5% Methanol, 0.2% FA), separated with an automated nanoLC device (nLC1200, Thermo Fisher), and analyzed by an Orbitrap Q Exactive HFX (Pharma mode) mass spectrometer (Thermo Fisher) as previously described (88, 89). Briefly, peptides were loaded onto an analytical column (C18, 1.6 μm particle size, 100 Å pore size, 75μm×25cm; IonOpticks) and eluted using a 120-min liquid chromatography gradient. The flow rate was approximately 300 nl/min. The spray voltage was 1.7 kV. The QE HF-X instrument was operated in the data-dependent mode, where the top 10 most abundant ions (mass range 380 – 2,000, charge state4 - 8) were fragmented by high-energy collisional dissociation (HCD). The target resolution was 120,000 for MS and 15,000 for tandem MS (MS/MS) analyses. The quadrupole isolation window was 1.8Th; the maximum injection time for MS/MS was set at 200ms.

The raw data were searched with pLink2 (90). An initial MS1 search window of 5 Da was allowed to cover all isotopic peaks of the cross-linked peptides. The data were automatically filtered using a mass accuracy of MS1 10 ppm (parts per million) and MS2 20 ppm of the theoretical monoisotopic (A0) and other isotopic masses (A+1, A+2, A+3, and A+4) as specified in the software. Other search parameters included cysteine carbamidomethyl as a fixed modification and methionine oxidation as a variable modification. A maximum of two trypsin missed-cleavage sites was allowed. The initial search results were obtained using a default 5% false discovery rate (FDR) expected by the target-decoy search strategy. Spectra were manually verified to improve data quality (7, 71). Cross-linking data were analyzed and plotted with CX-Circos (http://cx-circos.net).

### Mapping Nic96-Nup188/192 interactions by in vitro reconstitution

Full length (FL) Nic96 along with C and N-terminal truncations were constructed and purified as previously described (7, 30). Affinity purified and natively eluted full length Nup188 and Nup192 (30) were incubated with the indicated affinity matrix-bound Nic96 complexes in binding buffer (20 mM Hepes-KOH pH 7.4, 50 mM potassium acetate, 20 mM sodium chloride, 2 mM magnesium chloride, 0.1% Tween-20, 1 mM DTT, 1/250 PIC) for 1 hour at 4ºC. After appropriate washes, the bound complexes were resolved by SDS-PAGE and stained with Coomassie R-250.

### Quantitative fluorescence imaging

Standard PCR techniques were used to generate strains expressing C-terminally tagged Nup fusion proteins. The outer ring component Nup84 was tagged with the GFP1-10 fragment of split-GFP, and plasmids expressing GFP11 reporters localized to the nucleoplasm (GFP11-mCherry-Pus1) or outer nuclear membrane/ER (GFP11-mCherry-Scs2-TM) were used to reconstitute GFP fluorescence and visualize rings on the nucleoplasmic and cytoplasmic faces of the NPC independently. Nic96-mTurquoise2 was used to detect and segment NPCs during analysis (see below).

Cells were grown in synthetic complete media lacking leucine (SC-leu; 6.7 g Yeast Nitrogen Base without amino acids, 2.0 g SC-Leu supplement mixture (Sunrise Scientific), 2% Glucose, pH 5.8) overnight at 30°C, back diluted and grown to an OD600 of 0.6-0.8 prior to harvesting. Cells were then pelleted for 3 min at 3000 x g and resuspended in 1 mL of 4% paraformaldehyde, 100 mM sucrose and fixed for 15 min at room temperature. Following three washes with 1 mL 1xPBS, the cell pellet was resuspended in 20 μL Prolong Diamond mounting media (Invitrogen) and slides were prepared by applying 3 μL of cells between slides and number 1.5 glass cover slips. Slides were cured overnight at room temperature and stored at 4 degrees until imaging.

Image stacks covering the entire nuclear volume were acquired with sub-Nyquist sampling and 0.126-0.220 nm step sizes on a Leica SP8 confocal microscope equipped with a HC PL APO CS2 100x, 1.40 NA objective. 458 nm, 496 nm and 561 nm lasers were used to excite mTurquosie2, rGFP and mCherry respectively, and emission for the corresponding channels were in the ranges of 463-495 nm, 505-550 nm and 566-660 nm. Pinhole size was set to 0.6 Airy unit to increase resolution, and raw images were automatically processed with default HyVolution settings. Individual nuclei were manually cropped from deconvoluted image stacks and analyzed with custom plugins written in ImageJ (NIH). Downstream analysis and data visualization was performed using R (v. 3.6.1) and RStudio (v. 1.1.442). All original data and custom code used for analysis can be accessed from the Stowers Original Data Repository at http://www.stowers.org/research/publications/libpb-1583. NPCs were detected in three dimensions using a “track max not mask” approach. Briefly, the brightest voxel in the image was found and a spheroid with a diameter of 12 pixels in x and y and 3 slices in z was masked around that voxel. This process was repeated until no voxels were left unmasked above 20% of the maximum intensity in the image. Next the average intensities of these presumptive NPCs were measured with a spheroid of diameter 8 pixels in x and y and 2.5 slices in z. Average intensity measurements for each image and channel were normalized to the average NPC intensity for the entire image. After normalization, foci with an intensity less than 0.35 in the Nic96-mTurqouise2 channel were removed to avoid poorly detected NPCs and background regions. For analysis of Nup84 levels with respect to Mlp1 intensity, NPCs were classified as either “high” or “low” based on whether the normalized Mlp1-mCherry intensity was above or below the mean Mlp1 intensity (1.0). The ratio of Nup84-rGFP to Nic96-mTurqouise2 was used for measurement purposes as it corrects for potential detection efficiency issues that are shared across fluorescence channels.

### Cryo-EM single particle data collection and movie processing

Sample grids were optimized to obtain a range of NPC views with a high loading; thus, particles were adsorbed on a carbon film overlayed onto Quantafoil 400 mesh carbon holey grids (2/2, hole diameter 2.4 uM). Grids were glow discharged followed by an extended floatation with carbon side down on 5 μL drops in a humid chamber. This was followed by 3 stepwise transfers onto 20 μL drops of blotting buffer (NPC buffer without glycerol) supplemented with 0.1% DeoxyBigChaps (wt/wt), while keeping the backside of the grids dry. Each grid was mounted in a Mark III Vitrobot at 10º C and 100% relative humidity, blot buffer was removed with a manual blot from the bottom edge of the grid through a side port in the humid chamber; an appropriate volume of blot buffer was pipetted immediately onto the grid, which was double blotted and plunge frozen in liquid ethane cooled by liquid N2.

Super-resolution movies were collected as LZW compressed tiffs (2 movies per hole, 4015 in total) using SerialEM (Mastronarde, 2005) on a Titan-Krios at 300 kV with a K2 Summit direct electron detector (Cryo-EM Core, University of Massachusetts Medical School). A physical pixel size of 2.66 Å (corrected magnification 37,651) was used to target the 6-9 Å resolution range with these particles (∼ 1100 Å diameter by 600 Å height). To check particle distribution and contamination, a script written by Dr. Chen Xu was used to automatically collect test images that covered ∼ 9 holes at 200 um defocus from each candidate grid mesh with the attached CCD. Medium magnification montages were then collected for good meshes at a much lower dose to minimize total exposure. Navigator parameters were set to have 6-7 groups of holes per mesh, with one focus spot per group using an in-house script. For the movies, 40 frames were collected with 0.5 second exposure/frame, a total dose of 40 e/Å2 and a calculated defocus range of -1.5 to -3.8 um with a K2 Summit direct electron detector in super-resolution counting mode (Gatan Inc.) with a Bioquantum GIF and 20 eV slit width. Image quality was monitored “on the fly” using a script (Dr. Xu), which does local motion correction with IMOD (91) on the imaging computer and creates power-pair jpegs with a motion-corrected micrograph image paired with the fitted power spectrum from CTFFIND4.

After data collection, movies were decompressed, gain corrected and aligned with Motioncor2 (v1.2.3) (92) to produce dose corrected micrographs, which were used to estimate per micrograph contrast transfer function (CTF) parameters with GCTF (93). Manual triage was done with Motioncor2/GCTF power-pairs using the “eye of Gnome” viewer (eog) and awk scripts to eliminate poor micrographs (3218 remained; ∼ 80%). After parameter optimization, NPCs were picked with Gautomatch (http://www.mrclmb.cam.ac.uk/kzhang/) using an image stack of equi-spaced projection views that were calculated from our tomographic model with C8 symmetry ((7); EMD-7321) using EMAN2 (e2project3d.py (94). For initial screening, particles were extracted in 310 × 310 boxes with a pixel size of 5.32 Å to remove outliers with 2D and 3D Maximum likelihood-based classification in RELION 2.0 (Scheres), which resulted in 26049 selected NPCs.

### Single particle processing

Spokes and co-axial rings- To evaluate the overall quality and angular coverage, the data set was processed initially with particles extracted at 5.32 Å/pixel. Refinements with C8 symmetry and a low pass filtered starting model from tomography (7), gave a structure with RELION that was virtually identical to the published map but with many more features (global resolution 24 Å). Subsequent refinements with imposed C8 symmetry used particle images after subtraction of the central transporter density and local soft masks that focused respectively on the inner ring and each of the two distal rings (ring of cobra hood features and the hook ring). This resulted in a composite 3D map with a local resolution range of 14.4 to 30 Å in which features of the double outer ring began to emerge.

For multi-body refinements with density subtraction, we started with particles at 5.32 Å/pixel and after test refinements in RELION 3.0 (95), re-centered NPCs were extracted at 2.66 Å/pixel in 620 × 620 boxes. In brief, we divided the processing into two sequential multi-body cycles to avoid memory limitations due to the requirement for multiple masks, coupled with the large box size of the particles (Fig. S3). The first cycle identified the best protomers and was centered on ∼ 1.5x subunits within the full particle in the original box, wherein a subunit is comprised of a spoke and distal ring “features” located on either surface. Following subunit extraction and re-centering within a smaller box, a second multi-body refinement step focused on three regions within the protomer. Refinements in which masks were centered on the spoke 2-fold axis were of noticeably higher quality than those centered on the 2-fold axis that passes between spokes. To start, we expanded particle line entries in the consensus C8 alignment star file for the full NPC, by duplicating every particle line entry 8x with a cyclic permutation of the first Euler angle in 45º increments (relion_particle_symmetry_expand). This 8-fold expanded particle star file and the original image stack were used to generate an expanded image stack with relion_image_handler to give ∼ 208K images. A C1 refinement was then done with the expanded particle stack to bring each protomer into a common alignment and 2D classification was used to eliminate bad subunits (tau=2).

For the first multi-body refinement step, we created a soft mask that covered the remaining protomers (∼ 6.5x) in the ring-like particle, along with the central transporter using a C8 symmetrized 3D map and then subtracted the density within this mask from the particle stack in each refinement iteration by using a zero alignment flag in the multi-body star file. After refinement, density subtracted image stack files for the entire ∼ 1.5x protomer were written out and used for focused 3D classification with local refinements to eliminate any remaining damaged subunits. A local C1 consensus refinement was then done for the re-centered and density subtracted protomers, which were subjected to a second round of multi-body refinement with three overlapping masks that covered the spoke and each outer ring. We then focused on the spoke and appropriate images were re-extracted and a local 3D classification with a soft mask was run (+/- 9 degrees) to further clean the data set. Refinements with C1 symmetry and a mask produced a map for the spoke at ∼8 Å resolution (local range 7.9-11Å).

NPCs were imaged on a thin carbon support film that dominates the CTF estimation and this value reflects a particles position along the optical axis relative to true focus. Hence, the median value of the CTF for the particle center is dependent on its orientation on the carbon film. For the NPC, this offset will become more significant at higher resolution. We created a script based on the geometry of the particles and the Euler angles (96) to calculate the z-offset (median defocus change) to each NPC center. An ambiguity in sign and thus, the direction of z-change was resolved by carrying out two separate 3D refinements with all per particle CTF values calculated with the same directional offset: the resolution of the final map was used as an indicator of the correct orientation. This resulted in a refined spoke 3D map with an improved local resolution range (corrected 6.6-11 Å), but the median FSC0.143 improved by only 0.1 Å with a larger mask. Per particle CTF corrections improved the best ordered, central 60-70% of the map, while more peripheral, flexible regions were slightly worse, accounting for the small global change in FSC0.143. This spoke map was 2-fold symmetrized and normalized in EMAN2 (e2proc3d.py –process=normalize.local), which allows densities across the map to be represented more equally at a single threshold.

A similar two-step multi-body approach with particles at 3.99 Å/pixel was used to process the outer ring with a cobra hood shape and revealed a double outer ring of Nup84 Y-complexes. We found that overlapping V-shaped regions from Nup120, Nup85 and Seh1 in two offset Nup84 complexes were combined to create the hood shape in the original 3D maps. In the second round of multi-body refinement, two masks were applied to the image stack with density subtracted transporter; the zero subtraction mask contained the inner ring, membrane density and hook ring density, while the second alignment mask covered ∼ 1.5x subunits in the double outer ring. After re-extraction of centered outer ring particles, 3D classification with tau 8 and local refinement in multiple steps with gradually improved masks identified particles with a double outer ring (∼ 20%) and gave a final 3D map with a global resolution of ∼ 11.3 Å using a generous mask.

Maps from final refinements for the spoke and double outer ring were cloned to create 8 copies and aligned within the best 3D density map of the full NPC in Chimera (97), rotated appropriately and added together to create complete rings. A visual inspection with docked models in Coot did not reveal anomalous features at subunit boundaries in the reassembled 3D maps at the current resolution. Final 3D maps were masked in EMAN2 and dusted in Chimera to remove noise. The co-axial ring with hook shaped units could not be improved further with a multi-body approach due to its flexibility. The structure determination for the inner ring and double outer rings is summarized in Fig. S3 and Table S1.

### Central transporter and spoke connections

The central transporter and spoke connections were analyzed separately; an image stack at 5.32 Å/pixel was used for a C8 refinement in RELION. To focus on the transporter, a 3D classification with 8 classes was done with the skip-align option and a mask which included the central transporter and a small region of the surrounding spoke. Two classes with low density were eliminated and the remaining 6 classes were grouped together, auto-refined and post-processed. To focus on connections between the spokes and transporter, a 3D classification with 6 classes was carried out with skip-align and a mask that eliminated the central region of the transporter, both distal rings and part of the membrane remnant. Particles in the two best classes were merged, auto-refined and post-processed with C8 symmetry. Unfortunately, signal for the lumenal Pom152 ring was too delocalized to identify a better ordered subset with 3D focused classification in the isolated NPC.

### Modeling the spoke and double outer ring

We started with an integrative model of the yeast spoke (PDB-DEV_00000012) that was based on ∼ 25 crystal structures (see (7)) and docked 28 Nups within the spoke 3D density map as rigid bodies in Chimera. We then used manual modeling in Coot (98), Chimera, Phenix realspace_refine (99) and molecular dynamics flexible fitting (MDFF; (59)) to create a model based on a majority of the ∼ 715 α-helices within the spoke. We included flexible connectors from N-termini of four Nic96 molecules and 2 orphan helices to give 26 novel α-helices and 4 strands, along with tail domains with altered conformations from six Nups (two copies each of Nups 192, 188 and 170); the full spoke model contains ∼ 15,600 residues in the ordered domains (∼ 1.69 MDa). A novel Integrative threading was used to position sequences of Nic96 N-termini within the orphans (see below, (43)), which was followed by an additional round of refinements to create a final model (PDB 7N85; EMD-24232). Nup domains in the structure have undergone local movements and rearrangements with an average Cα rmsd of ∼ 8Å, when compared to an earlier integrative model of the spoke (7), with larger changes at Nup contacts and due to rearrangements of the Nsp1 complex. For the double outer ring, 14 Nups were docked into the repeating unit (∼ 1.1 MDa) using yeast homology models based on 5 crystal structures (32) that were split into 22 domains with a total of ∼ 10,000 residues. The improved resolution allowed us to model complete paths for two Y-complexes within the double ring along with a copy of Nup188/192 and a putative basket anchor in a rod-like orphan density. Initial models were subjected to manual remodeling in Coot, Chimera, and refinements with Phenix realspace_refine (while ensuring that perturbations of the β-propellers did not occur from over-refinement). Final fitting steps used MDFF: a gscale value of 0.6 was used for medium resolution models (7-10Å), while flexible fitting into lower resolution maps used values of 0.15-0.3. Typical restraints were used for protein geometry in refining the double ring model (PDB 7N84, EMD-24231).

### Mapping cross-links to the NPC structures

A total of 2,053 DSS and EDC crosslinks were obtained previously for the isolated NPC (7). To assess whether the crosslinks agree with NPC structure, these crosslinks were mapped on to the cryo-EM structure by determining the Cα–Cα distance of crosslinked residue pairs. A DSS or ECD crosslink was classified as satisfied by the structure if the corresponding Cα–Cα distance is less than 35 or 30 Å, respectively (Fig. S8B-E).

### Initial model for 30 orphan densities within the spoke

A total of 30 orphan densities were identified in the spoke that likely arise from N-termini of Nic96 and Nup53/Nup59, functional orthologs of CtNup53 and possibly from other Nups. We identified 20 orphans combined in the two diagonal walls. In addition, 10 orphan helices are associated with two copies Nup192 and two copies of Nup188. Physical, biochemical and structural restraints were used to create an initial model for connectors in the orphan densities. Previous work showed that a large connector is present in the N-terminus of CtNup53 which interacts with the NTD of Nup192 (20). We observed a long helix associated with the NTD of Nup192 that may be attributed to Nup59 because this protein interacts with Nup170 and Nup192 in pullout experiments and thus, behaves like CtNup53. This left 28 orphans in a spoke that may be contributed by connectors from four Nic96 N-termini with 10 in each diagonal wall.

The Nic96 N-terminus contains 204 residues, so the sum of orphan structural elements + linker residues in each modeled chain should not exceed this value. Moreover, the distance between diagonal walls is too great to allow a single Nic96 N-terminus to be located in both walls. Previous work identified two conserved sequence motifs in the N-termini of Nic96 and its homolog, Nup93 that interact with Nsp1 and p62 complexes, respectively and with the C-terminal tails of Nup188 and Nup192/Nup205 (20). The 5 orphans in each diagonal half-wall are present in two groups with linker density visible at a lower threshold for some pairs: a group of three structural elements (two helices and a strand) and a group of two helices. After manual docking of poly-Ala α-helices and strands in Chimera and Coot, we found that the topology of the group of 3 structural elements is similar to the helix-strand-helix motif of a Nic96 SLiM observed in a crystal structure with a Ct-Nsp1 heterotrimer ((33); PDB 5CWS; Fig. S8F). Structure prediction of the Nic96 N-terminus also suggested that two helices may be positioned between the group of 3 structural elements and a pair of helices located near the end of the NTD that are bound to Nup192 or Nup188. Thus, the final pair of connectors from the innermost copy of Nic96 may interact with a CTD of Nup192 while connectors from the outer copy of Nic96 probably bind to the Nup188 CTD. Manually docked helices and strands without linkers were refined with Phenix realspace_refine (99) and these coordinates were used as input to an integrated threading approach to test all possible structure element orientations and sequences from Nups as connectors within the orphan densities; this analysis did not use the crystal structure of the Nic96 SLiM (33).

### Integrative threading of Nup sequences into spoke orphan structure elements

A novel integrative threading approach (7, 43, 100–102) was applied to determine the protein, its copy identifier, and the sequence of orphan structure elements (SE) in the inner ring. All possible arrangements derived from the spoke C2-symmetry were considered, but only one resulted in a model that sufficiently satisfied the input data: this option splits the 30 orphan into two symmetric subsets of 15 orphan densities each. Integrative modeling proceeded through the standard four stages, as follows:

Stage 1 - Gathering information- The data used for integrative structure modeling is summarized in Table S2.

Stage 2 - Representing subunits and translating data into spatial restraints We represent the NPC model using a threading representation. The representation defines the degrees of freedom of the model whose values are fit to input information.

Threading representation

We first identified a total of 30 orphan densities in the inner-ring cryo-EM density map. These densities most likely correspond to distinct SEs, including α-helices and β-strands. A poly-Ala sequence of suitable length was manually fit into each rod-like orphan density, followed by refinement using Phenix realspace_refine as described. Each SE is defined by four degrees of freedom that map it to specific residues in the sequence: 1) Start Residue, defining the first residue in the sequence to which the SE is mapped; 2) Length, the number of residues in the sequence overlapping with the SE; 3) Offset, the index of the first SE residue that is assigned to the sequence; and 4) Polarity, which defines the direction of the mapping between the SE and the sequence (Fig. S7C). Coarse-grained structural representation of the NPC scaffold: If an atomic structure of a protein segment is present in the isolated NPC structure described above, it was represented by the corresponding rigid body configurations of a fine- and coarse-grained representation of 1- and 10-residue beads, respectively (Table S2, (7), references therein).

Scoring function for threading representation: Alternative models are ranked via a scoring function. The scoring function is a sum of terms, each one of which restrains some aspect of the model based on a subset of input information (e.g., stereochemistry, cross-links, sequences). We impose three scoring terms that restrain the four degrees of freedom for each of the 30 SEs. First, the secondary structure restraint is computed by comparing the secondary structure assignment in the model with the secondary structure propensity predicted from the input sequence. Second, the loop end-to-end distance (i.e., C-to N-termini distance between two sequentially adjacent SEs) restraint relies on the relationship between the model loop length and it’s end-to-end distance as derived from the CATH 3D structural database. Third, the SE crosslinking restraint corresponds to a harmonic upper-bound on the distance spanned by two cross-linked residues. When one or both crosslinked residue(s) have undefined position(s), we compute the distance from inferred localization volumes.

Stage 3 - Enumeration of threading solutions to produce an ensemble of structures that satisfy the restraints. Sequences from appropriate Nups were threaded through the backbone of the 30 SEs in a spoke. We improved the efficiency of threading by explicitly considering the C8- and C2-symmetries of the inner ring (Fig. 1).

Symmetry

We defined the coordinate system such that the C2-symmetry is imposed simply by cloning a bead in the C2-symmetry unit at (x, y, z) to (x, y,z) (equivalent to a rotation of 180° around the × axis). In addition, a second 2-fold axis was considered for sequence threading of the orphan densities (Fig. S7B). Based on symmetry, we grouped the SEs into five groups: 1) SEs 1-3 that bind to the Nsp1/Nup49/Nup57 coiled-coils, 2) SEs 4 and 5 that localize between copies of the Nsp1/Nup49/Nup57 coiled-coils, 3) SE 6 that binds to the Nup188 CTD, 4) SE 6 that binds to Nup192 CTD, and 5) SE 7 that binds the Nup192 NTD (Fig. S7B). All possible symmetry arrangements derived from the spoke C2-symmetry were considered, such that 30 orphan SE were split into two symmetric subsets of 15 SEs each.

Threading enumeration: We constructed all possible sequence assignments of a subset of sequences to positions of the SEs by enumerating of the threading degrees of freedom. The subset of sequences included Nic96 (residues 1-205), Nup53 (residues 1-247), Nup59 (residues 1-265), Nup100 (residues 551-815), Nup116 (residues 751-965), and Nup145N (residues 201-458). These segments correspond to regions that have previously been modeled as flexible beads and do not contain FG repeats (Table S2) (7). The enumeration begins by assigning a protein identity and copy number to each of the five SE groups. Next, the scoring function is evaluated for all possible assignments of the four SE keys described above. For each SE group, identical keys are applied to all symmetry copies of the group (Fig. S7).

Stage 4 - Analyzing and validating the model ensemble Model validation follows five steps (43, 103, 104): (i) selection of models for validation; (ii) estimation of sampling precision; (iii) estimation of model precision, (iv) quantification of the match between the model and information used to compute it, and (v) quantification of the match between the model and information not used in the computation. Next, we discuss each one of these validations separately.

(i) Model selection. Models were filtered to find models that sufficiently satisfy the input information; a secondary structure restraint is satisfied if the RaptorXProperties score for the sequence-to-SE mapping is higher than 0.6; a loop end-to-end distance restraint is satisfied if the distance between the termini of two sequential SEs is violated by less than 3 times the standard deviation parameter used to evaluate the restraint; similarly, a crosslink restraint is also satisfied when an evaluated distance is violated by less than 3 standard deviations.

(ii) Sampling precision. Because threading solutions were enumerated, no sampling precision needed to be calculated.

(iii) Model precision. Model precision (uncertainty) is defined by the variation among the filtered models; the precision of each SE in each cluster was assessed by computing the standard deviation of its starting residue over all models in the cluster.

(iv) Information used in modeling. By selection, filtered models satisfy the input information.

(v) Information not used in modeling. Perhaps the most convincing test of a model is by comparing it to information not used in the computation (a generalization of cross-validation). We compared the structure of the Nic96 NTD bound to the Nsp1/Nup49/Nup57 coiled-coil domain to the X-ray structure of the same sub-complex from Chaetomium thermophilum (33) (Fig. S8F). Additionally, the mapping of known Nic96 NTD phosphorylation sites from different species to the integrative threading model rationalized how these site modifications might regulate NPC disassembly and reassembly (Fig. S8A), further increasing our confidence in the modeling. Phosphorylation sites were retrieved from the PhosphoSitePlus server (105), a manually curated database of experimentally confirmed phosphorylation sites. Finally, the models are consistent with biochemical considerations as described in the initial modeling section (20).

### In situ tomography of S. cerevisiae NPCs-data collection and pre-processing

W303 yeast cells were harvested during log-phase growth and diluted to an 0.8 × 107 cells/mL in YPD media. Five μL of this diluted sample was applied to glow-discharged 200-mesh, Quantafoil R2/1 grids (Electron Microscopy Sciences), excess media was manually blotted from the grid back side (opposite to the carbon substrate where the cells were deposited) and the grid was plunge frozen in a liquid ethane-propane mixture (50/50 volume, Airgas) using a custom-built vitrification device (Max Planck Institute for Biochemistry, Munich). Frozen grids were clipped into AutoGrids with a notch to allow milling at shallow grazing angles (Thermo Fisher Scientific). The cryogenic, FIB milling was performed in an Aquilos DualBeam (Thermo Fisher Scientific). Clusters of a few yeast cells (Fig. S9) were marked for milling at an angle of ∼11-14° (corresponding to stage tilts of 18-21°). Lamellae of ~11-18 μm in width and thicknesses ranging from 120 to 180 nm were prepared using rectangular milling patterns as noted previously (106).

Grids containing lamellae were loaded into a Titan Krios G3 (Thermo Fisher Scientific) operated at 300 kV. First, lamellae were identified in low magnification TEM images and their coordinates were stored. At a nominal magnification of ∼ 2250X or 3600X, the eucentricity at each of the lamellae coordinates was determined and another set of images at a nominal magnification of 8700X was used to generate montages of each lamella in SerialEM (107). The nuclear membrane and NPCs were identified and marked in these montages as targets for data acquisition. Tilt series were acquired on the Krios equipped with a post-column energy filter and a K2 Summit direct detector in counting and dose fractionation modes (Gatan). Typical tilt-series parameters were as follows: tilt range: ∼ ± 60°, nominal magnification 42000X, tilt increment 3°(4-5° for some samples), pixel size of 3.43 Å and an effective defocus range of -2 to -11 μm. All image-acquisition was done using SerialEM software (107) (Table S3). Frames of the tilt-series movies were motion-corrected using Motioncorr2 (92). Non-dose weighted and motion-corrected image stacks were imported into EMAN2 for subsequent processing, including CTF estimation, reconstruction, sub-tomogram, and sub-tilt refinement (108).

### Sub-tomographic analysis of in situ NPCs

Data processing, beginning with collected tilt series, followed the standard sub-tomogram analysis workflow in EMAN2.9 (108). Briefly, a few tilt series were automatically reconstructed at 4x down-sampling (1k ×x 1k × 512). Automatic CTF processing was used to check handedness and determine the correct choice of tilt-axis. Once the tilt axis was determined, all tilt series were automatically aligned and reconstructed at 4x down-sampling using e2tomogram.py followed by automatic, tilt-aware CTF determination. NPCs were manually selected from 930 tomograms, avoiding partial NPCs that were cut by the FIB milling, yielding 518 particles. Complete NPCs were re-extracted and reconstructed from the raw tilt series at 2x down-sampling (6.9 Å/pix). The relatively low yield resulted from a compromise between preparing thin lamella to increase data quality and the goal of including complete NPCs, which are comparable in diameter to the optimal lamella thickness.

Numerous alignment runs were completed on whole NPCs and on individual protomers with a variety of different focused masks to identify regions of high variability and to determine the optimal approach for final refinement. In the end, we used a single mask centered on a protomer without additional focused refinements. For the final refinement NPCs were iteratively and globally aligned. The final alignment soft mask covered the spoke ring extending into the membrane and included parts of the cytoplasmic and nucleoplasm rings. Then, 8-fold symmetry was imposed, considering the clear symmetry in the scaffold of the NPC. Based on the alignment, NPCs were mapped back into the tomograms. A few particles (24) were found to lie outside the plane of the nuclear envelope, and thus were considered misaligned and were discarded. During alignment, other particles were placed with the cytoplasm and nucleoplasm inverted; these particles (91, ∼ 18%) were flipped. This was followed by another round of whole-NPC alignment with rotation and translation constrained to 10 pixels and 24 degrees of rotation in all three Euler angles.

Final NPC orientations from this refinement were used to extract 8 protomers from each NPC (C8 symmetry expansion), each in a box large enough to contain all of the density in a 90 degree wedge (one protomer plus two halves), centered on a spoke and extending just past the center of the NPC. These 3656 particles, which were pre-aligned based on the overall NPC alignment, were then iteratively refined using a single alignment soft mask focused on the spoke and outer rings. This protomer alignment was constrained to a maximum of 5 degrees of rotation and 8 pixels of translation per iteration with a classkeep of 0.8 (an EMAN parameter to limit the fraction of particles to include in the average for the next iteration). The final average was performed under a simple spherical mask extending well into the membrane. Per-particle and per-tilt CTF refinement did not yield useful results for the NPC, presumably due to the inherent flexibility and the small number of particles.

Local resolution was estimated using the EMAN2 implementation of mFSC (109) following the ‘gold standard’ refinements and indicates a range of 30-50 Å in the spoke, outer rings and Pom152 domains. Maps were locally Wiener filtered as part of the mFSC calculation, which computes an FSC curve for every voxel in the map. Reliable visual detail (comparison of even/odd maps) matches or exceeds that of EMD10198 for the yeast NPC, which claims 25 Å resolution. We hypothesize that this difference is due to masking used in the earlier structure, but cannot assess this as even and odd maps were not deposited. Furthermore, we provide a direct comparison of our tomographic 3D map with selected views of EMD10198 (Fig. S10C-D). As reported in this work, additional peripheral features are clearly resolved in our 3D map including: connections from Nsp1 complexes in the spoke and from the Nup82 complex, along with orphan densities and basket anchor complexes on the nuclear outer ring, improved Pom152 ring density and spoke contact sites on the pore membrane. These differences may arise from masking steps required to divide the protomer into three overlapping regions for separate alignments and from local filtering and reassembly of the C8 protomer that occurred in previous work (23), whereas we used a strategy that aligns the entire protomer within a wedge containing two subunits. Alternatively, the amplitudes of Fourier components in this resolution range may be weak; thus, resulting 3D maps may not differ significantly in this resolution range. In fact, some combination of these factors may relate to the observed differences in estimated resolution.

### Refinement of the Pom152 ring from the in-situ NPC

The NPC is an intrinsically flexible assembly, and thus refinements of one segment may smear out the density of other segments. To refine the Pom152 ring (lumenal ring), the 3656 C8 protomers, as described above (pre-aligned based on the overall NPC alignment), were iteratively refined using a soft alignment mask that focused on the double nuclear membrane region. For these iterative refinements, the rotation and translation of the pre-aligned protomer particles were constrained to 8 pixels and 5 degrees of rotation in all three Euler angles. The final average was performed under a simple soft mask that covered the entire asymmetric subunit of the protomer and extended into the membrane regions. Composite maps and modeling of the in situ NPC

The final 3D density map for the in situ NPC was used to create models for spokes in the inner ring and for Nup84 Y-complexes in the outer rings, using methods already described. Additional components were docked manually in Chimera including: the Nup82 complex with 3 pairs of Dyn2 dimers bound to sequential DID repeats of Nup159 in the cytoplasmic outer ring, connections from Nsp1 in the Nup82 complex and from Nsp1 complexes in the inner ring that extend into the central channel, which used rods as linear markers, along with 14 2-helix coiled coil segments in basket anchor densities that emanate from each Y-complex in the nuclear outer ring. Docked models were cloned to form complete rings and used to segment the map by zoning in Chimera with a 20 Å radius to capture additional, nearest neighbor densities and these mini-maps were used for figures. SEGGER (110) was used to segment the Pom152 ring prior to integrative structure modeling. To compare maps of the isolated and in situ NPCs segmented inner rings were used for an initial alignment: the inner ring from the isolated NPC was scaled, low pass filtered and boxed in EMAN2.1 to give a comparable resolution with the same pixel and box size as the inner ring in the in situ 3D map. Both inner rings were then aligned for D8 symmetry (which is approximate for the in situ inner ring) with e2symsearch3D.py, and the spoke 2-fold along the x-axis. The 8-fold symmetrized, inner ring for the in situ NPC was then aligned to the D8 symmetrized map from this feature to create mutually-centered 3D maps for the two configurations. Outer rings were then placed in the correct orientation relative to their respective, aligned inner ring maps in Chimera.

### Integrative modeling of the lumenal Pom152 ring

Integrative modeling of the Pom152 ring in isolated and in situ NPCs proceeded through the standard four stages as described above (7, 100–102). The data used for integrative structure modeling of the Pom152 rings is summarized in Supplementary Table S2. Each the 10 Pom152 Ig-like domains was represented as a rigid body using a 1-residueper-bead representation; all other regions were represented as flexible strings of beads (Table S2). We modeled 6 copies of Pom152 spanning 3 spokes. The scoring terms that restrain the spatial degrees of freedom included: first, cross-link restraints based on total of 1425 DSS and EDC. The same set of cross-links was used for modeling the isolated and in situ Pom152 ring because only small conformational changes are expected (111). Second, the EM density restraint corresponding to a cross-correlation between the Gaussian Mixture Model (GMM) representation of the Pom152 subunits and the GMM representation of the cryo-EM or FIB-milled cryo-ET density maps for the lumenal ring (i.e., EM maps; Table S2) (112); third, transmembrane domain restraints that localize the coarse beads in the predicted trans-membrane domains (Pom152111–200) within the pore membrane (7); fourth, Pom152 peri-nuclear restraints applied to the CTD of Pom152 (residues 201–1337). Finally, excluded volume and sequence connectivity restraints were applied to all components (7). Structural models were computed using Replica Exchange Gibbs sampling, based on the Metropolis Monte Carlo (MC) algorithm (71, 113). Each MC step consisted of a series of random transformations (i.e., rotations and/or translations) of the flexible beads and rigid bodies (Table S2). We improved the efficiency of structural sampling by explicitly considering the C8- and C2-symmetries of the inner ring. Analysis and validation of the structural models of the Pom152 rings followed the previously published five steps (114) and results from this validation are summarized in (Table S2). The integrative structure modeling protocol (i.e., stages 2, 3, and 4) was scripted using the Python Modeling Interface (PMI) package and our open-source Integrative Modeling Platform (IMP) package (102), version 2.8 (https://integrativemodeling.org). Files containing input data, scripts, and output results are available at https://github.com/integrativemodeling/NPC_3.0 and in the worldwide Protein Data Bank (wwPDB) PDB-Dev repository for integrative structures (pdb-dev.wwpdb.org).

## Bibliography

1. Maximiliano A D’Angelo, Daniel J Anderson, Erin Richard, and Martin W Hetzer. Nuclear pores form de novo from both sides of the nuclear envelope, 2006.

2. J Fernandez-Martinez and M P Rout. One Ring to Rule them All? Structural and Functional Diversity in the Nuclear Pore Complex. Trends Biochem Sci, 46(7):595–607, 2021. doi: 10.1016/j.tibs.2021.01.003.

3. T M Kuhn and M Capelson. Nuclear Pore Proteins in Regulation of Chromatin State. Cells, 8(11):1414, 2019. doi: 10.3390/cells8111414.

4. C Ptak, J D Aitchison, and R W Wozniak. The multifunctional nuclear pore complex: a platform for controlling gene expression. Curr Opin Cell Biol, 28C:46–53, 2014. doi: 10.1016/j.ceb.2014.02.001.

5. R Juhlen and B Fahrenkrog. Moonlighting nuclear pore proteins: tissue-specific nucleoporin function in health and disease. Histochem Cell Biol, 150(6):593–605, 2018. doi: 10.1007/s00418-018-1748-8.

6. Dan N Simon and Michael P Rout. Cancer and the nuclear pore complex. Advances in experimental medicine and biology, 773:285–307, 2014. doi: 10.1007/978-1-4899-8032-8_13.

7. S J Kim, J Fernandez-Martinez, I Nudelman, Y Shi, W Zhang, B Raveh, T Herricks, B D Slaughter, J A Hogan, P Upla, I E Chemmama, R Pellarin, I Echeverria, M Shivaraju, A S Chaudhury, J Wang, R Williams, J R Unruh, C H Greenberg, E Y Jacobs, Z Yu, M J de la Cruz, R Mironska, D L Stokes, J D Aitchison, M F Jarrold, J L Gerton, S J Ludtke, C W Akey, B T Chait, A Sali, and M P Rout. Integrative structure and functional anatomy of a nuclear pore complex. Nature, 555(7697):475–482, 2018. doi: 10.1038/nature26003.

8. A Ori, N Banterle, M Iskar, A Andres-Pons, C Escher, H Khanh Bui, L Sparks, V Solis-Mezarino, O Rinner, P Bork, E A Lemke, and M Beck. Cell type-specific nuclear pores: a case in point for context-dependent stoichiometry of molecular machines. Mol Syst Biol, 9:648, 2013. doi: 10.1038/msb.2013.4.

9. Javier Fernandez-Martinez, Seung Joong Kim, Yi Shi, Paula Upla, Riccardo Pellarin, Michael Gagnon, Ilan E Chemmama, Junjie Wang, Ilona Nudelman, Wenzhu Zhang, Rosemary Williams, William J Rice, David L Stokes, Daniel Zenklusen, Brian T Chait, Andrej Sali, and Michael P Rout. Structure and Function of the Nuclear Pore Complex Cytoplasmic mRNA Export Platform. Cell, 167(5):1215–1228.e25, nov 2016. doi: 10.1016/j.cell.2016.10.028.

10. Monika Gaik, Dirk Flemming, Alexander von Appen, Panagiotis Kastritis, Norbert Mücke, Jessica Fischer, Philipp Stelter, Alessandro Ori, Khanh Huy Bui, Jochen Baßler, Elisar Barbar, Martin Beck, and Ed Hurt. Structural basis for assembly and function of the Nup82 complex in the nuclear pore scaffold, 2015.

11. J Yamada, J L Phillips, S Patel, G Goldfien, A Calestagne-Morelli, H Huang, R Reza, J Acheson, V V Krishnan, S Newsam, A Gopinathan, E Y Lau, M E Colvin, V N Uversky, and M F Rexach. A bimodal distribution of two distinct categories of intrinsically disordered structures with separate functions in FG nucleoporins. Mol Cell Proteomics, 9(10):2205–2224, 2010. doi: 10.1074/mcp.M000035-MCP201.

12. K E Knockenhauer and T U Schwartz. The Nuclear Pore Complex as a Flexible and Dynamic Gate. Cell, 164(6):1162–1171, 2016. doi: 10.1016/j.cell.2016.01.034.

13. G Paci, J Caria, and E A Lemke. Cargo transport through the nuclear pore complex at a glance. J Cell Sci, 134(2), 2021. doi: 10.1242/jcs.247874.

14. Susan R Wente and Michael P Rout. The nuclear pore complex and nuclear transport. Cold Spring Harbor perspectives in biology, 2(10):a000562, oct 2010. doi: 10.1101/cshperspect.a000562.

15. Frank Alber, Svetlana Dokudovskaya, Liesbeth M Veenhoff, Wenzhu Zhang, Julia Kipper, Damien Devos, Adisetyantari Suprapto, Orit Karni-Schmidt, Rosemary Williams, Brian T Chait, Michael P Rout, and Andrej Sali. Determining the architectures of macromolecular assemblies, 2007.

16. Khanh Huy Bui, Alexander von Appen, Amanda L. DiGuilio, Alessandro Ori, Lenore Sparks, Marie-Therese Mackmull, Thomas Bock, Wim Hagen, Amparo Andrés-Pons, Joseph S. Glavy, and Martin Beck. Integrated Structural Analysis of the Human Nuclear Pore Complex Scaffold. Cell, 155(6):1233–1243, 2013. doi: 10.1016/j.cell.2013.10.055.

17. Jan Kosinski, Shyamal Mosalaganti, Alexander von Appen, Roman Teimer, Amanda L DiGuilio, William Wan, Khanh Huy Bui, Wim J H Hagen, John A G Briggs, Joseph S Glavy, Ed Hurt, and Martin Beck. Molecular architecture of the inner ring scaffold of the human nuclear pore complex., 2016.

18. A von Appen, J Kosinski, L Sparks, A Ori, A L DiGuilio, B Vollmer, M T Mackmull, N Banterle, L Parca, P Kastritis, K Buczak, S Mosalaganti, W Hagen, A Andres-Pons, E A Lemke, P Bork, W Antonin, J S Glavy, K H Bui, and M Beck. In situ structural analysis of the human nuclear pore complex. Nature, 526(7571):140–143, 2015. doi: 10.1038/nature15381.

19. Shyamal Mosalaganti, Jan Kosinski, Sahradha Albert, Miroslava Schaffer, Daniela Strenkert, Patrice A Salomé, Sabeeha S Merchant, Jürgen M Plitzko, Wolfgang Baumeister, Benjamin D Engel, and Martin Beck. In situ architecture of the algal nuclear pore complex. Nature communications, 9(1):2361, 2018. doi: 10.1038/s41467-018-04739-y.

20. D H Lin, T Stuwe, S Schilbach, E J Rundlet, T Perriches, G Mobbs, Y Fan, K Thierbach, F M Huber, L N Collins, A M Davenport, Y E Jeon, and A Hoelz. Architecture of the symmetric core of the nuclear pore. Science, 352(6283):aaf1015, 2016. doi: 10.1126/science.aaf1015.

21. G Huang, Y Zhang, X Zhu, C Zeng, Q Wang, Q Zhou, Q Tao, M Liu, J Lei, C Yan, and Y Shi. Structure of the cytoplasmic ring of the Xenopus laevis nuclear pore complex by cryo-electron microscopy single particle analysis. Cell Res, 30(6):520–531, 2020. doi: 10.1038/s41422-020-0319-4.

22. Y Zhang, S Li, C Zeng, G Huang, X Zhu, Q Wang, K Wang, Q Zhou, C Yan, W Zhang, G Yang, M Liu, Q Tao, J Lei, and Y Shi. Molecular architecture of the luminal ring of the Xenopus laevis nuclear pore complex. Cell Res, 30(6):532–540, 2020. doi: 10.1038/s41422-020-0320-y.

23. M Allegretti, C E Zimmerli, V Rantos, F Wilfling, P Ronchi, H K H Fung, C W Lee, W Hagen, B Turonova, K Karius, M Bormel, X Zhang, C W Muller, Y Schwab, J Mahamid, B Pfander, J Kosinski, and M Beck. In-cell architecture of the nuclear pore and snapshots of its turnover. Nature, 586(7831):796–800, 2020. doi: 10.1038/s41586-020-2670-5.

24. D Devos, S Dokudovskaya, F Alber, R Williams, B T Chait, A Sali, and M P Rout. Components of coated vesicles and nuclear pore complexes share a common molecular architecture. PLoS Biol, 2(12):e380, 2004. doi: 10.1371/journal.pbio.0020380.

25. S M Bailer, C Balduf, and E Hurt. The Nsp1p carboxy-terminal domain is organized into functionally distinct coiled-coil regions required for assembly of nucleoporin subcomplexes and nucleocytoplasmic transport, 2001.

26. D R Finlay, E Meier, P Bradley, J Horecka, and D J Forbes. A complex of nuclear pore proteins required for pore function. J Cell Biol, 114(1):169–183, 1991. doi: 10.1083/jcb.114.1.169.

27. M Fornerod, J van Deursen, S van Baal, A Reynolds, D Davis, K G Murti, J Fransen, and G Grosveld. The human homologue of yeast CRM1 is in a dynamic subcomplex with CAN/Nup214 and a novel nuclear pore component Nup88. EMBO J, 16(4):807–816, 1997. doi: 10.1093/emboj/16.4.807.

28. J. Mahamid, S. Pfeffer, M. Schaffer, E. Villa, R. Danev, L. Kuhn Cuellar, F. Forster, A. A. Hyman, J. M. Plitzko, and W. Baumeister. Visualizing the molecular sociology at the HeLa cell nuclear periphery. Science, 351(6276):969–972, feb 2016. doi: 10.1126/science.aad8857.

29. C M Feldherr and D Akin. Variations in signal-mediated nuclear transport during the cell cycle in BALB/c 3T3 cells. Exp Cell Res, 215(1):206–210, 1994. doi: 10.1006/excr.1994.1333.

30. Javier Fernandez-Martinez, Jeremy Phillips, Matthew D Sekedat, Ruben Diaz-Avalos, Javier Velázquez-Muriel, Josef D Franke, Rosemary Williams, David L Stokes, Brian T Chait, Andrej Sali, and Michael P Rout. Structure-function mapping of a heptameric module in the nuclear pore complex., 2012.

31. S Siniossoglou, C Wimmer, M Rieger, V Doye, H Tekotte, C Weise, S Emig, A Segref, and E C Hurt. A novel complex of nucleoporins, which includes Sec13p and a Sec13p homolog, is essential for normal nuclear pores, 1996.

32. K Kelley, K E Knockenhauer, G Kabachinski, and T U Schwartz. Atomic structure of the Y complex of the nuclear pore. Nat Struct Mol Biol, 22(5):425–431, 2015. doi: 10.1038/nsmb.2998.

33. Tobias Stuwe, Christopher J Bley, Karsten Thierbach, Stefan Petrovic, Sandra Schilbach, Daniel J Mayo, Thibaud Perriches, Emily J Rundlet, Young E Jeon, Leslie N Collins, Ferdinand M Huber, Daniel H Lin, Marcin Paduch, Akiko Koide, Vincent Lu, Jessica Fischer, Ed Hurt, Shohei Koide, Anthony A Kossiakoff, and André Hoelz. Architecture of the fungal nuclear pore inner ring complex., 2015.

34. Parthasarathy Sampathkumar, Seung Joong Kim, Paula Upla, William J Rice, Jeremy Phillips, Benjamin L Timney, Ursula Pieper, Jeffrey B Bonanno, Javier Fernandez-Martinez, Zhanna Hakhverdyan, Natalia E Ketaren, Tsutomu Matsui, Thomas M Weiss, David L Stokes, J Michael Sauder, Stephen K Burley, Andrej Sali, Michael P Rout, and Steven C Almo. Structure, Dynamics, Evolution, and Function of a Major Scaffold Component in the Nuclear Pore Complex, 2013.

35. D Flemming, D P Devos, J Schwarz, S Amlacher, M Lutzmann, and E Hurt. Analysis of the yeast nucleoporin Nup188 reveals a conserved S-like structure with similarity to karyopherins. J Struct Biol, 177(1):99–105, 2012. doi: 10.1016/j.jsb.2011.11.008.

36. M Beck, S Mosalaganti, and J Kosinski. From the resolution revolution to evolution: structural insights into the evolutionary relationships between vesicle coats and the nuclear pore. Curr Opin Struct Biol, 52:32–40, 2018. doi: 10.1016/j.sbi.2018.07.012.

37. M C Field and M P Rout. Pore timing: the evolutionary origins of the nucleus and nuclear pore complex. F1000Res, 8:369, 2019. doi: 10.12688/f1000research.16402.1.

38. J Pulupa, H Prior, D S Johnson, and S M Simon. Conformation of the nuclear pore in living cells is modulated by transport state. Elife, 9, 2020. doi: 10.7554/eLife.60654.

39. Jessica Fischer, Roman Teimer, Stefan Amlacher, Ruth Kunze, and Ed Hurt. Linker Nups connect the nuclear pore complex inner ring with the outer ring and transport channel, 2015.

40. B Hampoelz, A Andres-Pons, P Kastritis, and M Beck. Structure and Assembly of the Nuclear Pore Complex. Annu Rev Biophys, 48:515–536, 2019. doi: 10.1146/annurev-biophys-052118-115308.

41. H Chug, S Trakhanov, B B Hulsmann, T Pleiner, and D Gorlich. Crystal structure of the metazoan Nup62*Nup58*Nup54 nucleoporin complex. Science, 350(6256):106–110, 2015. doi: 10.1126/science.aac7420.

42. Stefan Amlacher, Phillip Sarges, Dirk Flemming, Vera van Noort, Ruth Kunze, Damien P Devos, Manimozhiyan Arumugam, Peer Bork, and Ed Hurt. Insight into Structure and Assembly of the Nuclear Pore Complex by Utilizing the Genome of a Eukaryotic Thermophile, 2011.

43. D J Saltzberg, M Hepburn, K B Pilla, D C Schriemer, S P Lees-Miller, T L Blundell, and A Sali. SSEThread: Integrative threading of the DNA-PKcs sequence based on data from chemical cross-linking and hydrogen deuterium exchange. Prog Biophys Mol Biol, 147: 92–102, 2019. doi: 10.1016/j.pbiomolbio.2019.09.003.

44. N Dephoure, C Zhou, J Villen, S A Beausoleil, C E Bakalarski, S J Elledge, and S P Gygi. A quantitative atlas of mitotic phosphorylation. Proc Natl Acad Sci U S A, 105(31): 10762–10767, 2008. doi: 10.1073/pnas.0805139105.

45. E Laurell, K Beck, K Krupina, G Theerthagiri, B Bodenmiller, P Horvath, R Aebersold, W Antonin, and U Kutay. Phosphorylation of Nup98 by multiple kinases is crucial for NPC disassembly during mitotic entry. Cell, 144(4):539–550, 2011. doi: 10.1016/j.cell.2011.01.012.

46. M I Linder, M Kohler, P Boersema, M Weberruss, C Wandke, J Marino, C Ashiono, P Picotti, W Antonin, and U Kutay. Mitotic Disassembly of Nuclear Pore Complexes Involves CDK1-and PLK1-Mediated Phosphorylation of Key Interconnecting Nucleoporins. Dev Cell, 43(2):141–156 e7, 2017. doi: 10.1016/j.devcel.2017.08.020.

47. Damien Devos, Svetlana Dokudovskaya, Rosemary Williams, Frank Alber, Narayanan Eswar, Brian T Chait, Michael P Rout, and Andrej Sali. Simple fold composition and modular architecture of the nuclear pore complex, 2006.

48. M C Field, A Sali, and M P Rout. Evolution: On a bender–BARs, ESCRTs, COPs, and finally getting your coat. J Cell Biol, 193(6):963–972, 2011. doi: 10.1083/jcb.201102042.

49. A Bhardwaj and G Cingolani. Conformational selection in the recognition of the snurportin importin beta binding domain by importin beta. Biochemistry, 49(24):5042–5047, 2010. doi: 10.1021/bi100292y.

50. G Cingolani, C Petosa, K Weis, and C W Muller. Structure of importin-beta bound to the IBB domain of importin-alpha. Nature, 399(6733):221–229, 1999. doi: 10.1038/20367.

51. M Lutzmann, R Kunze, A Buerer, U Aebi, and E Hurt. Modular self-assembly of a Yshaped multiprotein complex from seven nucleoporins. EMBO J, 21(3):387–397, 2002. doi: 10.1093/emboj/21.3.387.

52. S Rajoo, P Vallotton, E Onischenko, and K Weis. Stoichiometry and compositional plasticity of the yeast nuclear pore complex revealed by quantitative fluorescence microscopy. Proc Natl Acad Sci U S A, 115(17):E3969–E3977, 2018. doi: 10.1073/pnas.1719398115.

53. C J Smoyer, S S Katta, J M Gardner, L Stoltz, S McCroskey, W D Bradford, M McClain, S E Smith, B D Slaughter, J R Unruh, and S L Jaspersen. Analysis of membrane proteins localizing to the inner nuclear envelope in living cells. J Cell Biol, 215(4):575–590, 2016. doi: 10.1083/jcb.201607043.

54. P Vallotton, S Rajoo, M Wojtynek, E Onischenko, A Kralt, C P Derrer, and K Weis. Mapping the native organization of the yeast nuclear pore complex using nuclear radial intensity measurements. Proc Natl Acad Sci U S A, 116(29):14606–14613, 2019. doi: 10.1073/pnas.1903764116.

55. V Galy, O Gadal, M Fromont-Racine, A Romano, A Jacquier, and U Nehrbass. Nuclear retention of unspliced mRNAs in yeast is mediated by perinuclear Mlp1. Cell, 116(1): 63–73, 2004. doi: 10.1016/s0092-8674(03)01026-2.

56. M Niepel, C Strambio-de Castillia, J Fasolo, B T Chait, and M P Rout. The nuclear pore complex-associated protein, Mlp2p, binds to the yeast spindle pole body and promotes its efficient assembly. J Cell Biol, 170(2):225–235, 2005. doi: 10.1083/jcb.200504140.

57. C Strambio-de Castillia, G Blobel, and M P Rout. Proteins connecting the nuclear pore complex with the nuclear interior, 1999.

58. C Feldherr, D Akin, and M S Moore. The nuclear import factor p10 regulates the functional size of the nuclear pore complex during oogenesis. J Cell Sci, 111 (Pt 13):1889–1896, 1998.

59. L G Trabuco, E Villa, K Mitra, J Frank, and K Schulten. Flexible fitting of atomic structures into electron microscopy maps using molecular dynamics. Structure, 16(5):673–683, 2008. doi: 10.1016/j.str.2008.03.005.

60. E M Romes, A Tripathy, and K C Slep. Structure of a yeast Dyn2-Nup159 complex and molecular basis for dynein light chain-nuclear pore interaction. J Biol Chem, 287(19): 15862–15873, 2012. doi: 10.1074/jbc.M111.336172.

61. M P Rout and G Blobel. Isolation of the yeast nuclear pore complex., 1993.

62. Caterina Strambio-de Castillia, Mario Niepel, and Michael P Rout. The nuclear pore complex: bridging nuclear transport and gene regulation, 2010.

63. Mario Niepel, Kelly R Molloy, Rosemary Williams, Julia C Farr, Anne C Meinema, Nicholas Vecchietti, Ileana M Cristea, Brian T Chait, Michael P Rout, and Caterina Strambio-de Castillia. The nuclear basket proteins Mlp1p and Mlp2p are part of a dynamic interactome including Esc1p and the proteasome., 2013.

64. C W Akey and M Radermacher. Architecture of the Xenopus nuclear pore complex revealed by three-dimensional cryo-electron microscopy, 1993.

65. Matthias Eibauer, Mauro Pellanda, Yagmur Turgay, Anna Dubrovsky, Annik Wild, and Ohad Medalia. Structure and gating of the nuclear pore complex, 2015.

66. Q Yang, M P Rout, and C W Akey. Three-dimensional architecture of the isolated yeast nuclear pore complex: functional and evolutionary implications, 1998.

67. Claire E Atkinson, Alexa L Mattheyses, Martin Kampmann, and Sanford M Simon. Conserved Spatial Organization of FG Domains in the Nuclear Pore Complex, 2013.

68. G Drin, J F Casella, R Gautier, T Boehmer, T U Schwartz, and B Antonny. A general amphipathic alpha-helical motif for sensing membrane curvature. Nat Struct Mol Biol, 14 (2):138–146, 2007. doi: 10.1038/nsmb1194.

69. S J Kim, J Fernandez-Martinez, P Sampathkumar, A Martel, T Matsui, H Tsuruta, T M Weiss, Y Shi, A Markina-Inarrairaegui, J B Bonanno, J M Sauder, S K Burley, B T Chait, S C Almo, M P Rout, and A Sali. Integrative structure-function mapping of the nucleoporin Nup133 suggests a conserved mechanism for membrane anchoring of the nuclear pore complex. Mol Cell Proteomics, 13(11):2911–2926, 2014. doi: 10.1074/mcp.M114.040915.

70. N Meszaros, J Cibulka, M J Mendiburo, A Romanauska, M Schneider, and A Kohler. Nuclear pore basket proteins are tethered to the nuclear envelope and can regulate membrane curvature. Dev Cell, 33(3):285–298, 2015. doi: 10.1016/j.devcel.2015.02.017.

71. Y Shi, J Fernandez-Martinez, E Tjioe, R Pellarin, S J Kim, R Williams, D Schneidman-Duhovny, A Sali, M P Rout, and B T Chait. Structural characterization by cross-linking reveals the detailed architecture of a coatomer-related heptameric module from the nuclear pore complex. Mol Cell Proteomics, 13(11):2927–2943, 2014. doi: 10.1074/mcp.M114.041673.

72. J Mansfeld, S Guttinger, L A Hawryluk-Gara, N Pante, M Mall, V Galy, U Haselmann, P Muhlhausser, R W Wozniak, I W Mattaj, U Kutay, and W Antonin. The conserved trans-membrane nucleoporin NDC1 is required for nuclear pore complex assembly in vertebrate cells. Mol Cell, 22(1):93–103, 2006. doi: 10.1016/j.molcel.2006.02.015.

73. Benjamin Vollmer, Allana Schooley, Ruchika Sachdev, Nathalie Eisenhardt, Anna M Schneider, Cornelia Sieverding, Johannes Madlung, Uwe Gerken, Boris Macek, and Wolfram Antonin. Dimerization and direct membrane interaction of Nup53 contribute to nuclear pore complex assembly, 2012.

74. M Marelli, C P Lusk, H Chan, J D Aitchison, and R W Wozniak. A link between the synthesis of nucleoporins and the biogenesis of the nuclear envelope, 2001.

75. Q Hao, B Zhang, K Yuan, H Shi, and G Blobel. Electron microscopy of Chaetomium pom152 shows the assembly of ten-bead string. Cell Discov, 4:56, 2018. doi: 10.1038/s41421-018-0057-7.

76. P Upla, S J Kim, P Sampathkumar, K Dutta, S M Cahill, I E Chemmama, R Williams, J B Bonanno, W J Rice, D L Stokes, D Cowburn, S C Almo, A Sali, M P Rout, and J Fernandez-Martinez. Molecular Architecture of the Major Membrane Ring Component of the Nuclear Pore Complex. Structure, 25(3):434–445, 2017. doi: 10.1016/j.str.2017.01.006.

77. Colin P C De Souza, Aysha H Osmani, Shahr B Hashmi, and Stephen A Osmani. Partial nuclear pore complex disassembly during closed mitosis in Aspergillus nidulans, 2004.

78. M Heusel, M Frank, M Kohler, S Amon, F Frommelt, G Rosenberger, I Bludau, S Aulakh, M I Linder, Y Liu, B C Collins, M Gstaiger, U Kutay, and R Aebersold. A Global Screen for Assembly State Changes of the Mitotic Proteome by SEC-SWATH-MS. Cell Syst, 10(2): 133–155 e6, 2020. doi: 10.1016/j.cels.2020.01.001.

79. U Kutay, R Jühlen, and W Antonin. Mitotic disassembly and reassembly of nuclear pore complexes. Trends Cell Biol, 21:132–139, 2021. doi: 10.1016/j.tcb.2021.06.011.

80. Kasper R Andersen, Evgeny Onischenko, Jeffrey H Tang, Pravin Kumar, James Z Chen, Alexander Ulrich, Jan T Liphardt, Karsten Weis, and Thomas U Schwartz. Scaffold nucleoporins Nup188 and Nup192 share structural and functional properties with nuclear transport receptors, 2013.

81. T Stuwe, D H Lin, L N Collins, E Hurt, and A Hoelz. Evidence for an evolutionary relationship between the large adaptor nucleoporin Nup192 and karyopherins. Proc Natl Acad Sci U S A, 111(7):2530–2535, 2014. doi: 10.1073/pnas.1311081111.

82. M C Sumner and J Brickner. The Nuclear Pore Complex as a Transcription Regulator. Cold Spring Harb Perspect Biol, page a039438, 2021. doi: 10.1101/cshperspect.a039438.

83. U H Cho and M W Hetzer. Nuclear Periphery Takes Center Stage: The Role of Nuclear Pore Complexes in Cell Identity and Aging. Neuron, 106(6):899–911, 2020. doi: 10.1016/j.neuron.2020.05.031.

84. V Nofrini, D Di Giacomo, and C Mecucci. Nucleoporin genes in human diseases. Eur J Hum Genet, 24(10):1388–1395, 2016. doi: 10.1038/ejhg.2016.25.

85. Alberto Elosegui-Artola, Ion Andreu, Amy E M Beedle, Ainhoa Lezamiz, Marina Uroz, Anita J Kosmalska, Roger Oria, Jenny Z Kechagia, Palma Rico-Lastres, Anabel-Lise Le Roux, Catherine M Shanahan, Xavier Trepat, Daniel Navajas, Sergi Garcia-Manyes, and Pere Roca-Cusachs. Force Triggers YAP Nuclear Entry by Regulating Transport across Nuclear Pores. Cell, 171(6):1397–1410.e14, nov 2017. doi: 10.1016/j.cell.2017.10.008.

86. Emanuela Jacchetti, Ramin Nasehi, Lucia Boeri, Valentina Parodi, Alessandro Negro, Diego Albani, Roberto Osellame, Giulio Cerullo, Jose Felix Rodriguez Matas, and Manuela Teresa Raimondi. The nuclear import of the transcription factor MyoD is reduced in mesenchymal stem cells grown in a 3D micro-engineered niche. Scientific reports, 11 (1):3021, feb 2021. doi: 10.1038/s41598-021-81920-2.

87. Elena Kassianidou, Joanna Kalita, and Roderick Y.H. Lim. The role of nucleocytoplasmic transport in mechanotransduction. Experimental Cell Research, 377(1):86–93, 2019. doi: https://doi.org/10.1016/j.yexcr.2019.02.009.

88. Y Xiang, S Nambulli, Z Xiao, H Liu, Z Sang, W P Duprex, D Schneidman-Duhovny, C Zhang, and Y Shi. Versatile and multivalent nanobodies efficiently neutralize SARS-CoV-2. Science, 370(6523):1479–1484, 2020. doi: 10.1126/science.abe4747.

89. Y Xiang, Z Sang, L Bitton, J Xu, Y Liu, D Schneidman-Duhovny, and Y Shi. Integrative proteomics identifies thousands of distinct, multi-epitope, and high-affinity nanobodies. Cell Syst, 12(3):220–234 e9, 2021. doi: 10.1016/j.cels.2021.01.003.

90. Z L Chen, J M Meng, Y Cao, J L Yin, R Q Fang, S B Fan, C Liu, W F Zeng, Y H Ding, D Tan, L Wu, W J Zhou, H Chi, R X Sun, M Q Dong, and S M He. A high-speed search engine pLink 2 with systematic evaluation for proteome-scale identification of cross-linked peptides. Nat Commun, 10(1):3404, 2019. doi: 10.1038/s41467-019-11337-z.

91. James R Kremer, David N Mastronarde, and J. Richard McIntosh. Computer Visualization of Three-Dimensional Image Data Using IMOD. Journal of Structural Biology, 116(1): 71–76, 1996. doi: 10.1006/jsbi.1996.0013.

92. S Q Zheng. MotionCor2: anisotropic correction of beam-induced motion for improved cryo-electron microscopy. Nature methods, 14:331–332.

93. K Zhang. Gctf: Real-time CTF determination and correction. J Struct Biol, 193(1):1–12, 2016. doi: 10.1016/j.jsb.2015.11.003.

94. G Tang, L Peng, P R Baldwin, D S Mann, W Jiang, I Rees, and S J Ludtke. EMAN2: an extensible image processing suite for electron microscopy. J Struct Biol, 157(1):38–46, 2007. doi: 10.1016/j.jsb.2006.05.009.

95. T Nakane, D Kimanius, E Lindahl, and S H Scheres. Characterisation of molecular motions in cryo-EM single-particle data by multi-body refinement in RELION. Elife, 7, 2018. doi: 10.7554/eLife.36861.

96. Y T Liu, J Jih, X Dai, G Q Bi, and Z H Zhou. Cryo-EM structures of herpes simplex virus type 1 portal vertex and packaged genome. Nature, 570(7760):257–261, 2019. doi: 10.1038/s41586-019-1248-6.

97. Eric F Pettersen, Thomas D Goddard, Conrad C Huang, Gregory S Couch, Daniel M Greenblatt, Elaine C Meng, and Thomas E Ferrin. UCSF Chimera - A visualization system for exploratory research and analysis, 2004.

98. A Casanal, B Lohkamp, and P Emsley. Current developments in Coot for macromolecular model building of Electron Cryo-microscopy and Crystallographic Data. Protein Sci, 29(4): 1069–1078, 2020. doi: 10.1002/pro.3791.

99. P V Afonine, B K Poon, R J Read, O V Sobolev, T C Terwilliger, A Urzhumtsev, and P D Adams. Real-space refinement in PHENIX for cryo-EM and crystallography. Acta Crystallogr D Struct Biol, 74(Pt 6):531–544, 2018. doi: 10.1107/S2059798318006551.

100. Frank Alber, Svetlana Dokudovskaya, Liesbeth M Veenhoff, Wenzhu Zhang, Julia Kipper, Damien Devos, Adisetyantari Suprapto, Orit Karni-Schmidt, Rosemary Williams, Brian T Chait, Andrej Sali, and Michael P Rout. The molecular architecture of the nuclear pore complex, 2007.

101. M P Rout and A Sali. Principles for Integrative Structural Biology Studies. Cell, 177(6): 1384–1403, 2019. doi: 10.1016/j.cell.2019.05.016.

102. D Russel, K Lasker, B Webb, J Velazquez-Muriel, E Tjioe, D Schneidman-Duhovny, B Peterson, and A Sali. Putting the pieces together: integrative modeling platform software for structure determination of macromolecular assemblies. PLoS Biol, 10(1):e1001244, 2012. doi: 10.1371/journal.pbio.1001244.

103. A Sali. From integrative structural biology to cell biology. J Biol Chem, page 100743, 2021. doi: 10.1016/j.jbc.2021.100743.

104. S Viswanath, I E Chemmama, P Cimermancic, and A Sali. Assessing Exhaustiveness of Stochastic Sampling for Integrative Modeling of Macromolecular Structures. Biophys J, 113(11):2344–2353, 2017. doi: 10.1016/j.bpj.2017.10.005.

105. Peter V. Hornbeck, Bin Zhang, Beth Murray, Jon M. Kornhauser, Vaughan Latham, and Elzbieta Skrzypek. Phosphositeplus, 2014: mutations, ptms and recalibrations. Nucleic Acids Research, 43:D512–D520, 2015.

106. Felix Roman Wagner, Ruud Schampers, Reika Watanabe, Digvijay Singh, Hans Persoon, Miroslava Schaffer, Peter Fruhstorfer, and Elizabeth Villa. Preparing samples from whole cells using focused-ion-beam milling for cryo-electron tomography. Nature Protocols, 2020.

107. David N. Mastronarde. Automated electron microscope tomography using robust prediction of specimen movements. Journal of Structural Biology, 152(1):36–51, oct 2005. doi: 10.1016/j.jsb.2005.07.007.

108. Z A Chen and J Rappsilber. Quantitative cross-linking/mass spectrometry to elucidate structural changes in proteins and their complexes. Nat Protoc, 14(1):171–201, 2019. doi: 10.1038/s41596-018-0089-3.

109. P A Penczek. Reliable cryo-EM resolution estimation with modified Fourier shell correlation. IUCrJ, 7(Pt 6):995–1008, 2020. doi: 10.1107/S2052252520011574.

110. G D Pintilie, J Zhang, T D Goddard, W Chiu, and D C Gossard. Quantitative analysis of cryo-EM density map segmentation by watershed and scale-space filtering, and fitting of structures by alignment to regions. J Struct Biol, 170(3):427–438, 2010. doi: 10.1016/j.jsb.2010.03.007.

111. Muyuan Chen, James M. Bell, Xiaodong Shi, Stella Y. Sun, Zhao Wang, and Steven J. Ludtke. A complete data processing workflow for cryo-ET and subtomogram averaging. Nature Methods, 16(11):1161–1168, nov 2019. doi: 10.1038/s41592-019-0591-8.

112. M Bonomi, S Hanot, C H Greenberg, A Sali, M Nilges, M Vendruscolo, and R Pellarin. Bayesian Weighing of Electron Cryo-Microscopy Data for Integrative Structural Modeling. Structure, 27(1):175–188 e6, 2019. doi: 10.1016/j.str.2018.09.011.

113. R H Swendsen and J S Wang. Replica Monte Carlo simulation of spin glasses. Phys Rev Lett, 57(21):2607–2609, 1986. doi: 10.1103/PhysRevLett.57.2607.

114. D J Saltzberg, S Viswanath, I Echeverria, I E Chemmama, B Webb, and A Sali. Using Integrative Modeling Platform to compute, validate, and archive a model of a protein complex structure. Protein Sci, 30(1):250–261, 2021. doi: 10.1002/pro.3995.

