## Supplementary Data for "Comprehensive Structure and Functional Adaptations of the Yeast Nuclear Pore Complex"

***These authors contributed equally**

### Supplementary Info

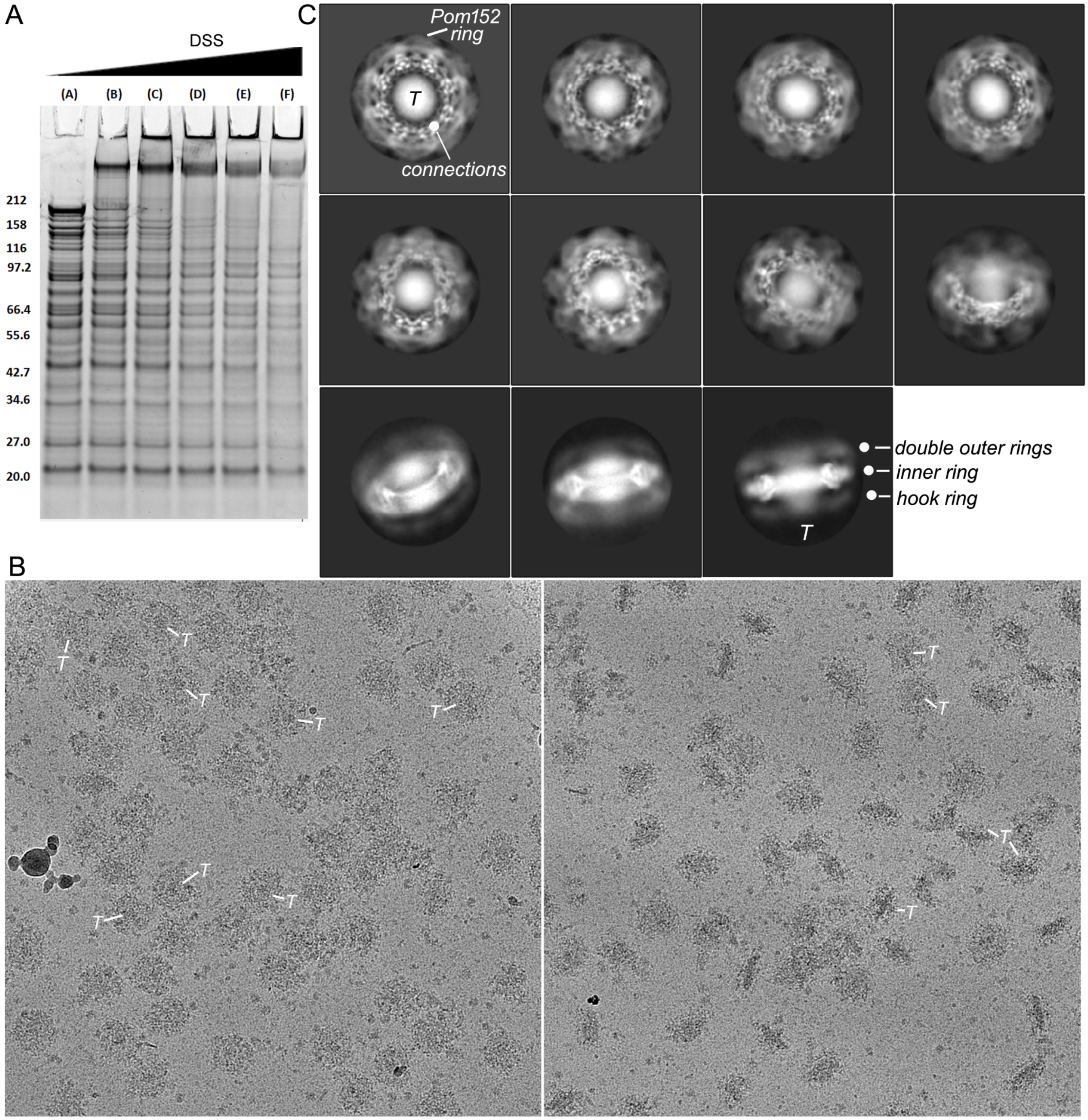


**Fig. S1. Sample preparation and characterization of NPCs. A**. Affinity purified NPCs were titrated with increasing concentrations of a bifunctional crosslinker (DSS) in an attempt to stabilize flexible regions. **B**. Typical micrographs of the DSS crosslinked yeast NPC imaged on a carbon film supported over holes. Images were taken at 160 kV on a Tecnai F20 microscope. (left) NPCs in thinner ice with top and slightly tilted views. (right) NPCs with a larger distribution of tilted and edge-on views. A central, ring-like transporter (T) is visible in most particles. **C**. 2D class averages show a range of views. Low density features may represent more flexible regions. Density for the double outer ring, central transporter and connections to the inner ring are evident along with the Pom152 ring.

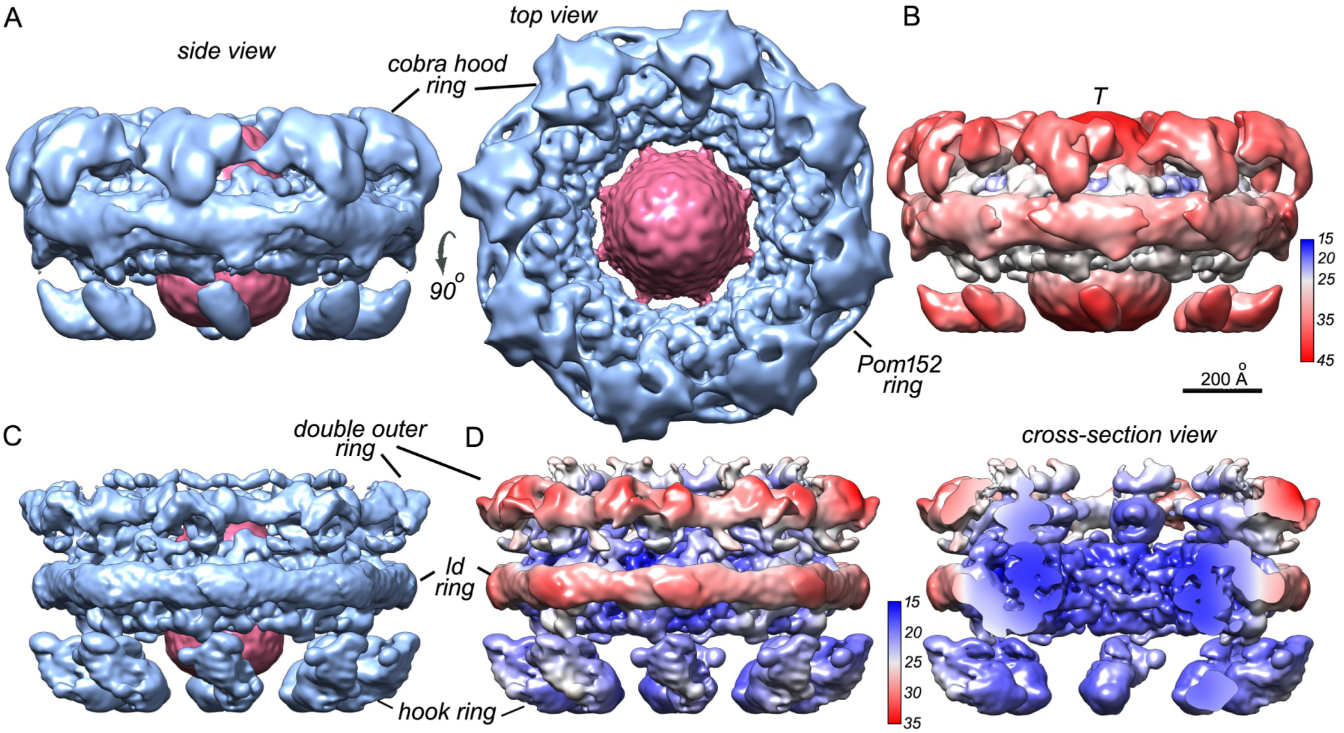


**Fig. S2. Initial single particle 3D maps of the isolated NPC refined with C8 symmetry. A**. (left) The initial single particle 3D map of the isolated NPC is similar to a previous structure determined with sub-tomogram averaging (**?** ) (scaffold in blue, transporter in red). (right) The cobra hood feature and Pom152 ring are visible in the top view. **B**. Local resolution map for 3D map in panel A. Scale bar 200 Å. **C**. Side view of a recombined yeast NPC 3D map: each ring was refined separately with C8 symmetry and a focused soft mask to improve local resolution. **D**. (left and right) Side and cross-section views show improved local resolution for the combined 3D map in panel C.

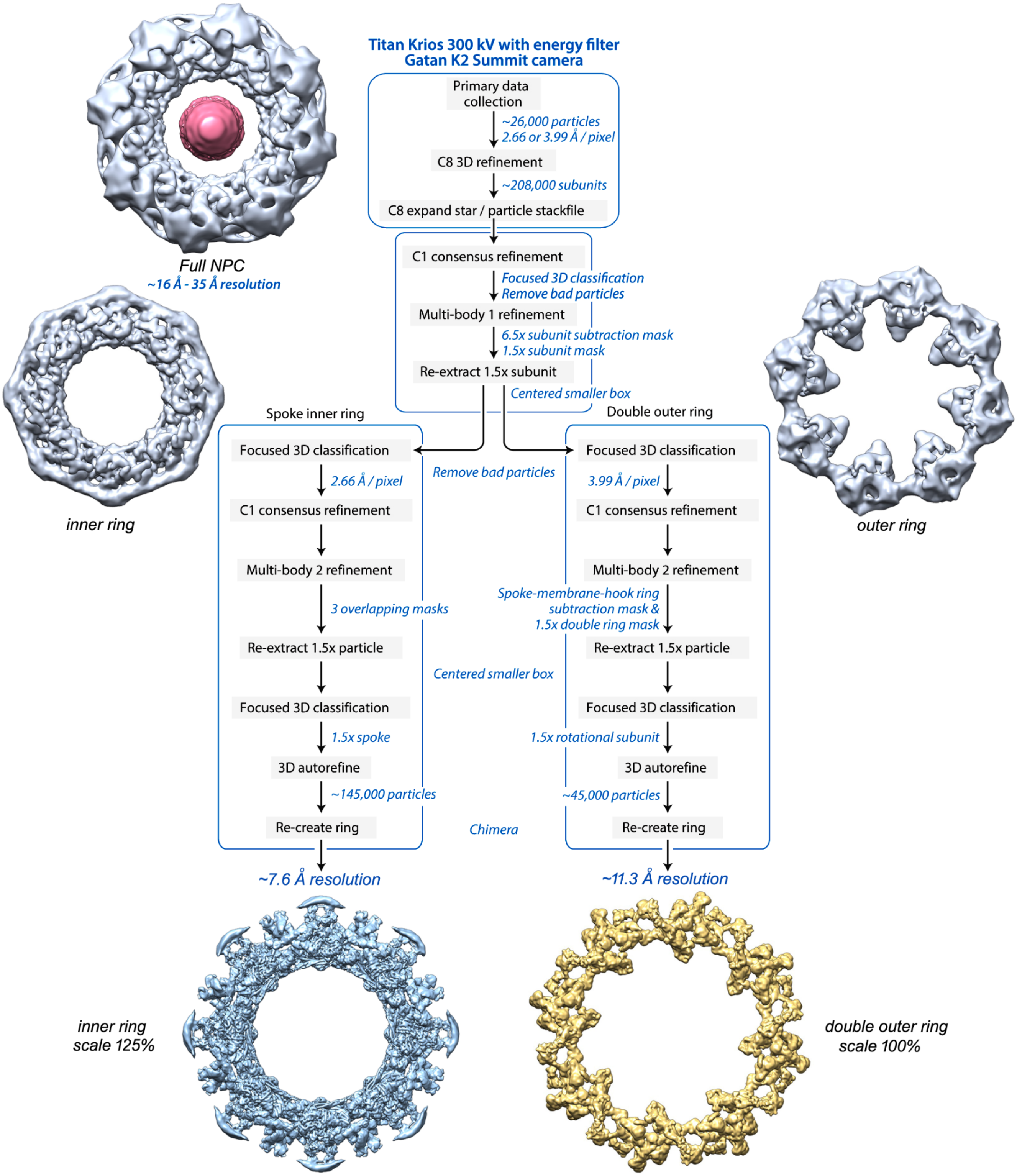


**Fig. S3. A flowchart diagram for multi-body image processing of the spoke and double outer ring.** Resolution for the spoke and double outer ring improved ∼3-fold after multi-body and downstream processing. (top) Images are shown for initial structures of the NPC, along with the inner and outer rings aligned with C8 symmetry and a focused mask. (bottom) Recombined 3D maps after two iterations of multi-body processing. Note scale difference.

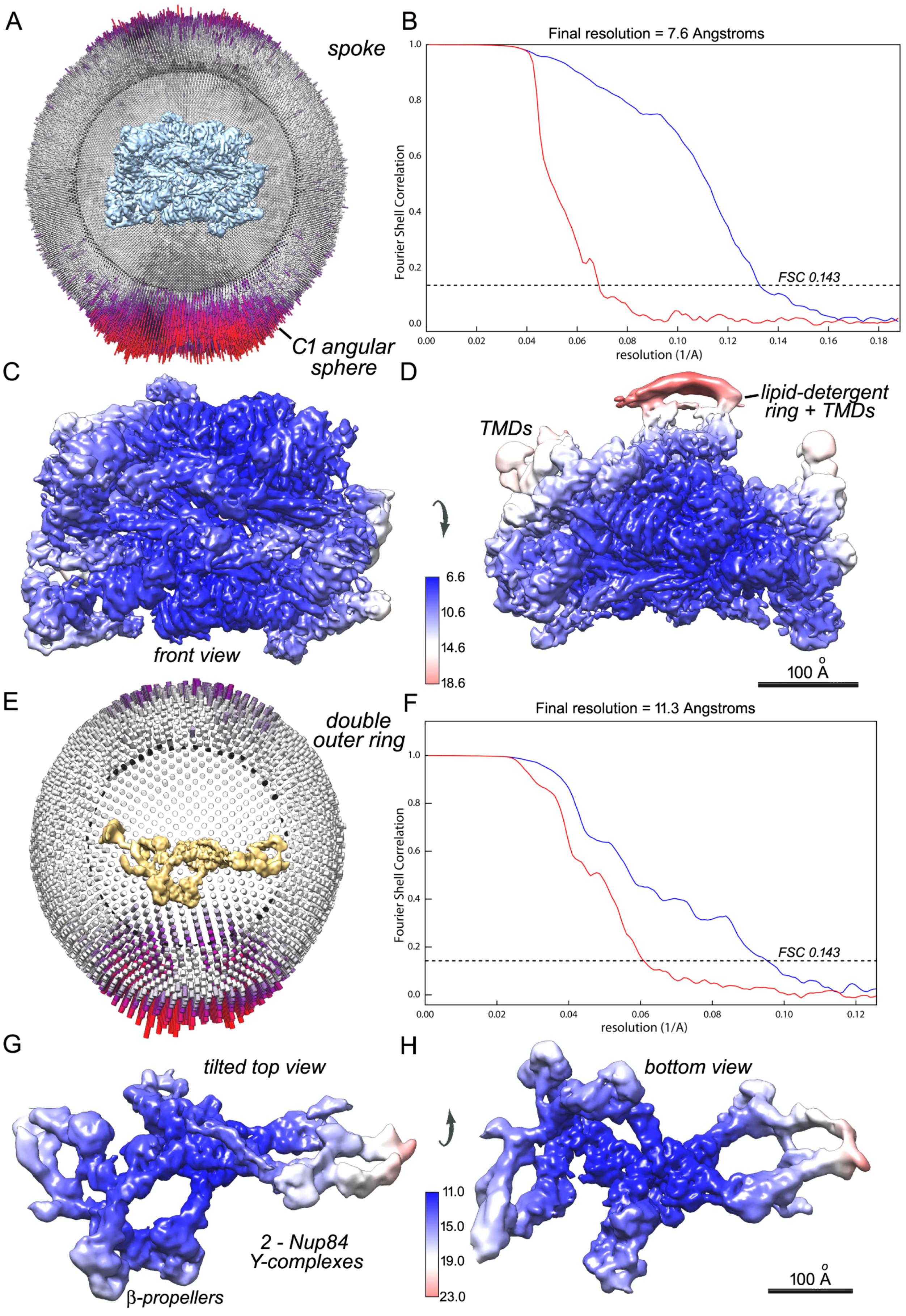
A
spoke
C1 angular sphere
B
Final resolution = 7.6 Angstroms
Fourier Shell Correlation
FSC 0.143
resolution (1/A)
C
front view
D
lipid-detergent ring + TMDs
TMDs
6.6
10.6
14.6
18.6
100 Å
E
double outer ring
F
Final resolution = 11.3 Angstroms
Fourier Shell Correlation
FSC 0.143
resolution (1/A)
G
tilted top view
2 - Nup84 Y-complexes
β-propellers
H
bottom view
11.0
15.0
19.0
23.0
100 Å

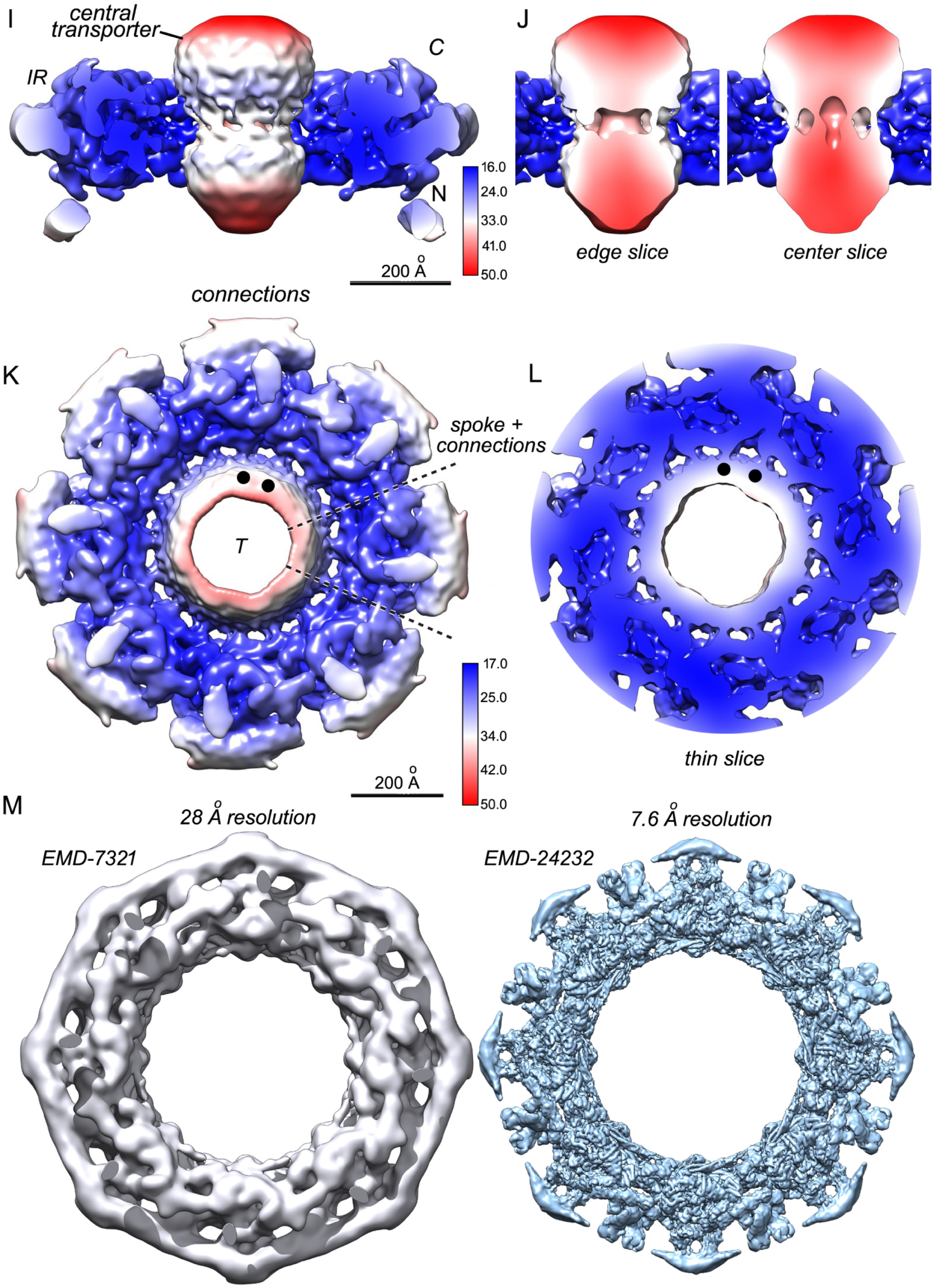


**Fig. S4. Angular coverage and resolution for single particle 3D maps. A**. Angular distribution for C1 refinement of a 1.5x spoke volume with inset map in blue. The 8-fold axis is aligned roughly vertical and the tendency to sit in a range of preferred orientations is apparent, which may involve basket interactions with the carbon support film. However, no obvious loss of angular resolution occurs due to the overall distribution of views. **B**. Fourier shell curve for a spoke with a final FSC0.143 resolution of 7.6 Å. Masked curve in blue, phase randomized curve in red. **C-D**. Local resolution calculated in RELION and plotted onto the spoke isosurface. Left- face view of the spoke, Right- 90º tilted view from the top showing part of the lipid-detergent ring and likely TMDs. **E**. Angular distribution for C1 refinement of the double outer ring with inset map in gold. This map contains roughly 1.5x rotational subunits, each with a pair of offset Nup84 Y-complexes **F**. Fourier shell curve for the double outer ring with a final FSC0.143 resolution of ∼11.3 Å. A generous mask was used in these calculations. **G**. Local resolution calculated in RELION and plotted on the double outer ring as viewed from the outside edge of the ring. Central dimples are present in each β-propeller. **H**. A reverse view showing the underside of the double ring. Lower resolution features at the edges were not included in the reconstituted full ring. <page 2> **I**. Local resolution for a global 3D map of the central transporter and inner ring. **J**. Two cross-sections through the central transporter. Note the radial and vertical resolution gradient within this feature. **K**. Focused 3D refinement gives a local resolution of ∼20-25 Å for connections that cross between the inner ring and central transporter (features above the black dots) . **L**. A section through the connections is shown for the local resolution 3D map.

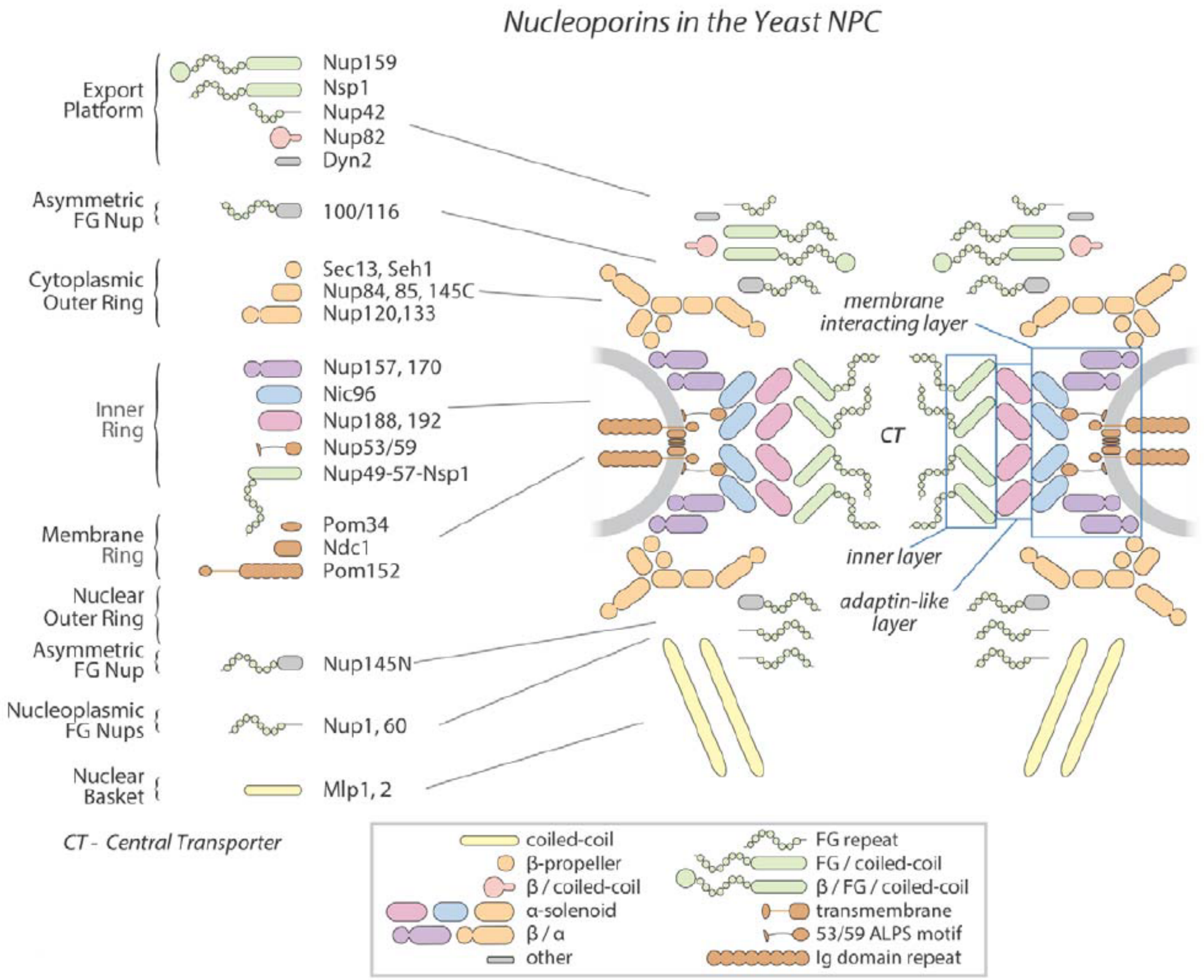


**Fig. S5.** Nucleoporins within the yeast NPC are shown diagrammatically. The core scaffold with additional Nups is broken down into distinct vertical levels or rings, while radial layers within a spoke are indicated within light blue boxes on the right.

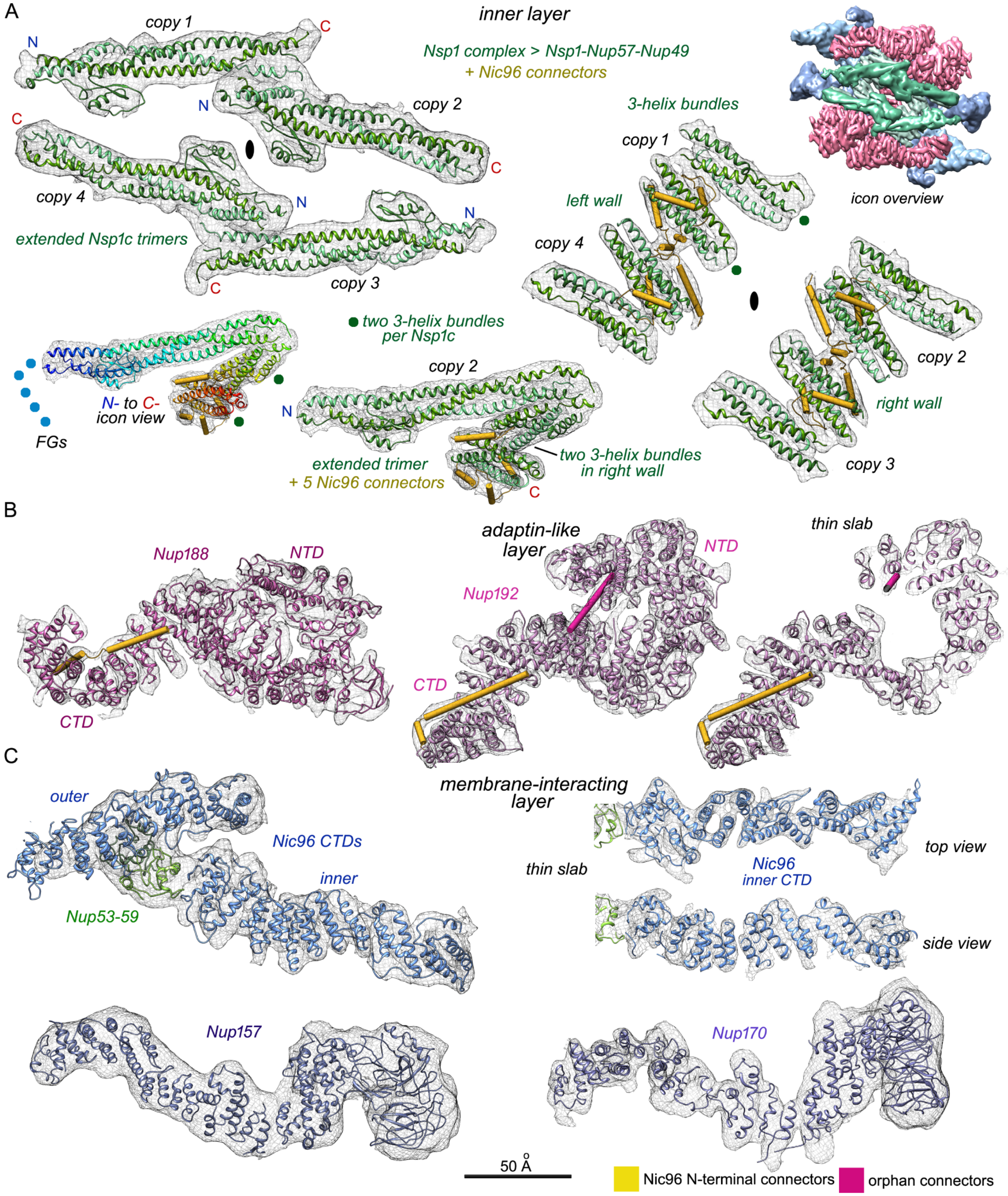


**Fig. S6. Density segmentation for Nups in the isolated spoke.** Electron density maps are shown as a black meshwork superimposed on a thin ribbon representation for each Nup. **A**. Four extended Nsp1 complex trimers from the inner layer of the spoke. Panels: (left) Nsp1 heterotrimers are comprised of extended trimers that face the central channel; individual copies are indicated (copy 1-4, N- to C-) along with the local 2-fold axis (black ellipse). (right) Left and right diagonal walls formed by 3-helix bundles with bound Nic96 connectors. (upper corner) An icon view shows the color coding for three layers in the spoke that is followed in this figure. (lower left and middle) An icon view in standard rainbow coloring on the left shows the N- to Cconnectivity for helices in copy 2. FG domains at the N-terminus are indicated by blue dots. A complete Nsp1 complex is shown for copy 2 with 3-helix bundles and Nic96 connectors that contribute to the right-hand wall. **B**. Nups from the adaptin-like layer. (left) Nup188 with Nic96 connector helices. (right) Nup192 with helical connectors (in gold and pink). **C**. Nups from the membrane interacting layer are shown; (top) the Nic96 CTD with a Nup53- Nup59 pair at two different thresholds; (bottom) the paralogs Nup157 (left) and Nup170 (right) in their respective density maps.

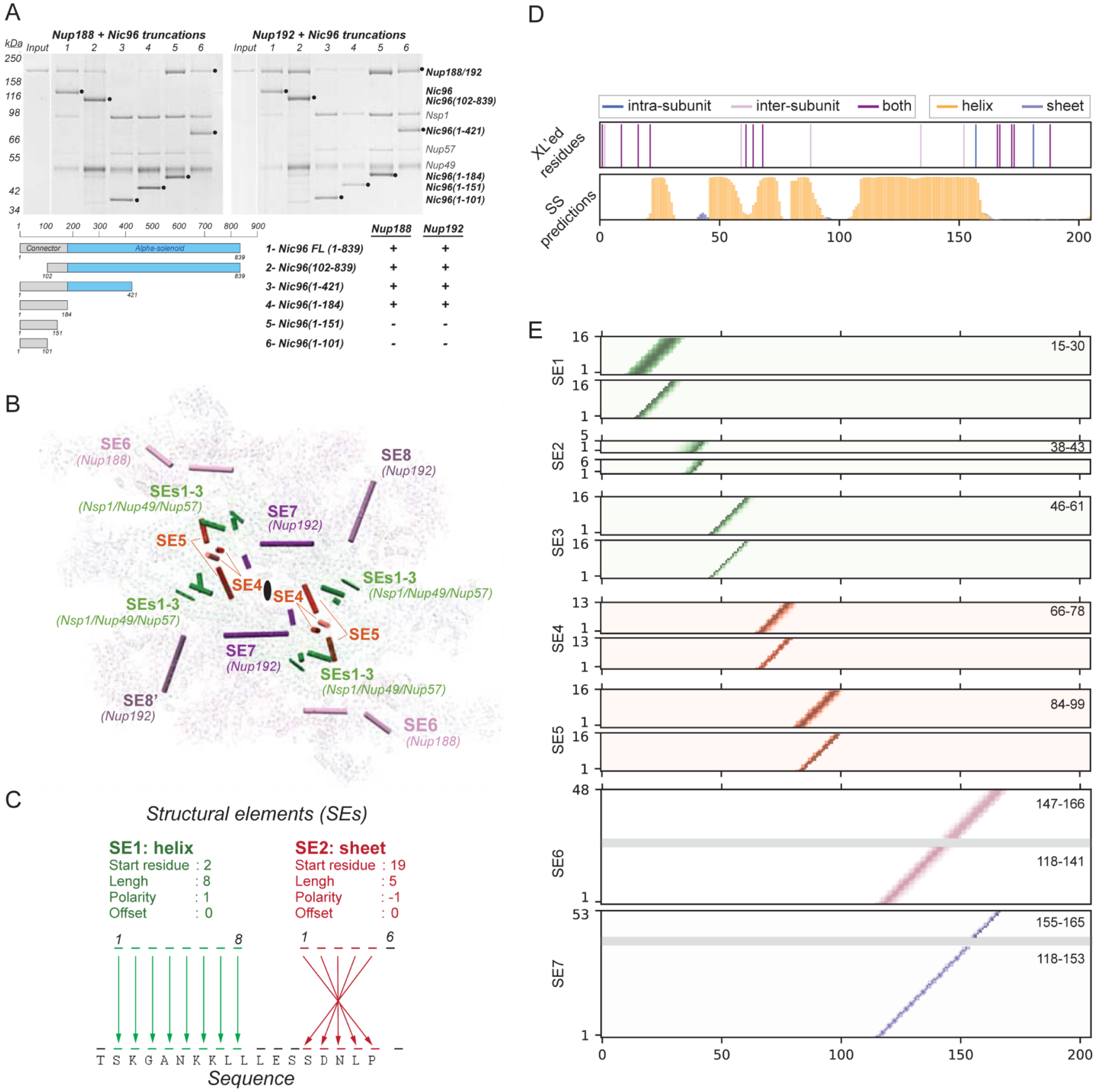
A
Nup188 + Nic96 truncations
Nup192 + Nic96 truncations
kDa
Input
250
158
116
98
66
55
42
34
Nup188/192
Nic96
Nic96(102-839)
Nsp1
Nic96(1-421)
Nup57
Nup49
Nic96(1-184)
Nic96(1-151)
Nic96(1-101)
Connector
Alpha-solenoid
Nup188
Nup192
1- Nic96 FL (1-839)
2- Nic96(102-839)
3- Nic96(1-421)
4- Nic96(1-184)
5- Nic96(1-151)
6- Nic96(1-101)
B
SE6
(Nup188)
SEs1-3
(Nsp1/Nup49/Nup57)
SE7
(Nup192)
SE8
(Nup192)
SE5
SE4
SE8'
C
Structural elements (SEs)
SE1: helix
Start residue : 2
Lengh : 8
Polarity : 1
Offset : 0
SE2: sheet
Start residue : 19
Lengh : 5
Polarity : -1
Offset : 0
T S K G A N K K L L L E S S D N L P
Sequence
D
intra-subunit
inter-subunit
both
helix
sheet
XL'ed residues
SS predictions
E
SE1
SE2
SE3
SE4
SE5
SE6
SE7
15-30
38-43
46-61
66-78
84-99
147-166
118-141
155-165
118-153

F

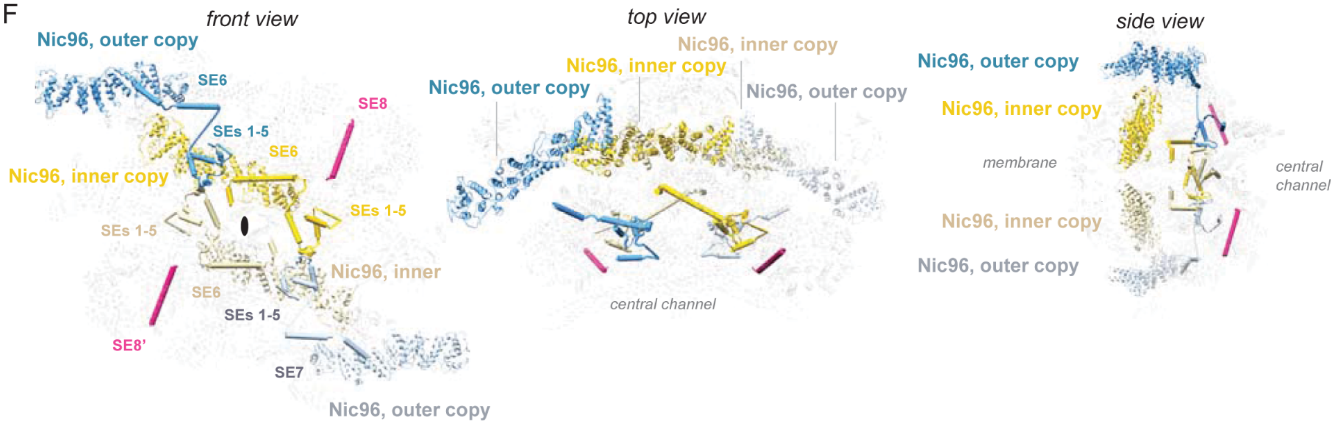


G

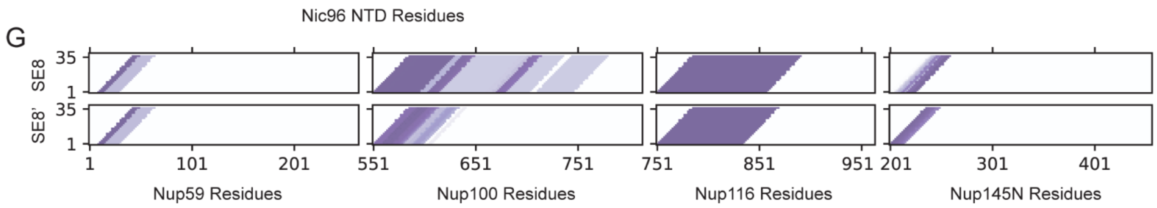


**Fig. S7. Integrative threading of orphan structure elements.** **A**. Nup188 and Nup192 interact within an overlapping region of the Nic96 flexible connector. (Top) SDS-PAGE gels with results of in vitro reconstitution experiments between Nup188 (left) or Nup192 (right) with different Nic96 truncation mutants. Relevant proteins are indicated with a dot and their identity is indicated on the right. (Bottom) Schematic showing the boundaries of the Nic96 constructs (left) and a summary of the in vitro reconstitution results (right). FL, full length. **B**. The SEs in a spoke are indicated and some elements (SEs 6 and 7) include features from two orphan densities. Based on symmetry, the SEs are grouped into five groups: 1) SEs 1-3 bind to the Nsp1/Nup49/Nup57 coiled-coils (green), 2) SEs 4 and 5 localize between copies of the Nsp1/Nup49/Nup57 coiled-coils (red) at the interface between two half walls, 3) SE 6 binds to the Nup188 CTD (pink), 4) SE 7 binds to the Nup192 CTD (dark purple), and 5) SE 8 and 8' bind to the Nup192 NTDs (light purple). **C**. Application of SE movers for integrative threading. The specification of an SE includes a secondary structure designation, a set of Cα coordinates, and four keys that map these coordinates to residues in the primary sequence. For example, SE1 defines the Cα coordinates of ten residues of a helix and the set of four keys map the six green coordinates onto sequence. The start residue, 2, denotes that the threaded sequence begins at residue two, the length, 10, denotes that the SE is 10 residues long, and a polarity of 1 indicates an increasing residue index. SE2 shows a similar assignment, beginning at residue 19; however, the polarity of -1 flips the assignment, such that the last assigned coordinate in the SE is threaded to the sequence at residue 19 and the remainder of the SE is assigned in reverse order. **D**. Relevant cross-links and secondary structure predictions used for integrative threading of the Nic96 NTD. **E**. Residue occupancy of good-scoring models following enumeration of all possible states in the integrative threading. Each bin represents the mapping of a residue in sequence (X-axis) to a coordinate in a structure element (Y-axis). Darker colors indicate that 100models map the corresponding residue to the structure element coordinate. Grey boxes indicate regions in which the helices are bent and/or have missing segments. <page 2> **F**. Centroid assignments of the orphan SEs 1-7 to Nic96 NTDs; front view (left), top view (middle), and side view (right). Of the SEs in the spoke, only SE8 and 8' are not assigned to Nic96 (magenta). **G**. Assignments for SE 8 and 8' are ambiguous, including possible segments in Nup59, 100, 116, and 145N.

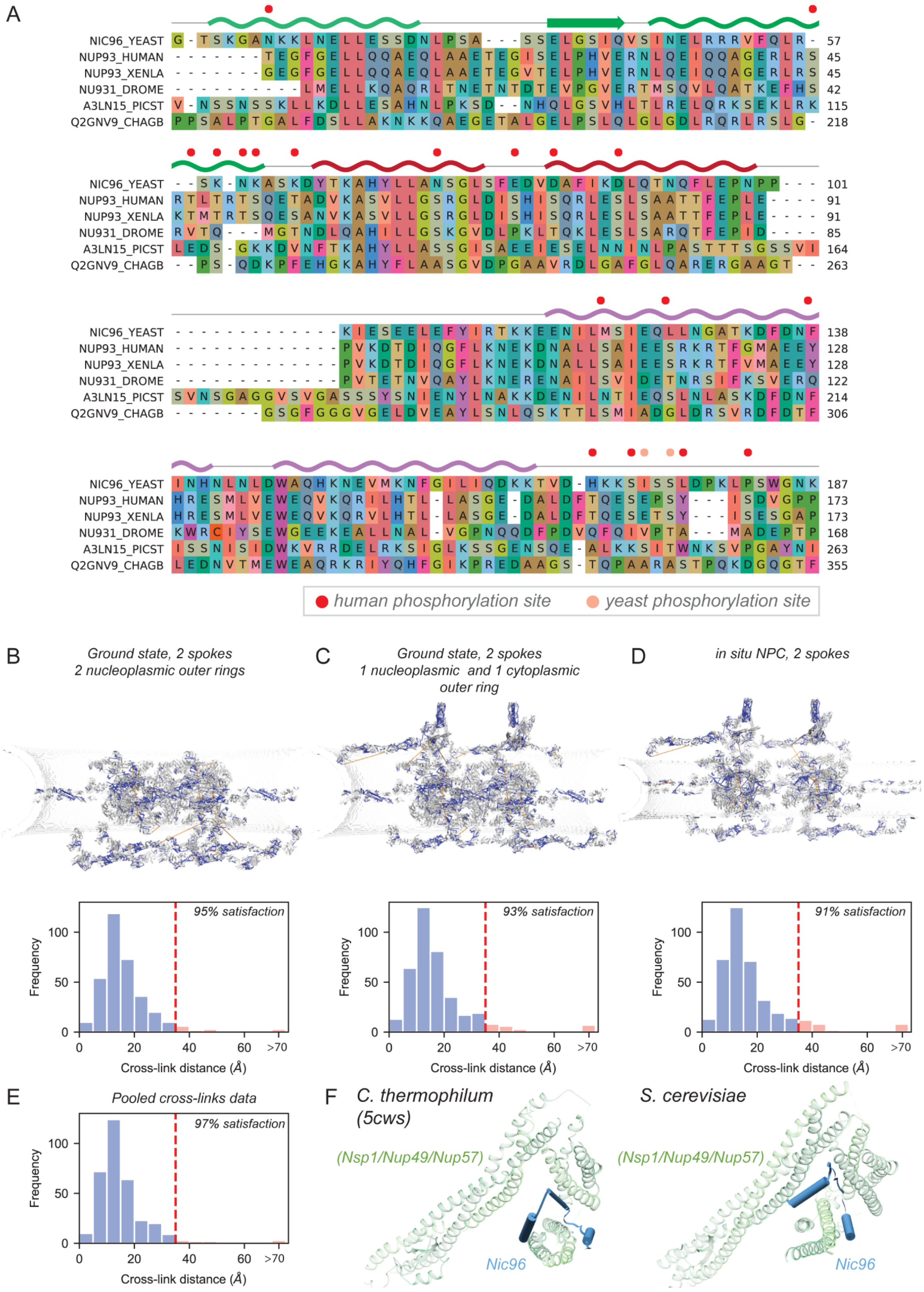


**Fig. S8. Validation of NPC structures** A. Sequence alignment of Nic96 NTD orthologs from S. cerevisiae (yeast), Homo sapiens (human), Xenopus laevis (xenla), Drosophila melanogaster (drome), Pichia stipitis (picst), and Chaetomium globosum (chagb). The secondary structure of the SE centroid assignments is shown above the sequence alignment. Mapped phosphorylation sites from human and yeast are shown as red and pink dots, respectively. B. Satisfaction of unique chemical cross-links for the cryo-EM structure of the isolated NPC. Identified chemical cross-links were mapped onto the structure of two adjacent spokes of the NPC. Satisfied cross-links with Cα–Cα distances that fall below the distance threshold of 35 Å are shown in blue. Violated cross-links with Cα–Cα distances that are larger than 35 Å are shown in orange. The histogram below the model shows the distribution of the cross-linked Cα–Cα distances, validating the NPC structures. C. Same as panel B for a model of the isolated NPC created with one nucleoplasmic and one cytoplasmic outer ring that frame a radially-constricted inner ring. D. Same as panel B for the current in situ NPC structure. E. Cross-link satisfaction summed over all three models. A cross-link restraint is satisfied if the corresponding Cα–Cα distance is less than 35 Å in any of the structures in panels C-E. F. (left) Ribbon representation of the X-ray structure of the Nsp1/Nup49/Nup57-Nic96 N-terminal SLiM complex from C. thermophilum. (right) Ribbon representation of the current cryo-EM structure for this complex from S. cerevisiae.

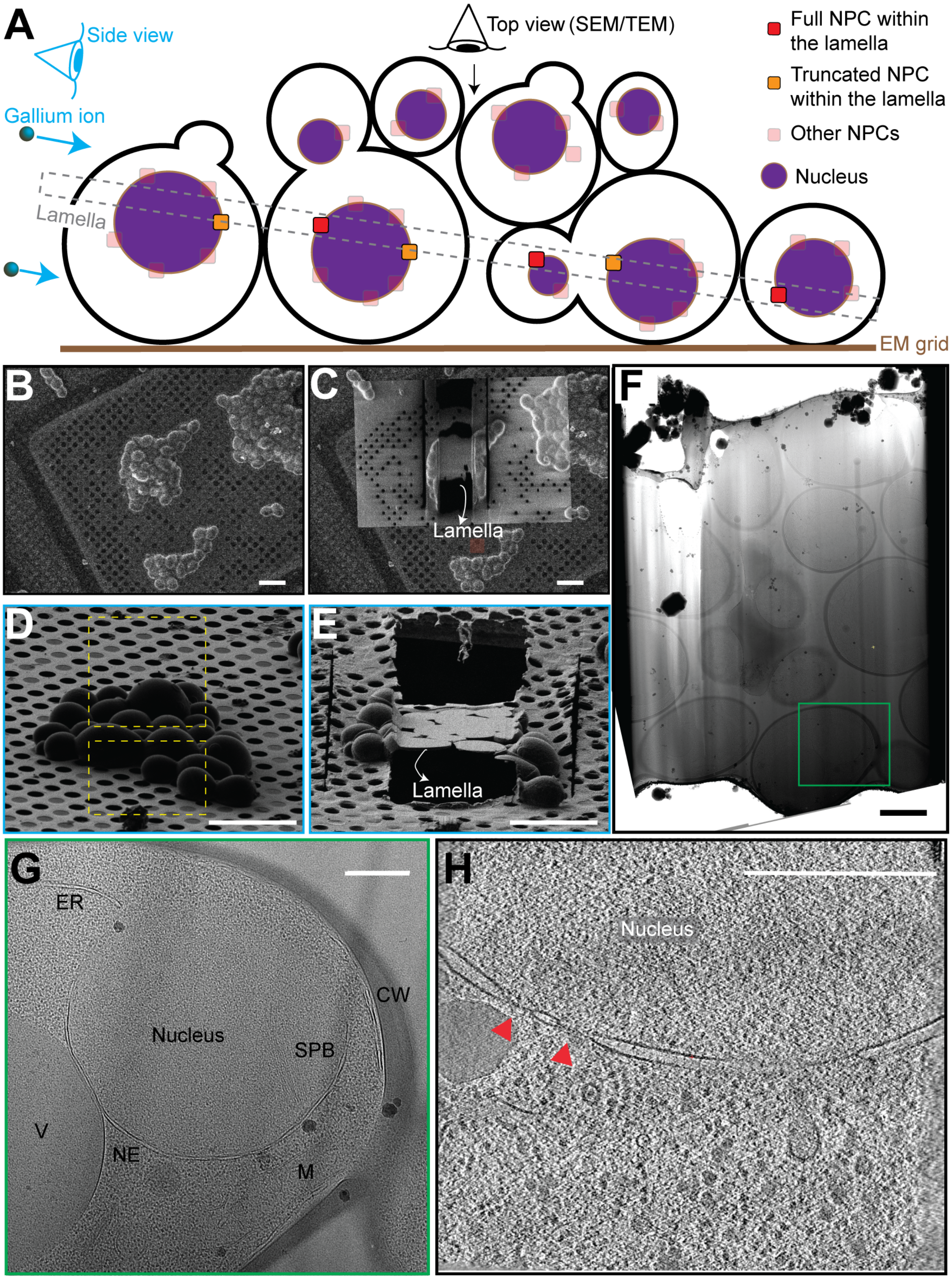


**Fig. S9. Overview of cryo-ET sample preparation for in situ yeast NPCs.** **A**. Schematic of cryo-FIB milling of cryogenically frozen yeast cells. A focused beam of Gallium ions is used to micro-machine a thin lamella from clusters of yeast cells. Probabilistically, a lamella will contain a few whole NPCs (indicated by red rectangles) and a few truncated NPCs (indicated by orange rectangles). In the milling configuration, the FIB can be used to get a milling eyes view of the cluster topography, whereas the scanning electron microscopy (SEM) can be used to get a top view of the cell cluster. **B**. SEM image of multiple clusters of yeast cells. **C**. Post milling, SEM image of the milled center cluster super-imposed on the original image with other clusters. **D**. FIB image of a cluster of the yeast cells before milling. The milling patterns (yellow rectangles) are the area over which the FIB is rastered to sputter away the cell material and create the intervening thin lamella. **E**. FIB image of a cluster of milled yeast cells with the lamella indicated. **F**. Transmission EM montage of the lamella. **G**. Transmission EM micrograph of a region of the lamella showing a section of a vitrified cell. **H**. A 14 nm slice through a tomogram with NPCs indicated by red arrows. NPC: Nuclear pore complex; NE: nuclear envelope; SPB: Spindle pole body; V: Vacuole, CW: Cell-wall; M: Mitochondria; ER: Endoplasmic reticulum. Scale bars: (B-E) 10 µm; (F) 2 µm; (G-H) 500 nm.

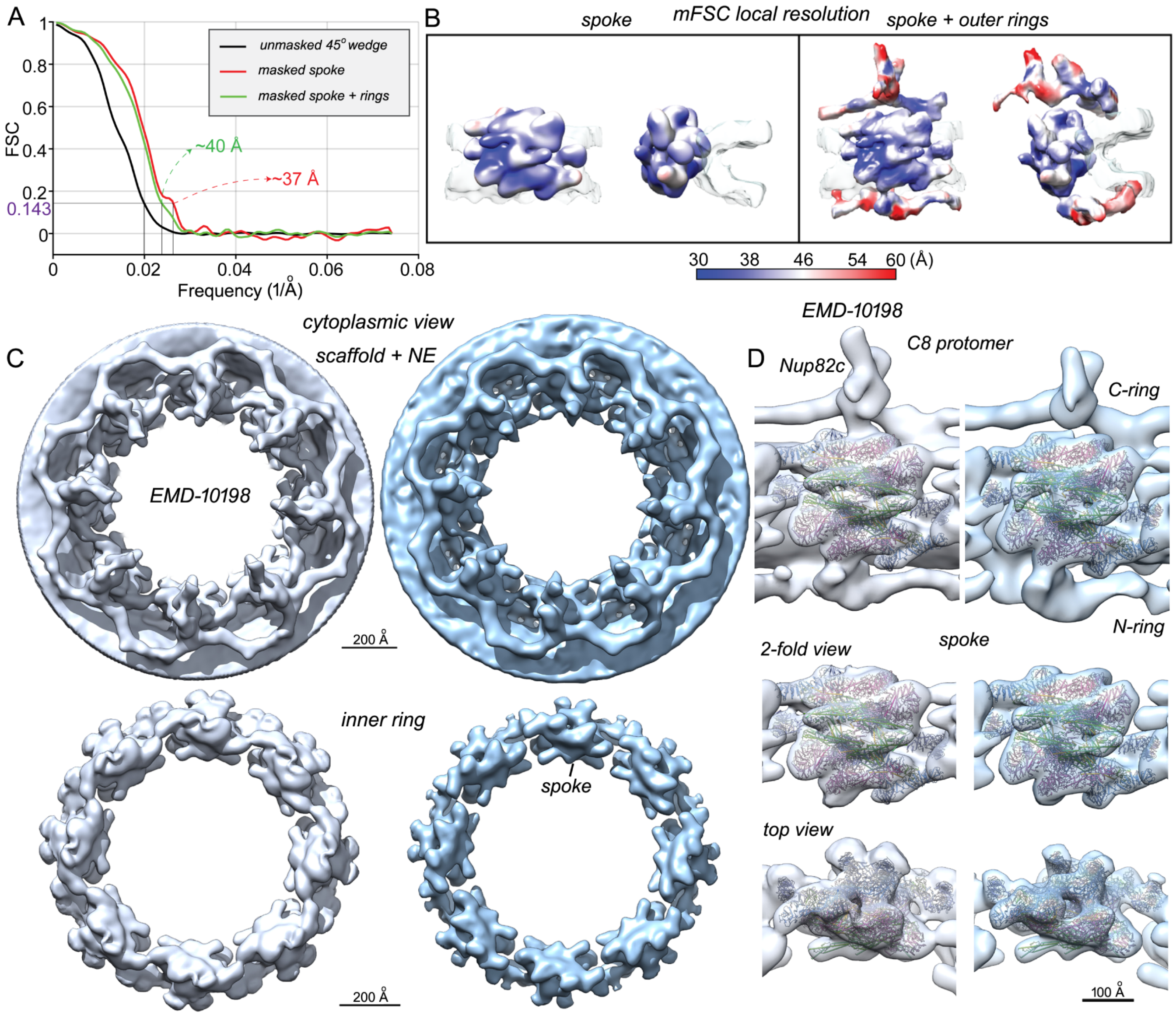


**Fig. S10. Resolution of the core scaffold from the in situ NPC and comparison with a previously determined tomographic structure A**. FSC curves are shown for the spoke (red) and a larger volume containing the spoke and outer rings (green) along with a comparison of unmasked half volumes (black). **B**. Local resolution calculated with mFSC is displayed on the 3D map for the spoke (left) and a full protomer (right). **C**. Side-by-side comparisons for 3D maps are shown for EMD-10198 ((**?** )) in grey and our structure (in blue), as viewed from the cytoplasm. (top) Full 3D maps; (bottom) segmented inner rings. **D**. Each column shows the density map for the C8 protomer and two views of the spoke with docked models for the spoke. (left) EMD-10198, (right) current 3D density map.

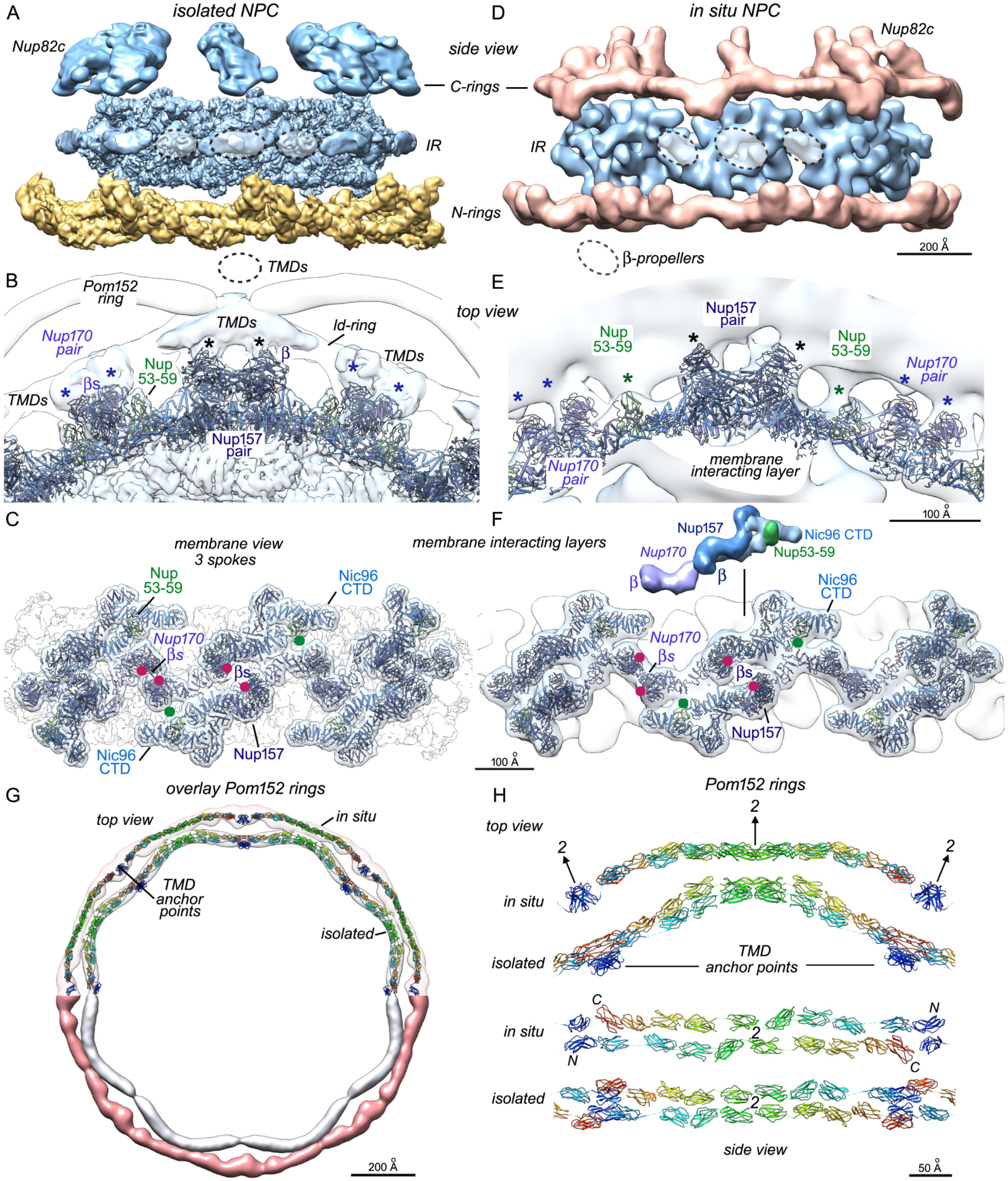


**Fig. S11. Membrane contacts of the inner ring and arrangement of the Pom152 ring A-C**. Membrane interactions of the isolated NPC. A. A side view of the isolated NPC is shown with contact sites to the lipid-detergent ring that lie on local 2-fold axes (dashed ovals). **B**. A view from the cytoplasm of the membrane interacting layer in the inner ring shows contact sites for Nup157 and Nup170 β-propellers (black and blue asterisks) with densities in the ld-ring that may represent TMDs for Pom152/34 and Ndc1, respectively. **C**. A side view of the outer surface shows a nearly-continuous band of membrane interacting layers from spokes of the inner ring. Approximate positions of membrane anchor sites for β-propellers (red dots) and Nup53-59 (green dots) are indicated. **D-F**. Membrane interactions of the in situ NPC. **D**. The dimeric clustering of β-propellers from Nup157 (center dashed oval) and Nup170 (left and right) is highlighted in a side view of the in situ NPC without the pore membrane. **E**. A view from the cytoplasm shows membrane interacting sites for respective Nups marked with asterisks. The density distribution is asymmetric across the spoke 2-fold axis that passes between a Nup157 pair. **F**. A side view is shown for the dilated inner ring with approximate positions of membrane anchor sites for β-propellers (red dots) and Nup53-59 (green dots). An icon view shows the relationship of the Nic96 CTD and Nup53-59 with extended Nup157 and Nup170 molecules in the top half of the membrane interacting layer. The inner ring makes stabilizing contacts with the pore membrane at three different levels. **G**. Pom152 rings from the two structures are overlaid to show changes in shape and diameter that occur during radial expansion. (top) Rainbow colored models (N to C, blue to red) are shown as "pipes and planks" for isolated and in situ Pom152 rings in transparent density maps. Spoke anchor points are indicated for the two rings. (bottom) Isosurface representations are shown for density maps of the Pom152 rings. **H**. A 45 degree wedge from the aligned Pom152 rings is shown with their respective models. (top) The two Pom152 rings are viewed from the cytoplasm with ribbons in rainbow colors; 2-fold axes and TMD anchors are labeled. (bottom) A side view of the Pom152 rings is shown from the NE lumen, in which two anti-parallel strands of Ig-like domains form a flat ribbon.

**Table S1. Single particle data collection, processing and modeling for the yeast NPC.**
*Related to Figures 1, 4.*

| | | |
|---|---|---|
| **Data collection** | | |
| Microscope | Titan-Krios-GIF | |
| Detector | K2 Summit | |
| Image mode | Super-resolution counting<br>40 frames | |
| Total electron dose ($e^-/Å^2$) | 40.0 | |
| Nominal pixel size (Å) | 2.66 | |
| Defocus range (µm) | -1.5 to -3.8 | |
| Movies (after triage)/ frames | 4015 (3218)/40 | |
| NPCs after cleaning | 26049 | |
| **Multi-body 3D reconstruction** | **Spoke** | **Double thin ring** |
| Symmetry imposed | C1 | C1 |
| Software | RELION 2.1, 3.0 | RELION 2.1, 3.0 |
| Initial subunits/final particles | 208393/ 145000 | 208393/ 45000 |
| Pixel size (Å) | 2.66 | 3.99 |
| Map resolution (Å)-masked<br>FSC threshold 0.143 | 7.6 | 11.3 |
| Local resolution range (Å) | 6.6-11.0 | 11-17 |
| B-factor for sharpening ($Å^2$) | -423 | -800 (ad hoc) |
| **Modeling** | | |
| Software- | Chimera, Coot, Phenix, MDFF | Chimera, Coot, Phenix, MDFF |
| Ordered mass / C8 protomer (MDa) | 1.69 | 1.11 (~2.8 MDa) |
| Number of proteins, excluding Pom152 | 28 | 15 + 1 unknown |
| Chimera map cross-correlation | 0.915 | 0.952 |

**Fig. T1. Table S1**

**Table S2.** Summary of integrative threading and modeling spoke orphan structural elements and the Pom152 ring

*Related to Figures 3 and 8.*

| | |
|---|---|
| **1) Gathering information** | |
| Prior models | Structure of the isolated andi̇ditu NPCs |
| Physical principles and statistical preferences | Excluded volume<br>Sequence connectivity<br>Predicted secondary structure (RaptorXProperties)<br>Predicted transmembrane domains (http://yeastgenome.org and HeliQuest) |
| Experimental data | 1425 DSS and EDC chemical crosslinks<br>Cryo-EM map; EMDB TBD<br>FIB-milled CET map; EMDB TBD |
| **2) Representing the system** | |
| Resolution of structured components | 1 [R1] residue per bead |
| Resolution of unstructured components | 10 [R10] residues per bead |
| Structural coverage | 71 % |
| Number of structural elements (SEs) | 30 |
| Composition (number of copies of Pom152) | 8 |
| Atomic (structured) components (Pom152) | 260-362, 379-472, 520-611, 616-714, 722-818, 824-918, 931-1026, 1036-1141, 1150-1229, 1244-1337 |
| Unstructured components (Pom152) | 1-259, 361-378, 471-519, 610-615, 713-721, 819-823, 919-930, 1027-1035, 1142-1149, 1230-1243 |
| Spatial restraints encoded into scoring function (Pom152) | Excluded volume; applied to the R1 representation<br>Sequence connectivity; applied to the R1 representation<br>Cross-link restraints; applied to the R1 representation<br>EM density restraint using Gaussian Mixture Model (GMMs) representations<br>Transmembrane domain restraint (Pom152 (111-200))<br>Peri-nuclear restraint (Pom152 (201-1337))<br>Sequence connectivity and excluded volume |
| Spatial restraints encoded into threading scoring function | Secondary structure restraint; applied to the SEs<br>Loop end-to-end disntance restraint; applied to the SEs<br>Cross-link restraints; applied to the SEs and R1 representation |
| **3.1) Enumeration of threading of degrees offreedom** | |
| Sequences used for threading | Nic96 (residues 1-205), Nup53 (residues 1-247), Nup59 (residues 1-265), Nup100 (residues 551-815), Nup116 (residues 751-965), and Nup145N (residues 201-458) |
| **3.2) Structural Sampling** | |
| Sampling method | Replica Exchange Gibbs sampling, based on Metropolis Monte Carlo |
| Replica exchange temperature range | 1.0 - 2.5 |
| Number of replicas | 8 |
| Number of runs | 50 |
| Number of structures generated | 300000 |
| Movers for flexible string of bead | Random translation up to 4.0 Angstroms |
| CPU time | 14 hours on 42 processors |
| **4.1) Validating the threading models** | |
| Protein, sequence range, and start residue standard deviation | SE1: Nic96, 15-30 (1)<br>SE2: Nic96, 38-43 (2)<br>SE3: Nic96, 46-61 (0.3)<br>SE4: Nic96, 66-78 (2)<br>SE5: Nic96, 84-99 (1)<br>SE6: Nic96, 118-141 and 147-166 (1.4)<br>SE7: Nic96, 118-153 and 155-165 (0)<br>SE8: undefined |
| **4.2) Validating the Pom152 ring models** | |
| Models selected for validation | |
| Number of models after equilibration (isolated/in situ) | 300000/300000 |
| Number of models that satisfy the input informaton (isolated/in situ) | 92281/51183 |
| Number of structures in samples A,B (isolated/in situ) | 52831,39450/18722,32461 |

**Table S2** - continued

| | |
|---|---|
| p-value of non-parametric Kolmogorov-Smirnov two sample test (isolated/in situ) | 0.446/0.34 |
| Kolmogorov-Smirnov two-sample test statistic (D (isolated in situ) | 0.0,1E-16 |
| Thoroughness of the structural sampling | |
| Sampling precision (isolated/in situ) | 14.66/12.6 Angstroms |
| Homogeneity of proportions $X^2$ test p-value (Cramers V value) (isolated/in situ) | 1.000 (0.000)/ 1.000 (0.000) (thresholds: p - value > 0.05 OR Cramer's V < 0.1) |
| Number of clusters (isolated/in situ) | 1/1 |
| Cluster populations (isolated/in situ) | Cluster 1: 99/98 % |
| Cluster precisions (isolated/in situ) | Cluster 1: 15.42/10.1 Angstroms |
| Average cross-correlation between localization probability densities of samples A and B (isolated/in situ) | Cluster 1: 0.84/0.92 |
| Validation by information used for modeling | |
| Percent of sequence connectivity restraints satisfied per structure (isolated/in situ) | 99/99 % |
| Percent cross-link restraints satisfied by ensemble (isolated in situ) | 89/89 % |
| Percent of transmembrane domain restraints satisfied by ensemble (isolated/in situ) | 98/99 % |
| Percent of excluded volume restraints satisfied per structure (isolated/in situ) | 99/99 % |
| 6) Software and data availability | |
| Software | |
| Modeling programs | IMP PMI module, version develop-548de65454<br>Integrative Modeling Platform (IMP), version develop-548de65454 |
| Modeling scripts | https://github.com/integrativemodeling/NPC v3.0 |
| Homology detection and structure prediction | HHPred, version 2.0.16 |
| Visualization and plotting | UCSF Chimera<br>Matplotlib, version 3.0.3 |
| Data | |
| PDB-dev accesion code | TBD |

**Fig. T2. Table S2**

**Table S3.** ***In-situ*** **tomographic data collection, processing and modeling for the yeast NPC.** *Related to Figures 5-8.*

| | | | | |
|---|---|---|---|---|
| **Data collection** | | | | |
| Microscope | Titan-Krios-GIF | | | |
| Detector | K2 Summit | | | |
| Energy filter slit | 20 KeV | | | |
| Image mode | Counting | | | |
| Tilt-series scheme | Dose-symmetric & bi-directional | | | |
| Frames per tilt image | 12 | | | |
| Total electron dose ($e^-/Å^2$) | ~100-140 | | | |
| Nominal pixel size (Å) | 3.43 | | | |
| Defocus range: nominal/ actual (μm) | -3 to -5 (-3 to -11) | | | |
| Tomograms | 293 | | | |
| NPCs | 518 | | | |
| **3D reconstruction** | **per-spoke refinement** | | | |
| Symmetry imposed | C1 | | | |
| Software | EMAN2 | | | |
| Particles: total/ used | 3656/ 3290 | | | |
| Adjusted pixel size (Å) | 3.37 (bin2- 6.74) | | | |
| Map resolution (Å)-masked | 37 (spoke) | | | |
| FSC threshold 0.143 | 40 (full C8 protomer) | | | |
| Local resolution range (Å) mFSC | 30-60 | | | |
| **Modeling** | | | | |
| Software- | Chimera, Coot, MDFF | | | |
| NPC component | Spoke | N-ring | C-ring + Nup82c-Dyn2 | |
| Nucleoporins/ proteins, excluding Pom152 | 28 | 7 | 19 | |
| Fitted mass/ C8 protomer (MDa) | 1.69 | 0.49 | 0.75 | (~2.9 MDa) |
| Cross-correlation to final map | 0.924 | 0.875 | 0.882 | |

**Fig. T3. Table S3**

**Table S4. Nucleoporin FG repeats in the central transport path of the yeast NPC.** *Related to Figure 7.*

| Nucleoporins/NPC from C- to N-side (number of copies) | | FG repeat residues | Total amino acids | Mass for FG repeats approximate |
|---|---|---|---|---|
| Nup42 | (8) | 364 | 2912 | $0.28 \times 10^6$ |
| Nsp1 | (16) | 590 | 9440 | $1.00 \times 10^6$ |
| Nup159 | (16) | 876 | 14016 | $1.51 \times 10^6$ |
| Nup116 | (16) | 686 | 10976 | $1.18 \times 10^6$ |
| Nup100 | (16) | 570 | 9120 | $0.98 \times 10^6$ |
| | | | | ~4.9 MDa |
| Nsp1 | (32) | 590 | 18880 | $2.00 \times 10^6$ |
| Nup49 | (32) | 236 | 7522 | $0.81 \times 10^6$ |
| Nup57 | (32) | 220 | 7040 | $0.76 \times 10^6$ |
| | | | | ~3.6 MDa |
| Nup145N | (16) | 209 | 3344 | $0.36 \times 10^6$ |
| Nup1 | (8) | 695 | 5560 | $0.6 \times 10^6$ |
| Nup60 | (8) | ~100 | 800 | $0.09 \times 10^6$ |
| | | | | ~1.1 MDa |
| | | | | Sum ~9.6 MDa |

**Fig. T4. Table S3**
